# Changes in the Dopaminergic circuitry and Adult Neurogenesis linked to Reinforcement Learning in Corvids

**DOI:** 10.1101/2023.08.31.555829

**Authors:** Pooja Parishar, Madhumita Rajagopalan, Soumya Iyengar

**Affiliations:** National Brain Research Centre, Manesar, Nainwal Mode, NH-8, Gurugram, Haryana – 122052, India

**Keywords:** Adult Neurogenesis, Corvids, Dopamine, Neurite pruning, Reinforcement Learning, Visual discrimination

## Abstract

The caudolateral nidopallium (NCL, an analogue of the prefrontal cortex) is known to be involved in learning, memory, and discrimination in crows, whereas the involvement of other brain regions in these phenomena are unknown. However, recent studies on pigeons have demonstrated that besides NCL, basal ganglia-thalamocortical loops connected to this region play are also crucial for learning. The present study demonstrates that besides NCL, other parts of the caudal nidopallium (NC), avian basal ganglia, and intriguingly, vocal control regions in house crows (*Corvus splendens*), are involved in visual discrimination. We have also found that training on the visual discrimination task can be correlated to neurite pruning in mature dopaminoceptive neurons and immature doublecortin-positive neurons in the NC of house crows. Furthermore, there is an increase in the incorporation of new neurons throughout NC and the medial striatum which can also be linked to learning. For the first time, our results demonstrate that a combination of structural changes in mature and immature neurons and adult neurogenesis are linked to learning in corvids.

## 1. Introduction

Both classical conditioning and operant learning depend on topographically organized fronto- striatal pathways composed of thalamo-cortical-basal ganglia loops (1–3), which receive dopaminergic input. In avian brains, the caudolateral nidopallium is functionally analogous to the mammalian prefrontal cortex (PFC) (4,5). Besides NCL, the medial striatum is reported to be involved in task acquisition (6) and multicomponent behavior (7) in pigeons. Furthermore, an anterior nidopallial nucleus (NIML, nidopallium intermedium medialis pars laterale) is known to be involved in sequential learning and serial processing in pigeons (8,9). Its songbird equivalent, LMAN (lateral magnocellular nucleus of the anterior nidopallium), is a part of the anterior forebrain-basal ganglia loop involved in song learning (10). We were interested in studying whether similar circuits underlie operant learning in corvids, known for their cognitive abilities, including physical and social cognition, facial recognition and discrimination (11–14). For these experiments, we used the expression of the immediate early gene Arc (activity-regulated cytoskeletal protein) after house crows (*Corvus splendens*) were trained on a shape discrimination task (15) involving reinforcement learning. The expression of Arc is increased by high-frequency stimulation (16,17), spatial learning (18), and housing in enriched environments (19) in various cortical areas and in the rodent hippocampus and was therefore localized to identify brain areas in crows activated by learning and decision-making.

Furthermore, we wanted to understand whether changes in neural activity associated with learning led to changes in the dopaminergic system in house crows. The dopaminergic system plays a key role in the motivation to learn, reward prediction, and the subjective valuation of reward (20). For example, infusions of a dopamine D1 receptor antagonist in the pigeon NCL led to a decrease in their performance on a novel attention-based task (21). Dopamine release and an increased expression of D1 receptors were also observed in the pigeon NCL linked to working-memory based tasks (22,23). We, therefore, quantified changes in DARPP-32 (Dopamine- and cAMP-regulated phosphoprotein, Mr 32kDa)-positive dopaminoceptive neurons in different parts of the house crow NC after training them on visual discrimination.

Different strategies adopted by the brain for learning include changes in the number of synapses and/or dendritic remodeling in existing neural circuits to facilitate the consolidation of new information (24) and the addition of new neurons to already existing circuits. For example, shifting rats to an enriched environment (25), training them on a T-maze task or spatial reversal learning in a parallel alley maze led to layer-specific changes in the dendritic field of pyramidal neurons in the medial prefrontal cortex (mPFC, CG3) and in the orbitofrontal cortex (OFC).

These findings suggest that the plasticity induced by different kinds of experience varies in different cortical areas and layers (24). Learning is also known to induce changes in adult neurogenesis (26). In seasonal songbirds such as male canaries which change their songs annually during the breeding season, projections from the pallial region HVC to the motor nucleus RA (Robust nucleus of the arcopallium) are remodeled during adulthood as a result of neurogenesis. Since these neurons are lost after this period, they may be necessary or permissive for learning or producing new songs (27). New neurons are also incorporated in the caudomedial nidopallium (NCM), an auditory area important for perception and storage of conspecific songs in zebra finches (28), in the NCL and hippocampus of adult house crows (29) and in the striatum of humans (30), rodents (31–33) and songbirds (zebra finches) (34), where they mature into medium spiny neurons (MSNs). Although the striatum has been widely studied for its role in goal-oriented learning and decision-making (35), changes in adult neurogenesis induced by these processes in this region have not been reported so far.

We were interested in understanding whether learning and decision-making induced structural changes in mature dopaminoceptive (DARPP-32) and immature Doublecortin-labeled (DCX) neurons (29) in the house crow NC. Furthermore, we decided to study adult neurogenesis in the striatum and NC using DCX as a marker for immature neurons, since these areas were involved in visual discrimination and are known to recruit new neurons during adulthood (34).

## 2. Results

### 2.1. Quantification of motor activity related to learning

We quantified the total time taken by crows to retrieve the food reward, which included sequences of motor activity wherein the crow hopped down from the perch, walked towards the shape and overturned it, after which it retrieved the food reward. The amount of time taken for individual crows to obtain rewards provides an indirect measure to quantify motor activity of the crows. Due to the greater number of trials associated with learning the task, the cumulative activity period was higher in Trained birds (Kruskal-Wallis test; χ2 = 8.0952, df = 2, P = 0.01746) compared to that of Undertrained (Dunn’s post hoc test; P= 0.0271) and No-Association (P= 0.0337) birds [**Supplementary Figure** 1(a)]. We did not find any differences in motor activity between No-Association and Undertrained birds since both groups received approximately two training sessions. However, we could not compare motor activity of the Baseline group with that of other groups, since for this set of birds, the experimenter left the room and did not perform further trials as for other groups. Moreover, the purpose of the Baseline group was to serve as a control for visual input induced by the behavioral apparatus without learning the visual discrimination task.

### 2.2. Activation of house crow brain regions associated with visual discrimination

Besides NC [**Figure 1(a)**], areas including LMAN, Area X, RA, and AId, generally associated with song control (10) expressed the Arc protein in different groups of house crows [**Supplementary Figure** 2(a)**, (b), and (e)**]. Intensely stained neurons were also present in components of the basal ganglia including LSt, GP [**Supplementary Figure** 2(c) and 3(a)] and MSt, and the midbrain dopaminergic areas, VTA and SN [**Supplementary Figure** 3(c); see **Supplementary Figure** 2(d) for a negative control devoid of label at the level of GP and LSt)].

**Figure 1.**
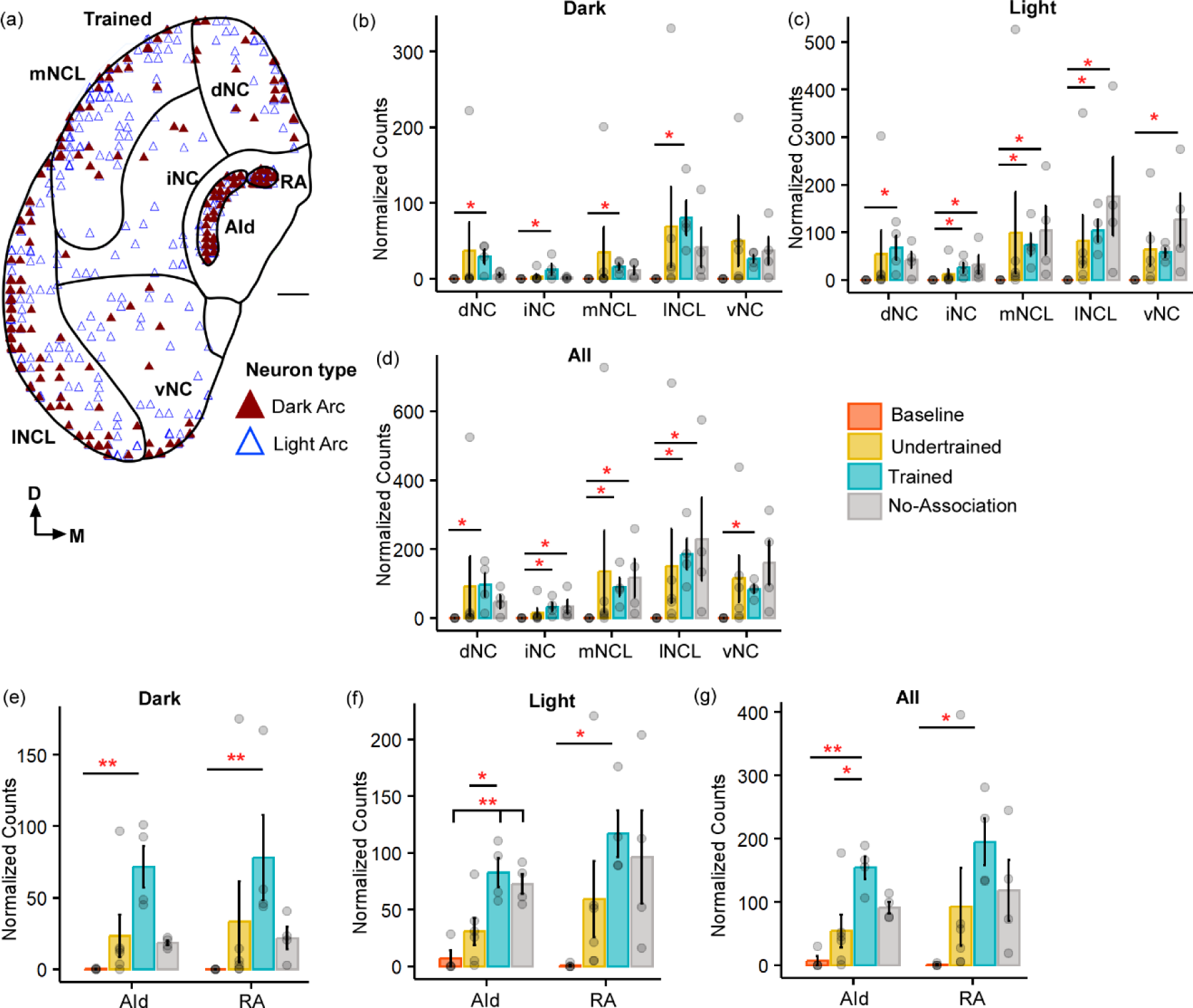
The expression of Arc in the caudal nidopallium (NC) and arcopallium across different sets of experimental birds. (**a**) A coronal schematic of the posterior part of the house crow telencephalon demonstrating label for Arc in the caudal nidopallium and areas of the arcopallium, that is, AId and RA, from one of the Trained birds. Scale bar, 1mm. A comparison of (**b**) darkly stained, (**c**) lightly stained and (**d**) the total Arc-positive neuronal population across different subdivisions of the caudal nidopallium revealed that the highest levels of activity were observed in lNCL (mean ± SEM). The number of dark Arc-positive neurons was the highest in the Trained category followed by that in the No-Association group and the Baseline group had negligible numbers of Arc-labeled neurons. Levels of neural activity were highly variable in the Undertrained group in all divisions of the caudal nidopallium. The number of lightly stained and total Arc cells were also the highest in the Trained and No-Association groups. A comparison of (**e**) darkly stained, (**f**) lightly stained and (**g**) the total Arc population across different behavioral groups in AId and RA. The highest number of Arc-labeled neurons were present in AId and RA of the Trained group, followed by that in the No-Association and Undertrained group. The lowest number of Arc-positive neurons were present in the Baseline group in both AId and RA. * P < 0.05; ** P < 0.01.

Details of the statistical analyses for all data are provided in the **Supplementary Tables.**

#### 2.2.1. Caudal Telencephalon

All five subdivisions of the caudal nidopallium (dNC, iNC, lNCL, mNCL, and vNC) in house crows (36) [**Figure 1(a)**] contained Arc-positive neurons following performance on the visual discrimination task. There were significantly fewer Arc-positive neurons in Baseline controls versus those in Trained and No-Association birds throughout NC [**Figure 1(b-d);** P<0.05; **Table S1**]. We also observed that the number of Arc-labeled neurons was highly variable across NC in Undertrained house crows [**Figure 1(b-d)**].

Besides activation in NC, there were significantly more Arc-positive neurons in the arcopallial region AId of Trained birds compared to those in Baseline controls (P<0.01) as well as in Trained versus Undertrained house crows [**Figure 1(e-g)**; P<0.05; **Table S1**]. Furthermore, there were significantly more Arc-positive neurons in RA of Trained versus Baseline birds [**Figure 1(e-g)**; P<0.05 and P<0.01; **Table S1**], which plays an important role in vocalization and breathing (37).

#### 2.2.2. Regions adjacent to the Anterior Commissure

There were significant differences in the number of Arc-labeled neurons at the level of GP and LSt in Undertrained, Trained, and No-Association birds versus those in Baseline controls [**Supplementary Figure** 3**(a-b)**; P<0.05; **Table S1**]. Similar results were also obtained for the midbrain regions VTA and SN [**Supplementary Figure** 3**(c-d);** P<0.05; **Table S1**].

#### 2.2.3. Anterior Forebrain

There was an increase in neural activity in LMAN of No-Association birds compared to that in Baseline, Trained and Undertrained groups. However, a post-hoc analysis revealed that these differences were statistically significant only for comparisons of Arc-positive neuronal counts between Baseline and No-Association birds [**Figure 2(a-f);** P<0.05 and P<0.01; **Table S1**].

**Figure 2.**
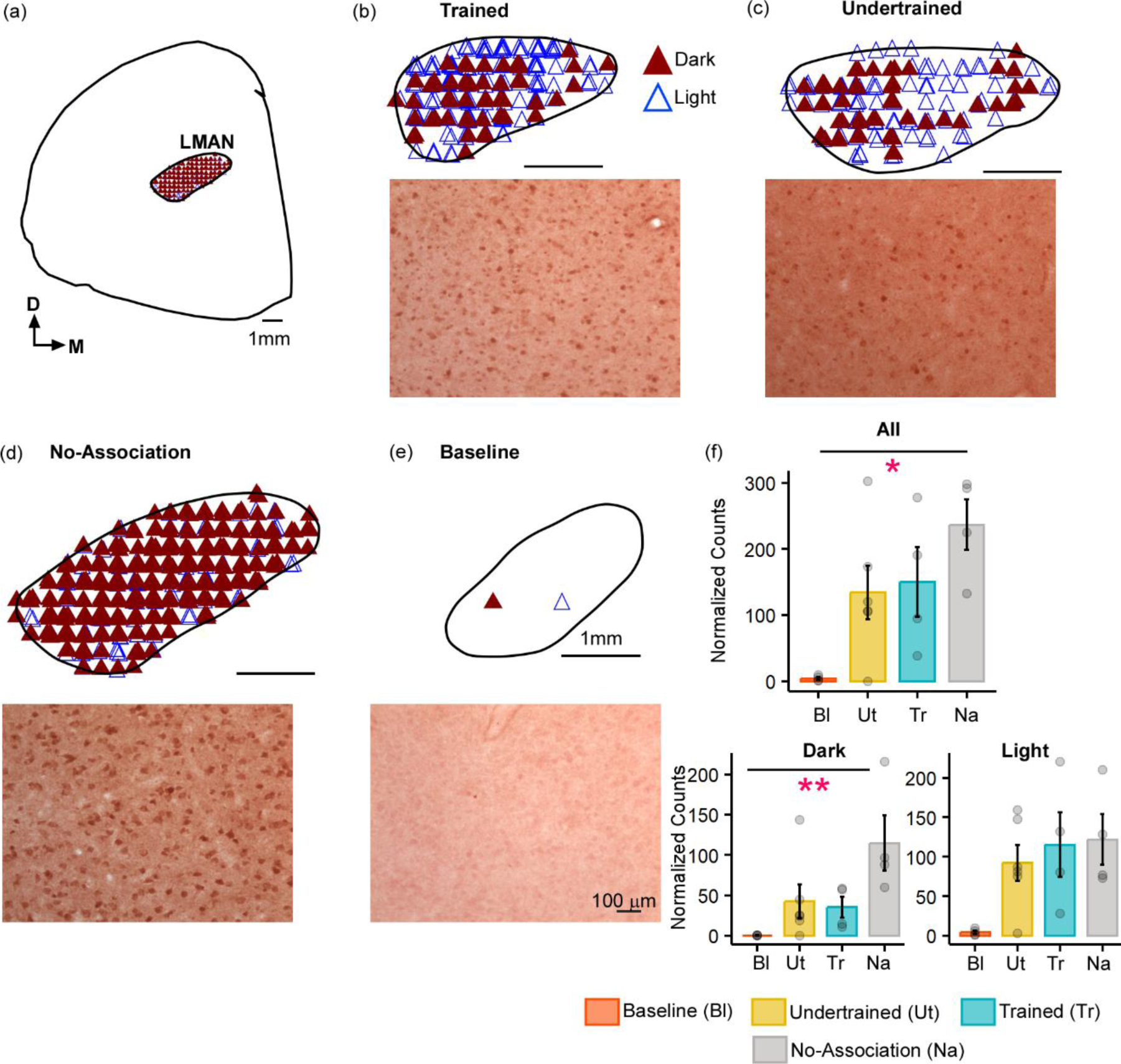
Task-related changes in Arc expression in LMAN. (a) A schematic of the anterior forebrain at the level of LMAN showing Arc expression from a house crow from the No-Association group. Counts of Arc-positive neurons are shown in the schematics at the top and high magnification images of LMAN are shown below in each panel for (b) Trained, (**c**) Undertrained, (**d**) No-Association and (**e**) Baseline groups of experimental birds. The highest expression of Arc was present in the No-Association group (**d**), whereas there was minimal expression in the (**e**) Baseline group. (**f**) Comparisons of normalized total Arc counts (mean ± SEM) in LMAN demonstrated that levels of neural activity were significantly higher in the No-Association group versus the Baseline group. This was true for both the total Arc population and darkly stained populations of neurons (shown in the lower panel). Scale bar, 1mm for the section and LMAN schematic and 100µm for Arc-labeled magnified images. * P < 0.05; ** P < 0.01.

In the striatum at this level, Arc-labeled neurons were observed in Area X and the surrounding MSt in both large parvalbumin-positive (interneurons/pallidal neurons) or smaller parvalbumin-negative neurons [**Supplementary Figure** 4**(a-e)**]. A Two-way ANOVA on Ranks comparing the effect of area and experimental condition on neural activity demonstrated that these parameters interacted only for the total Arc-labeled neuronal population (P=0.048). When the number of Arc-positive neurons of different staining intensities and sizes in Area X and MSt were compared across different experimental groups, the largest number of these neurons was observed in Trained and No-Association birds, whereas the smallest number were present in Baseline controls [**Figure 3(a-e)**; P<0.05 and P<0.01; **Table S1**]. We also found that overall, there were more Arc-labeled neurons in MSt than in Area X following the visual discrimination task [**Figure 3(e)**; P<0.01; **Table S1**].

**Figure 3.**
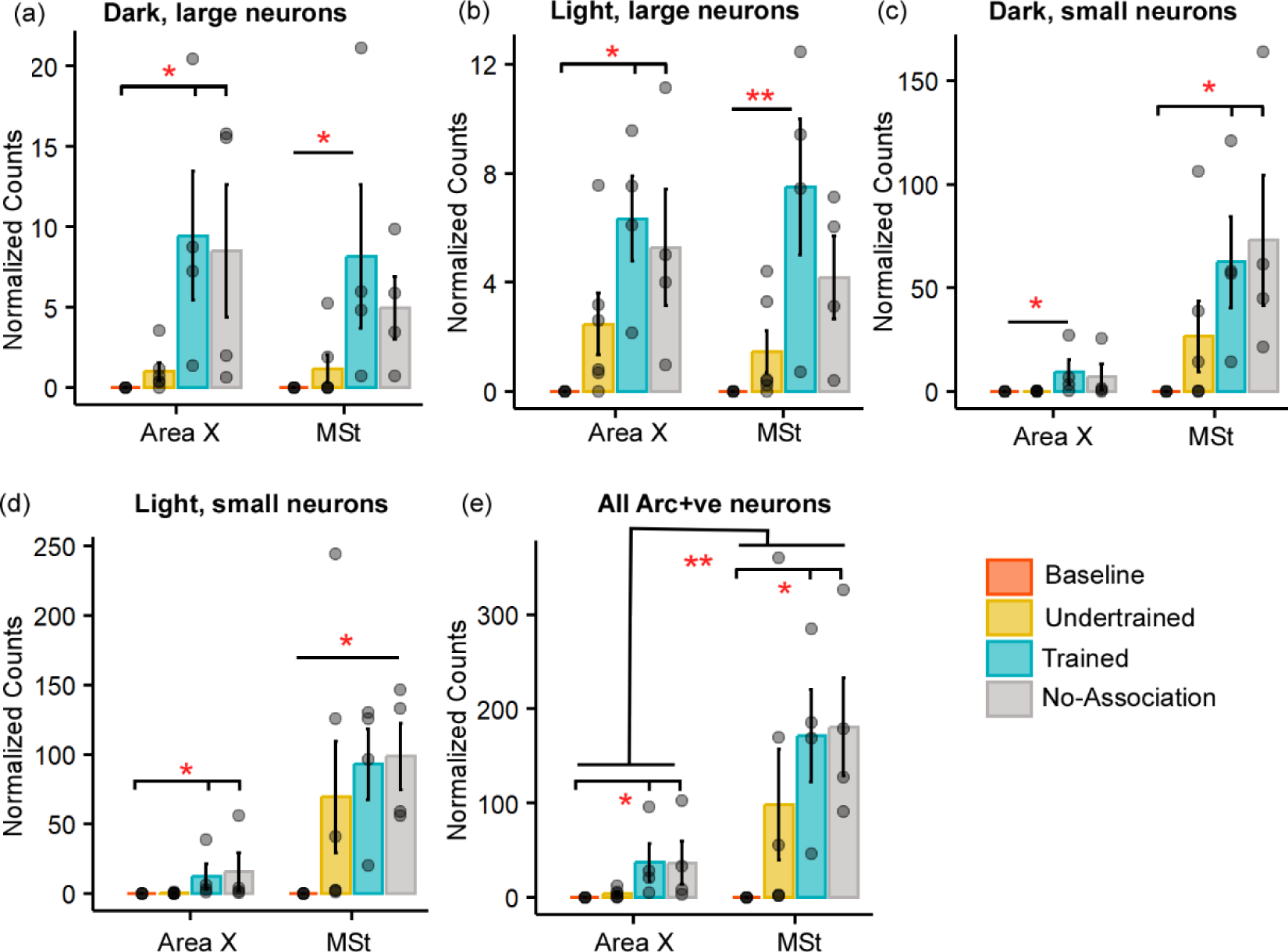
Quantification of Arc-positive neurons in Area X and MSt following the visual discrimination task. Bar graphs representing normalized counts (mean ± SEM) for large neurons which are (**a**) darkly (intensely) stained and (**b**) lightly stained, (**c**) darkly stained small neurons, (**d**) lightly stained small neurons, and (**e**) total Arc-positive neuronal population in Area X and MSt across various behavioral groups. The Baseline group has the lowest number of Arc-positive neurons in both Area X and MSt regardless of the cell type or staining intensity, followed by the Undertrained group, whereas there were a large number of Arc-labeled neurons in the Trained and No-Association groups. * P<0.05; ** P<0.01.

### 2.3. Circuitry involved in Motivation and Reward

#### 2.3.1. Tyrosine Hydroxylase (TH-positive) fibers

Different subdivisions of NC were demarcated in all house crows, based on TH expression (a marker for catecholaminergic synthesis) as previously described (38) [**Supplementary Figure** 5(a)]. The only difference across various groups of birds was a greater density of TH-positive fibers in Baseline birds versus that in Undertrained birds in lNCL [**Supplementary Figure** 5(b); ANOVA, P = 0.0126, F_(3,14)_ = 14.271; Tukey’s post hoc test, P = 0.0084].

#### 2.3.2. Dopaminoceptive (DARPP-32-positive) neurons in NC and the medial striatum

We quantified dopaminoceptive neurons in MSt, Area X [**Supplementary Figure** 5(c)] and NC of all experimental birds [**Supplementary Figure** 5(e)]. The boundaries of Area X were decided based on staining patterns of DARPP-32 and Nissl. Whereas the density of DARPP-32 neurons was lower in Area X compared to that in MSt [Two-way ANOVA; F_(1,28)_ = 9.225, P = 0.0051; **Supplementary Figure** 5(d)], there were no significant differences in the number of dopaminoceptive neurons in the striatum or NC [**Supplementary Figure** 5(f)] in various experimental groups.

##### 2.3.2.1. Learning-induced changes in Dopaminoceptive neurons

There was a statistically significant interaction between the subdivision of NC and experimental condition for the number of nodes (P=7.2935e-10), neurite length (P<2.22e-16), and neurite surface area (P<2.22e-16). The complexity of DARPP-32-labeled neurons [based on neurite length and nodes; **Supplementary Figure** 6**(a-e); Table S2**] increased in all pallial subdivisions in Trained birds versus those in other groups. The neurite length and number of nodes of DARPP-32-labeled neurons located in mNCL and lNCL of the No-Association group increased significantly compared to those in Undertrained and Baseline birds [**Supplementary Figure** 6**(d-j); Table S2**]. The three-dimensional surface area of DARPP-32-labeled neurons and their projections was significantly greater in Trained, Undertrained and No-Association groups versus that in Baseline controls [**Supplementary Figure** 6(f)**; Table S2**]. Furthermore, amongst these three experimental groups, the neurite field and soma size of DARPP-32-positive neurons was significantly greater in all divisions of NC in Trained birds versus that in other groups.

Sholl analysis revealed that the greatest changes in neurite length and number of intersections in dopaminoceptive neurons occurred between the 20-70µm Sholl radii for all parts of NC. The number of intersections and neurite length increased significantly in dNC and iNC in Trained, No-Association and Undertrained birds versus Baseline controls and in Trained versus Undertrained crows [**Supplementary Figure** 7(a)**, (b), (d) and (e); Table S3**]. However, these parameters were similar in Trained and No-Association birds. Neurite length of DARPP-32- positive neurons was significantly greater in mNCL, lNCL and vNC of Trained crows versus that of other groups [**Supplementary Figure** 6(h)**, (j), and 7(f); Table S3**] whereas the number of intersections was significantly greater only in mNCL and vNC of Trained versus No-Association categories of experimental birds [**Supplementary Figure** 6(g) and 7(c)**; Table S3**]. Overall, learning appeared to lead to an increase in the number and length of neurites of dopaminoceptive neurons in all divisions of the house crow NC.

##### 2.3.2.2. Counts and structural complexity of active DARPP-32-labeled neurons in NC

Neurons double-labeled for Arc and DARPP-32 (active dopaminoceptive neurons) were present in NC in all groups of experimental birds except Baseline controls, which were not analyzed further [**Figure 4(a)**]. Active (Arc-positive) dopaminoceptive neurons were reconstructed only in mNCL and lNCL, since the greatest changes in the number of Arc-positive neurons and in the structural complexity of DARPP-32-labeled neurons occurred in these subdivisions. A One-way ANOVA revealed that the number of active dopaminoceptive neurons was significantly greater in Undertrained versus Trained and No-Association birds only in lNCL [**Supplementary Figure** 8**(a-b);** ANOVA, P = 0.0138; F_(2,11)_; Tukey’s post hoc, Ut vs Na: P = 0.0219; Ut vs Tr: P = 0.0414]. A non-significant increase in this measure was observed in Undertrained birds versus other experimental groups of house crows in all other subdivisions of NC.

**Figure 4.**
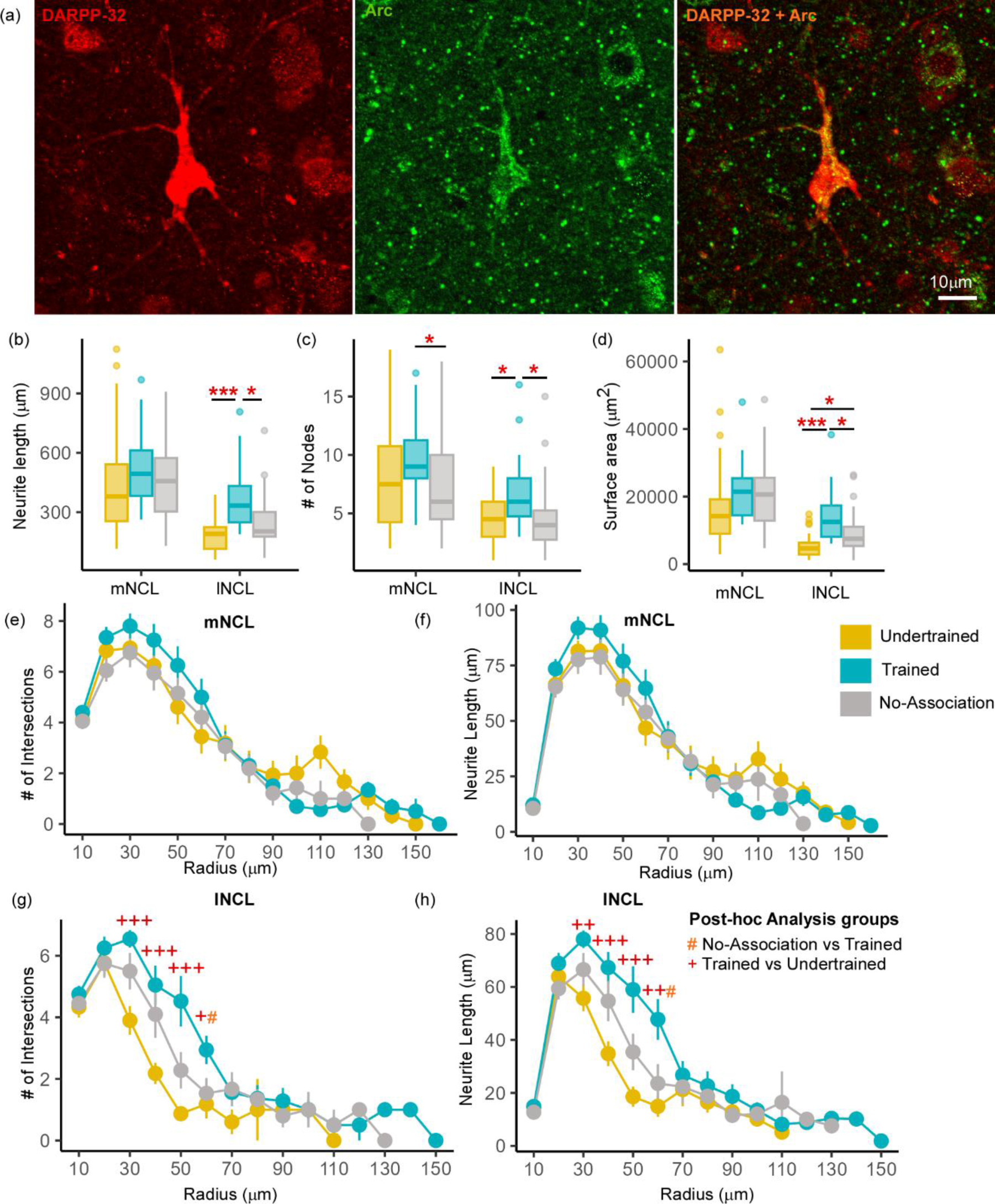
Differences in the structure of neurons double-labeled for Arc and DARPP-32 in NCL. (**a**) High power image of a DARPP-32-labeled neuron (red) from the lNCL region which was also labeled for Arc (green). The third image in this panel shows the co-localization of DARPP-32 and Arc in this neuron. Scale bar: 10µm. Box and whisker plots demonstrate a statistically significant increase in (**b**) neurite length and the (**c**) number of nodes in Trained birds compared to that in the No-Association and Undertrained categories in lNCL and in Trained versus No-Association birds in mNCL. (**d**) An increase in the surface area of neurites was seen in lNCL of Trained birds versus that in Undertrained and No-Association birds. A Sholl analysis demonstrated non-significant increases in the (**e**) number of intersections and (**f**) neurite length (mean ± SEM) in mNCL of the Trained group compared to that in others. A significant difference was observed in (**g**) the number of intersections and (**h**) neurite length in Trained versus Undertrained birds and No-Association versus Trained birds in lNCL. */#/+ P<0.05; ***/###/+++ P<0.001.

##### 2.2.2.3. Morphometric analysis of Active and Inactive Dopaminoceptive neurons

Active dopaminoceptive neurons were observed throughout mNCL and lNCL [**Figure 4(a)**]. A One-Way ANOVA revealed that neurite length of these neurons was significantly greater in lNCL of Trained versus Undertrained and No-Association birds [**Figure 4(b)**; P<0.05; **Table S4**]. The number of nodes increased significantly with training in both mNCL and lNCL [**Figure 4(c)**; P<0.05; **Table S4**]. Furthermore, the three-dimensional surface area of these neurons increased significantly in lNCL in Trained versus Undertrained and No-Association groups and in No-Association versus Undertrained crows [**Figure 4(d)**; P<0.05 and P<0.001; **Table S4**].

Based on Sholl analysis, there were no significant differences in the complexity of active dopaminoceptive neurons in mNCL, except for a non-significant increase in the number of intersections and neurite length in Trained birds versus other groups [**Figure 4(e-f)**; **Table S5**]. Both the number of intersections and neurite length were significantly higher in lNCL in Trained versus Undertrained and No-Association birds between Sholl radii of 30-60µm [P<0.05, P<0.01 and P<0.001; **Figure 4(g-h)**; **Table S5**]. Taken together, our findings suggest that active dopaminoceptive neurons in lNCL increased in complexity as a result of training.

The complexity of inactive (Arc-negative) DARPP-labeled neurons in mNCL and lNCL (based on neurite length, number of nodes, and surface area) was significantly greater in Trained and No-Association birds compared to those in Undertrained birds [**Figure 5(a-c)**; P<0.001; **Table S6**]. Their soma area was also significantly greater in Trained and No-Association birds versus that in Undertrained birds [**Figure 5(d)**; P<0.01 and P<0.001; **Table S6**]. Similarly, the number of intersections and neurite length increased significantly in both regions between Sholl radii 20– 70µm in Trained and No-Association crows versus Undertrained birds [**Figure 5(e-h)**; **Table S7**].

**Figure 5.**
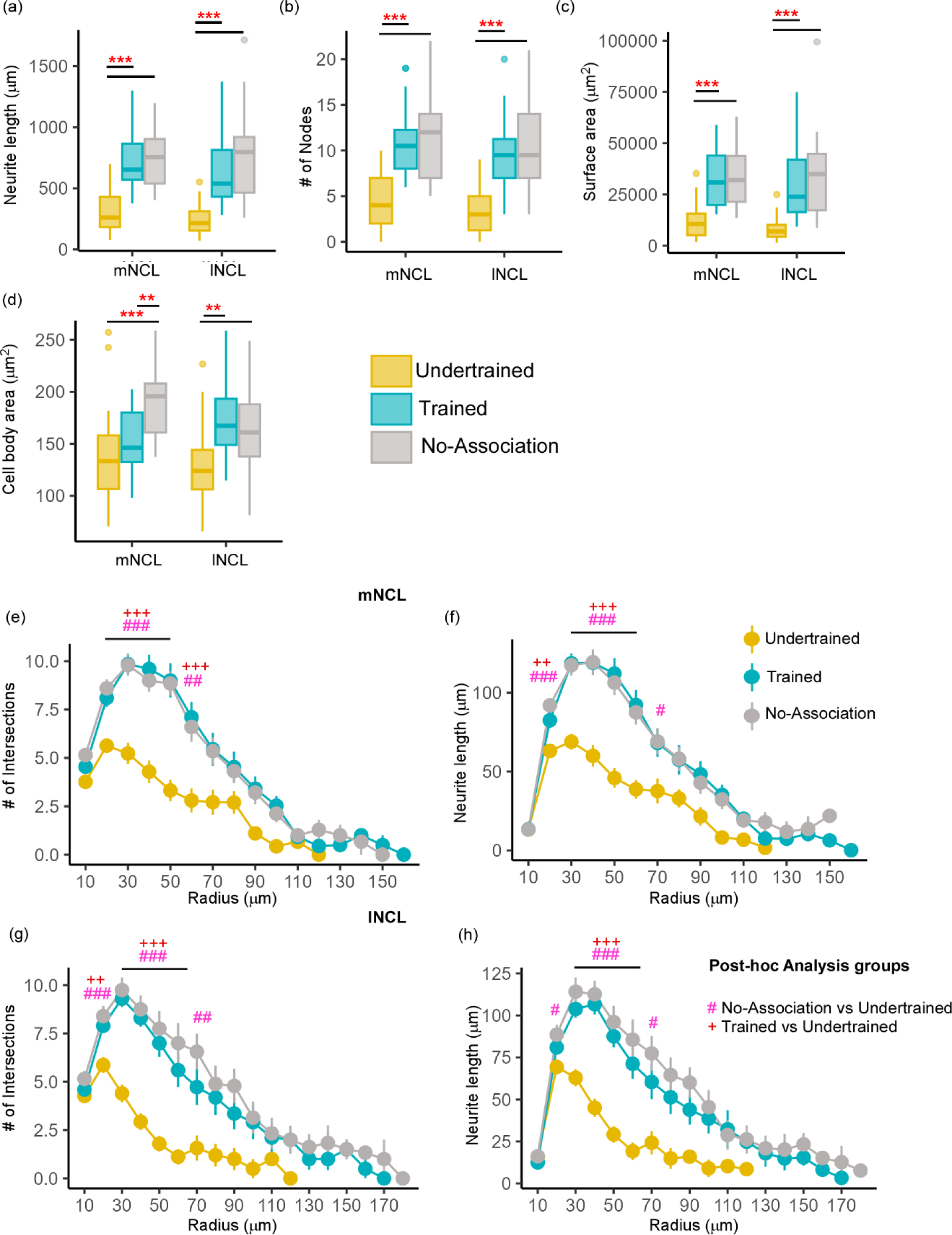
Differences in the morphology of inactive DARPP-32 (Arc-negative) neurons in NCL. Box and whisker plots demonstrating changes in (**a**) neurite length, (**b**) number of nodes, (**c**) neurite field area, and (**d**) cell body area in mNCL and lNCL. In both regions, inactive DARPP- 32 neurons demonstrated greater complexity and cell body expansion in the Trained and No-Association groups compared to that in Undertrained birds. A Sholl analysis in mNCL demonstrated an increase in (**e**) the number of intersections and (**f**) neurite length. Similar results were obtained for the (**g**) number of intersections and (**h**) neurite length in lNCL. */#/+ P<0.05; **/##/++ P<0.01; ***/###/+++ P<0.001.

##### 2.3.2.4. Differences between Active and Inactive Dopaminoceptive neurons

All measures of complexity of neurites for active DARPP-32-positive neurons in lNCL were significantly lower than for inactive dopaminoceptive neurons in Trained and No-Association birds [**Supplementary Figure** 9**(a-d);** P<0.05, P<0.01 and P<0.001; **Table S8**]. However, the soma area of active and inactive dopaminoceptive neurons did not vary across groups [**Supplementary Figure** 9(e); **Table S8**].

As for mNCL, active DARPP-32-labeled neurons were less complex, with lower neurite length, number of branches and neurite surface area compared to inactive ones in No-Association birds [**Supplementary Figure** 9**(f-i);** P<0.01 and P<0.001; **Table S8**]. Whereas active neurons of Trained crows demonstrated similar trends, significant differences were only observed for neurite length and neurite field area. In contrast to lNCL, active dopaminoceptive neurons in mNCL of Undertrained birds demonstrated a significant increase in branching and soma area versus inactive ones [**Supplementary Figure** 9**(f-j);** P<0.05 and P<0.001; **Table S8**].

### 2.4. Changes in adult neurogenesis: Immature (DCX-labeled) neurons in House Crows

Based on their morphology, DCX-positive neurons in house crows could be categorized as (i) spherical neurons which were devoid of processes, (ii) spindle-shaped unipolar or bipolar fusiform neurons, and (iii) multipolar neurons whose somata were rounded (27,29,39) or triangular (40) [**Supplementary Figure** 10**(a-c)**; see **Supplementary Figure** 10(g) for the negative control].

#### 2.4.1. Stereological Counts of DCX-positive neurons

##### 2.4.1.1. Medial and Lateral striatum

The boundaries of Area X could be clearly delineated from the surrounding MSt since it was more myelinated and contained comparatively fewer DCX-positive neurons (**Supplementary Figure** 10(d)). An interaction effect was observed for spherical cells (P<0.05) between the area and experimental condition. Furthermore, a one-way ANOVA/ Kruskal-Wallis Rank Sum test revealed significantly greater numbers of spherical neurons in Area X [P<0.05 and P<0.01; **Figure 6(a)**; **Table S9**] and MSt [P<0.01; **Figure 6(b)**; **Table S9**] of Trained birds versus those in Baseline and Undertrained groups, whereas there were no differences in the number of fusiform, multipolar and total numbers of DCX-labeled neurons in these striatal regions across different experimental groups. Whereas DCX-labeled neurons were present in LSt [**Supplementary Figure** 10(e)], there were no significant differences in their number across different groups of house crows [**Figure 6(c)**]. Our results, therefore, suggest that training on the visual discrimination task may induce an increase in spherical DCX-positive neurons in the medial striatum of Trained birds.

**Figure 6.**
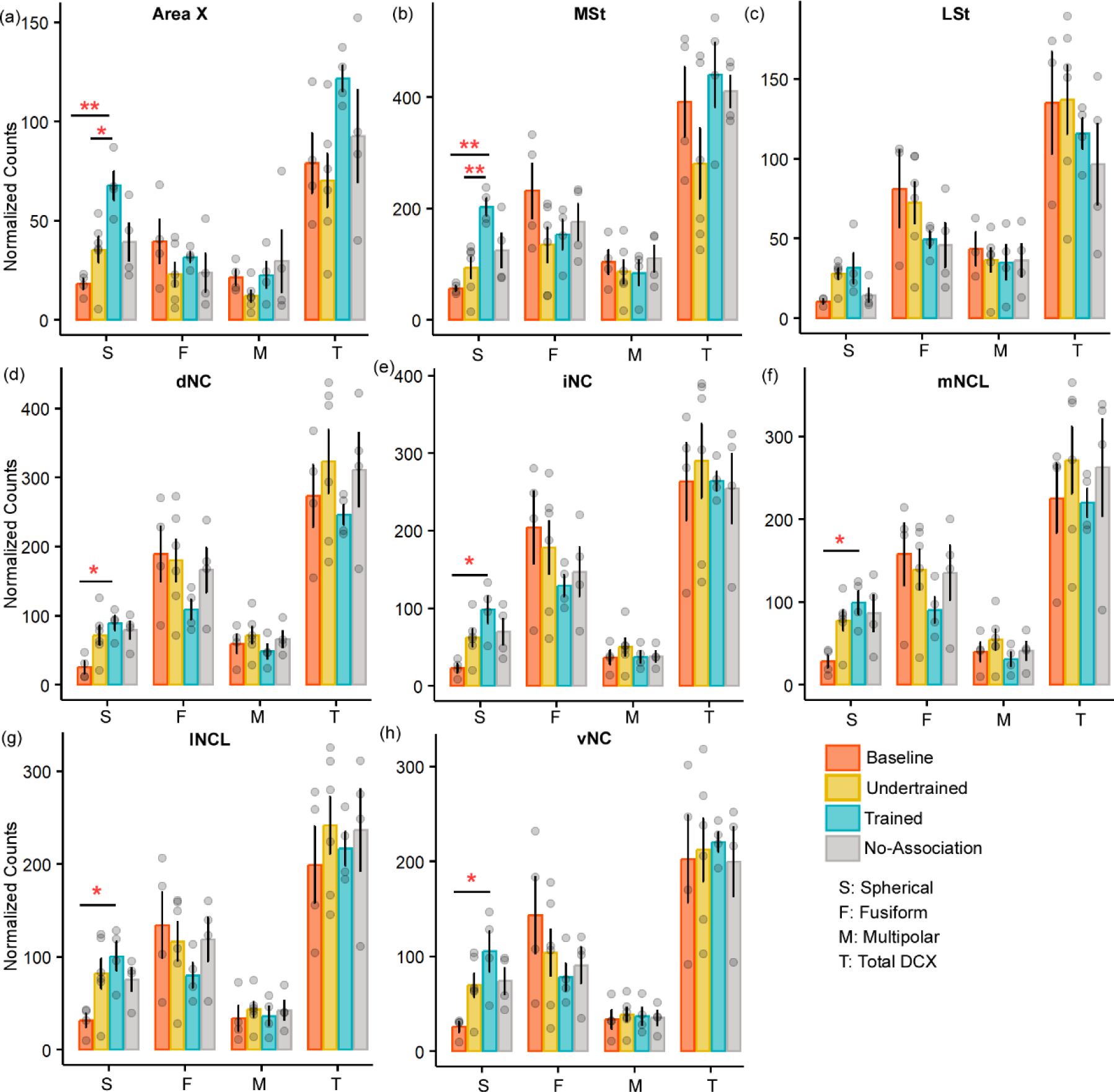
A comparison of the number of various types of DCX neurons in the medial striatum and NC of the four experimental groups of house crows. Bar graphs (mean ± SEM) were plotted to compare the number of DCX-positive neurons across the four experimental groups in (**a**) Area X and (**b**) MSt. The number of spherical DCX-positive neurons was significantly higher in the Trained group versus Baseline and Undertrained groups in both areas, whereas there were no significant differences in (**c**) LSt. The only change observed in all five subdivisions, (**d**) dNC, (**e**) iNC, (**f**) mNCL, (**g**) lNCL, and (**h**) vNC was a significant increase in spherical DCX-positive neurons in the Trained versus Baseline control groups. * P<0.05; ** P<0.01.

##### 2.4.1.2. Caudal Nidopallium

There were no interactions for any DCX-positive cell type in NC across any of the experimental groups of house crows [see **Supplementary Figure** 10(f) for staining patterns]. The only difference was a significant increase in spherical cells in Trained versus Baseline birds in all subdivisions of NC [P<0.05; **Figure 6(d-h)**; **Table S9**].

#### 2.4.2. Morphometric Analysis of Multipolar DCX-positive neurons in MSt

There was no effect of interactions between area and experimental condition for any parameter analyzed. A one-way ANOVA/Kruskal-Wallis test revealed a significant increase in neurite length of DCX-positive multipolar neurons in No-Association and Trained birds compared to Baseline and Undertrained birds in Area X and MSt [**Supplementary Figure** 11**(a-b)**; P<0.001; **Table S10**]. Furthermore, neurite length was significantly greater in DCX-positive neurons of No-Association versus Trained crows in both areas [P<0.05 and P<0.01**; Supplementary** Figure 11(b); **Table S10**]. Neurite branching was greater in DCX-labeled neurons of No-Association birds versus that in Trained, Baseline and Undertrained groups in Area X and MSt [P<0.05 and P<0.001**; Supplementary** Figure 11(c); **Table S10**]. The Dunn’s post-hoc test demonstrated that neurite field area of these neurons in Area X and MSt of No-Association and Trained crows was higher than that in Baseline and Undertrained birds [P<0.001; **Supplementary Figure** 11(d); **Table S10**]. Furthermore, neurite field area of DCX-labeled neurons in No-Association birds was greater than that of Trained birds (P<0.05 and P<0.01) for both areas. We also found an expansion of the neurite field of these neurons in Undertrained versus Baseline birds (P<0.01**; Supplementary** Figure 11(d); **Table S10**). Additionally, the area of DCX-positive neuronal somata was significantly greater in Undertrained, Trained, and No-Association groups versus that in Baseline controls in Area X and MSt [P<0.01 and P<0.001**; Supplementary** Figure 11(e); **Table S10**].

These findings were reflected in Sholl analysis, demonstrating significant differences in the number of intersections and neurite length between Sholl radii 28-68µm in DCX-positive neurons in Area X. Post-hoc tests (Dunn’s or Tukey’s) demonstrated that neurites of DCX-positive neurons in No-Association and Trained birds had significantly more intersections and were longer compared to those of Baseline and Undertrained birds [P<0.05, P<0.01, and P<0.001; **Supplementary Figure** 11**(f-g)**; **Table S11**]. Both parameters were found to be even greater for neurites of DCX-labeled neurons in No-Association versus Trained birds (P<0.05 and P<0.01).

Furthermore, there were a greater number of intersections and an increase in neurite length between Sholl radii 38-58µm in neurons of Undertrained versus Baseline birds [P<0.05 and P<0.01; **Supplementary Figure** 11**(f-g)**; **Table S11**]. Similar differences were observed in MSt for all groups between Sholl radii 18-68µm except for No-Association versus Trained birds [P<0.05, P<0.01 and P<0.001; **Supplementary Figure** 11**(h-i)**; **Table S11**].

##### 2.4.2.1. Morphometric analysis of Active and Inactive DCX-positive neurons

Whereas fusiform DCX-positive neurons were not double-labeled for Arc [**Supplementary Figure** 12(a)], spherical or multipolar double-labeled neurons were very sparsely distributed in the house crow brain [**Figure 7(a)**; **Supplementary Figure** 12**(a-b)**]. As for DARPP-32-labeled neurons, active (Arc- and DCX-labeled) and inactive (only DCX-positive) multipolar neurons in MSt, mNCL and lNCL were reconstructed for further analysis. Since none of the DCX-positive neurons in Area X were double-labeled, this area was excluded from further analysis.

**Figure 7.**
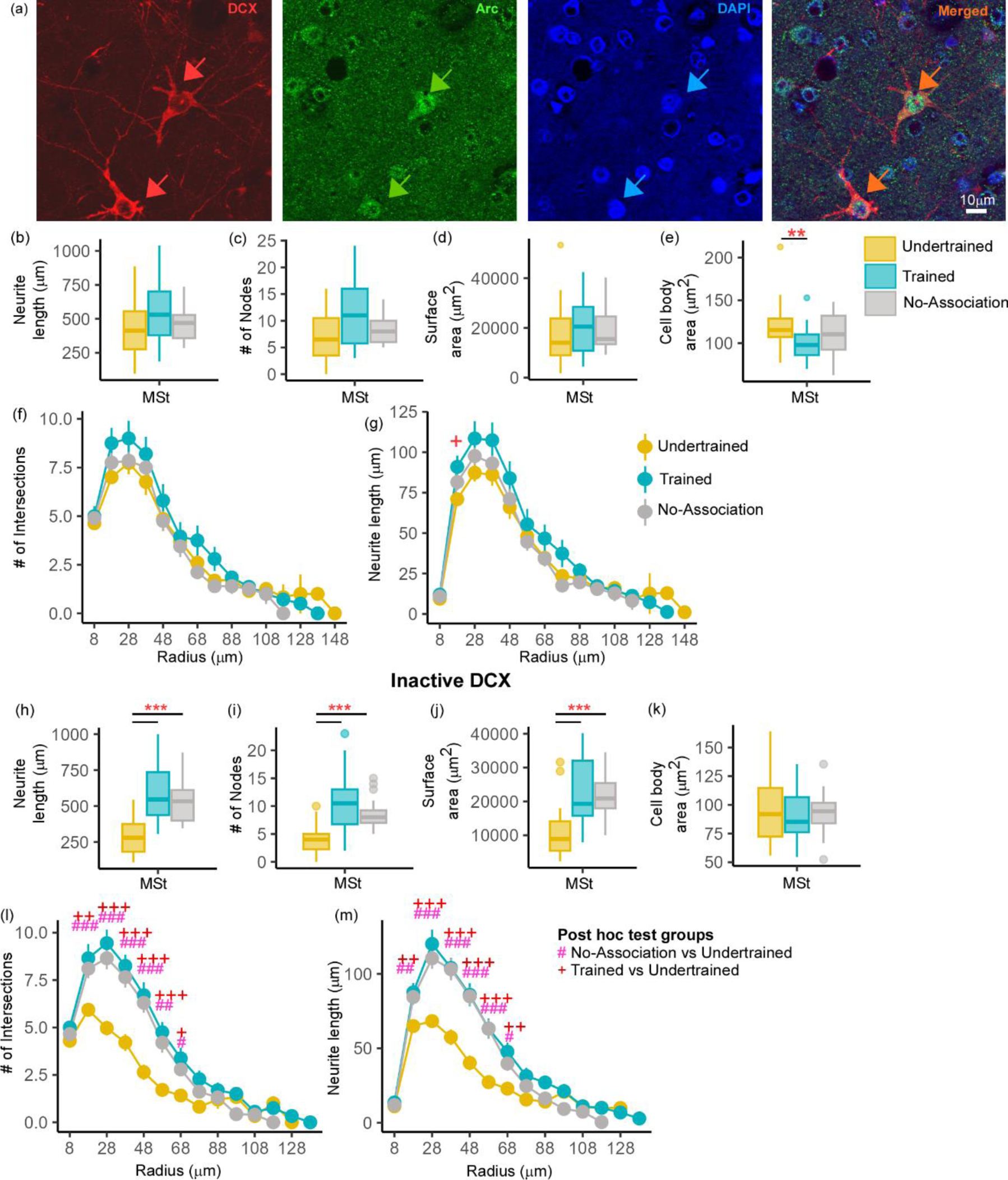
An analysis of changes in active and inactive DCX neurons in MSt. (**a**) A representative image from a Trained bird showing two DCX-labeled neurons (red; red arrows). The same section was labeled for Arc (green) and the nuclear label, DAPI (blue). The merged image demonstrates that these DCX-labeled neurons were positive for Arc and ‘active’. Very few double-labeled multipolar DCX neurons were found in MSt and the distribution of Arc was observed in the cytosol as well as in the nucleus of these neurons. Scale bar, 10µm. An analysis of these DCX and Arc-labeled neurons in MSt demonstrated no significant differences across Undertrained, Trained, and No-Association groups in terms of their (**b**) neurite length, (**c**) number of nodes, and (**d**) neurite field area. There was a significant decrease in the (**e**) area of somata in Trained versus Undertrained birds. Using Sholl analysis, we found that there were no changes in the (**f**) number of intersections and (**g**) and that the neurite length was significantly higher at only one Sholl radius in the Trained versus that in the Undertrained group. In the inactive DCX neuronal population in MSt, we observed significant increases in the (**h**) neurite length, (**i**) number of nodes, and (**j**) neurite field area in the Trained and No-Association groups compared to that in the Undertrained category. (**k**) There were no differences in the size of somata in any of the experimental groups. A Sholl analysis demonstrated an increase in the (**l**) number of intersections and (**m**) neurite length in the Trained and No-Association groups. */#/+ P<0.05; **/##/++ P<0.01; ***/###/+++ P<0.001.

##### 2.4.2.2. Structural changes in Arc- and DCX-positive neurons within MSt

Whereas there were no differences in neurite length, number of nodes or neurite field area [**Figure 7(b-d)**], neuronal somata of Arc- and DCX-double-labeled neurons in MSt were significantly larger in Undertrained versus Trained crows [P<0.01; **Figure 7(e)**; **Table S12**]. We also found no significant differences in the number of intersections [**Figure 7(f)**]. However, neurite length had increased significantly at one Sholl radius (18µm) in these neurons in Trained versus Undertrained birds [P<0.05; **Figure 7(g)**; **Table S13**].

In contrast, neurite length [P<0.001; **Figure 7(h)**; **Table S12**], number of nodes [P<0.001; **Figure 7(i)**; **Table S12**] and neurite field area [P<0.001; **Figure 7(j)**; **Table S12**] were significantly greater in inactive DCX-labeled neurons within MSt of Trained and No-Association birds versus that in Undertrained birds. Unlike activated neurons, there were no changes in the area of cell bodies of inactive DCX neurons across different groups [**Figure 7(k)**]. Besides these measures, Sholl analysis revealed significantly greater differences in the number of intersections and neurite length of DCX-positive neurons between Sholl radii 18-68µm of Trained and No-Association groups versus that in Undertrained birds [P<0.05, P<0.01 and P<0.001; **Figure 7(l-m)**; **Table S13**].

Comparisons between active and inactive DCX-labeled multipolar neurons in MSt revealed no significant differences in complexity or soma size in Trained crows [**Supplementary Figure** 13**(a-e)**]. Furthermore, the area of inactive DCX-positive somata was significantly lower than that of active neurons in No-Association birds [P<0.05; **Supplementary Figure** 13(e); **Table S14**]. Interestingly, neurite length (P<0.01), number of endings (P<0.001), number of nodes (P<0.01), neurite field area [P<0.01; **Supplementary Figure** 13**(a-d)**; **Table S14**] and cell body area [P<0.01; **Supplementary Figure** 13(e); **Table S14**] were significantly higher in active versus inactive DCX-labeled neurons of Undertrained birds. These findings suggest that active DCX-positive multipolar neurons in MSt increase in complexity specifically in Undertrained birds.

#### 2.4.3. Complexity of DCX-positive Neurons in NC

We found that neurite length, number of nodes, and neurite field area of multipolar DCX-labeled neurons in NC of Undertrained, Trained and No-Association crows were significantly greater than in Baseline controls [P<0.05, P<0.01 and P<0.001; **Supplementary Figure** 14**(a-c)**; **Table S15**]. Both Trained and No-Association group had significantly greater neurite length, number of nodes, and neurite field area compared to Undertrained birds in dNC (P<0.05, P<0.01, and P<0.001) whereas in iNC, these parameters were significantly greater only for No-Association versus Undertrained birds [P<0.05, P<0.01, and P<0.001; **Supplementary Figure** 14**(a-c)**; **Table S15**]. Neurite length and number of nodes of DCX-labeled neurons in mNCL were significantly greater in Trained versus Undertrained birds, whereas neurite field area was higher in both Trained and No-Association versus Undertrained birds in both mNCL and lNCL (P<0.05, P<0.01 and P<0.001; **Table S15**). In lNCL, branching was significantly greater in No-Association versus Undertrained birds whereas neurite length differences were observed for both Trained and No-Association versus Undertrained birds [P<0.05, P<0.01 and P<0.001; **Supplementary Figure** 14**(a-b)**; **Table S15**]. Similarly, neurite length and neurite field area of DCX-labeled neurons in vNC were significantly greater in No-Association versus Undertrained birds [P<0.05 and P<0.01; **Supplementary Figure** 14**(a and c)**; **Table S15**]. The area of DCX-positive neuronal somata was significantly greater in No-Association versus Baseline birds in dNC (P<0.05; **Table S15**) and also Trained versus Baseline birds in iNC [P<0.01; **Supplementary Figure** 14(d); **Table S15**]. Sholl analysis demonstrated an increase in the number of intersections and neurite length at most of the Sholl radii in Undertrained, Trained and No-Association versus Baseline birds in all subdivisions of NC [P<0.05, P<0.01 and P<0.001; **Supplementary Figure** 14**(e-h) and 15(a-f)**; **Table S16**].

##### 2.4.3.1. Morphometric analysis of active and inactive DCX-positive Neurons in NCL

Since there was greater activation in mNCL and lNCL following performance on the visual discrimination task, we decided to analyze Arc- and DCX-double labeled neurons specifically in these regions. Neurons positive for Arc and DCX were sparsely distributed in NCL [**Figure 8(a)**]. There were no changes in neurite length, number of nodes, neurite field area and area of the somata of these neurons in mNCL and lNCL [**Figure 8(b-e)**[or the number of intersections and neurite length measured by Sholl analysis based on experimental condition [**Figure 8(f-i)**].

**Figure 8.**
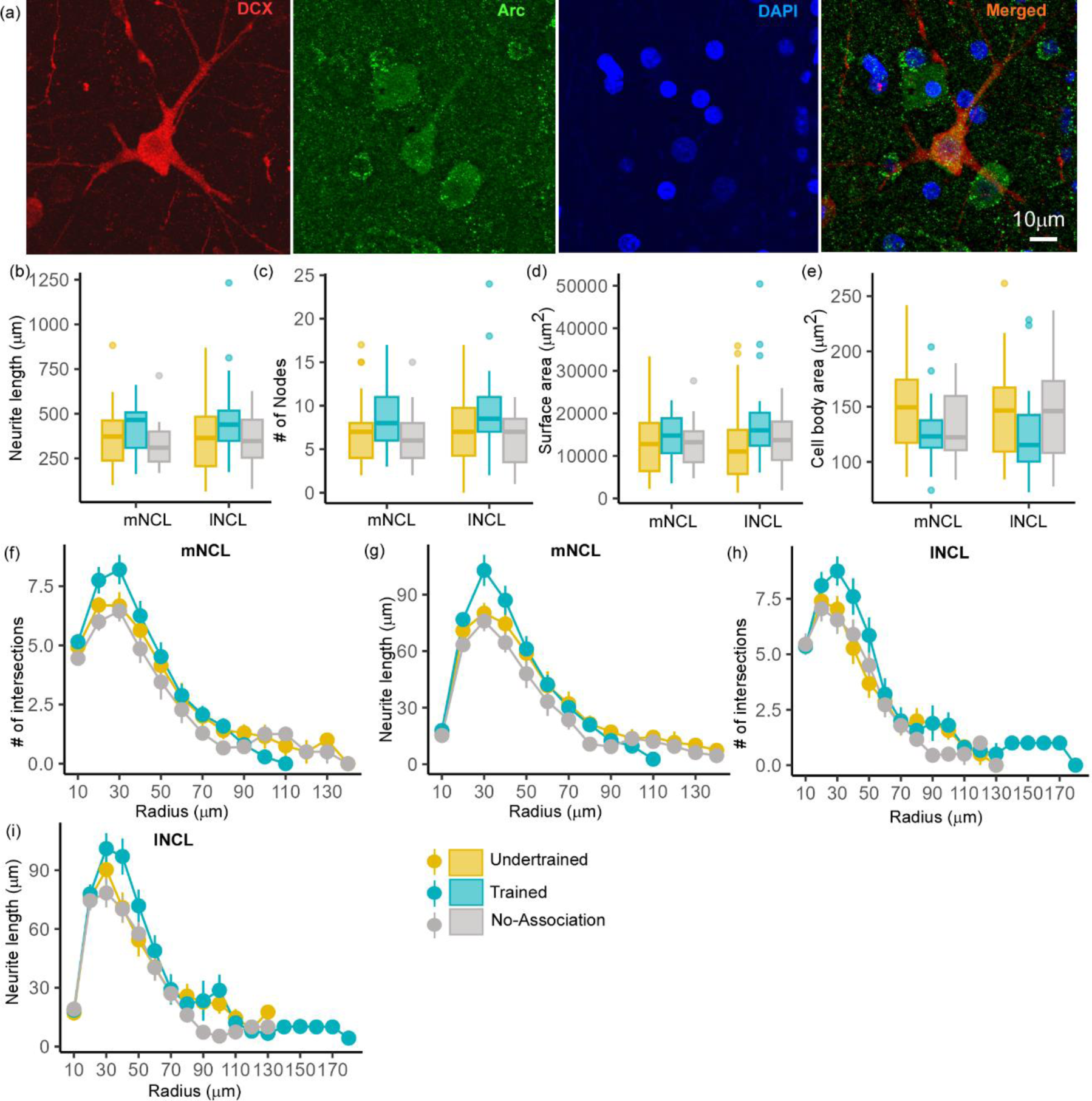
The structure of DCX and Arc double-labeled neurons in NCL. A representative neuron from the lNCL region of the Undertrained group showing colocalization of (**a**) DCX (red) with Arc (green), also labeled for DAPI (blue). Scale bar, 10µm. An analysis of changes in the structure of these active DCX neurons demonstrates that there were no significant differences in the (**b**) neurite length, (**c**) number of nodes and area (**d**) the neurite field, or (**e**) somata across different experimental groups. Whereas a Sholl analysis demonstrated that the (**f**) number of intersections in mNCL were greater in Trained birds versus those in other groups, these differences were not significant. There was a non-significant increase in the (**g**) neurite length of double-labeled mNCL neurons in the Trained versus the Undertrained and No- Association groups. A similar analysis of DCX and Arc double-labeled neurons in lNCL revealed that there were no significant differences in the (**h**) number of intersections or (**i**) neurite length. As seen for such neurons in mNCL, there were non-significant increases in DCX and Arc double-labeled neurons in the lNCL of the Trained group versus other groups.

In contrast, the complexity of inactive (Arc-negative) DCX-labeled multipolar neurons in lNCL and mNCL varied across different groups of experimental birds. The Dunn’s post-hoc test demonstrated that neurite length [P<0.001; **Figure 9(a)**; **Table S17**], number of nodes [P<0.01 and P<0.001; **Figure 9(b)**; **Table S17**], and neurite field area [P<0.01 and P<0.001; **Figure 9(c)**; **Table S17**] were significantly greater in Trained and No-Association versus Undertrained birds. There were no differences in the size of somata of inactive DCX-positive neurons in lNCL and mNCL in any of the groups [**Figure 9(d)**]. Lastly, Sholl analysis demonstrated that the number of intersections and neurite length of inactive DCX neurons was higher in Trained and No- Association crows versus that in Undertrained birds in both lNCL and mNCL [P<0.05, P<0.01 and P<0.001; **Figure 9(e-h)**; **Table S18**].

**Figure 9.**
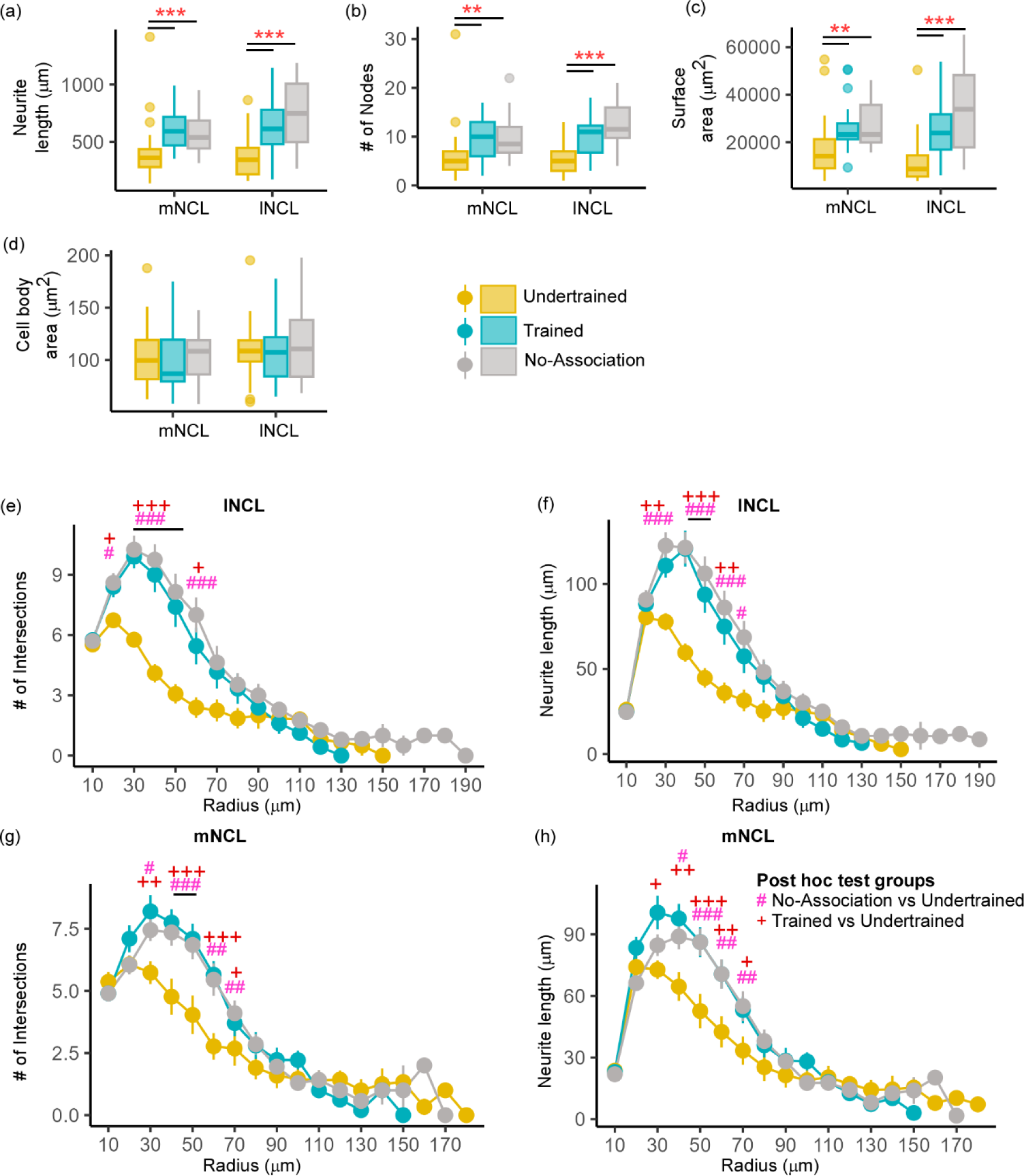
Structural changes in inactive DCX-positive neurons in the NCL. The (**a**) neurite length and (**b**) number of nodes were significantly greater in Trained and No-Association birds compared to those in the Undertrained category. (**c**) The neurite field of inactive DCX-labeled neurons in the Trained and No-Association groups was significantly larger in both mNCL and lNCL compared to that of Undertrained birds. (**d**) The area of the somata of these neurons was similar in all three groups. A Sholl analysis performed on reconstructed lNCL neurons demonstrated an increase in the (**e**) number of intersections and (**f**) neurite length in Trained and No-Association birds compared to that of Undertrained birds. Similar results for these parameters were observed for mNCL neurons, that is, an increase in the (**g**) number of intersections and (**h**) neurite length in Trained and No-Association birds versus that in the Undertrained group. */#, P < 0.05; **/##, P < 0.01; ***/###, P < 0.001.

A comparison of active and inactive DCX-labeled neurons across various experimental groups revealed that there were no significant differences in complexity based on neurite length, number of endings and nodes, and neurite field area in mNCL and lNCL of Undertrained birds [**Supplementary Figure** 16**(a-i)**]. However, in both these regions, the area of the somata of active DCX-positive neurons was greater than that of inactive DCX-positive neurons [P<0.001; **Supplementary Figure** 16(e) and (j)**; Table S19**]. In lNCL of Trained birds, the only significant change was an increase in neurite length of inactive versus active DCX-labeled neurons [P<0.05; **Supplementary Figure** 16(a)**; Table S19**]. In contrast, neurite length [P<0.01; **Supplementary Figure** 16(f)**; Table S19**] and neurite field area [P<0.001; **Supplementary Figure** 16(i)**; Table S19**] were significantly greater in inactive versus active DCX-labeled neurons in mNCL of Trained birds. However, the somata of inactive versus active neurons were significantly smaller in mNCL [P<0.01; **Supplementary Figure** 16(j)**; Table S19**]. All measures of complexity of neurites for inactive DCX-positive neurons were significantly higher in No-Association birds in both lNCL and mNCL [P<0.05 and P<0.001; **Supplementary Figure** 16**(a-i); Table S19**], although their cell bodies were smaller than those of active neurons in these regions [P<0.05 and P<0.001; **Supplementary Figure** 16(e) and (j)**; Table S19**]. These results suggest that active DCX-positive neuronal somata increase in size in all three groups, but the complexity of neurites increases significantly in inactive versus active DCX-positive neurons only in Trained and No-Association birds.

## 3. DISCUSSION

Whereas training on the visual discrimination task led to an increase in activation throughout NC in house crows based on Arc expression, we found that mNCL, lNCL, and iNC of Trained and No-Association birds were significantly more activated versus that in Baseline birds. The highest levels of neural activity overall were present in lNCL of Trained crows, which was similar to results in carrion crows (*Corvus corone*) demonstrating that NCL is involved in predicting behavioral rules (5), working memory (41), reversal learning (42), reward valuation (43), and performance on multicomponent behavioral tasks (7). Interestingly, training on visual discrimination leads to activation of other parts of NC including dNC, iNC and vNC in house crows, which needs further investigation. Besides NC, we found that MSNs and larger neurons (likely pallidal neurons and interneurons) were activated in different parts of the basal ganglia. Levels of activation were significantly higher in LSt and GP (important for motor functions in birds (44,45)) of Trained, No-Association, and Undertrained birds which attempted to obtain the reward versus Baseline controls. Furthermore, Area X and MSt, which are components of the anterior forebrain pathway in zebra finches (10) were activated in crows after training on the visual discrimination task. Whereas Area X is important for song crystallization (10,46) and context-dependent singing (47), MSt is activated when male zebra finches perform dance-like movements while courting females (48), in the selection of multicomponent behavior in pigeons (7), spatial and color-cued learning (6) and aversive learning (49). Interestingly, MSt receives projections from the parvicellular ‘shell’ of LMAN, which receives input from dNCL in zebra finches. Therefore, MSt in different species of songbirds may receive information about learning and decision-making processed in NC (50).

Both PFC and the striatum are involved in reinforcement learning (51,52) and are extensively connected in mammals and birds (53,54). Furthermore, local field potentials become more synchronized in these regions, suggesting that their connections are strengthened during learning (55). These findings suggest that similar mechanisms may underlie learning visual discrimination in crows.

### 3.1. Song Control Areas are activated following Visual Discrimination in House Crows

Surprisingly, brain regions generally associated with song control (10) were activated after house crows performed the visual discrimination task. The song control area LMAN was significantly activated in No-Association birds versus other groups of birds. In songbirds such as zebra finches, LMAN is important for generating variability in vocalizations during the sensitive period for song learning (46,56). Recent studies further demonstrated that LMAN lesions prevented somatosensory-based non-auditory learning which affected vocal output in zebra finches (57) and an LMAN-like region in pigeons (NIML) was involved in the execution of sequence learning (58) and serial processing (9), but not in generating variability. In our study, trial-by-trial variability would be the highest in No-Association birds since the reward was randomly associated with either of the shapes presented during the task. These findings suggest that besides vocal learning, LMAN is involved in modulating variability associated with learning the visual discrimination task in house crows.

We also observed higher activation in RA and AId of Trained crows versus other groups. Besides projecting to syringeal musculature for controlling vocalization, RA projects to the ventral respiratory column for controlling respiration (37). Since experimental crows never vocalized during or immediately after training, it is possible that this heightened activity in RA may be important for synchronizing breathing with performing the correct sequence of actions during the task. Another arcopallial region, AId, is involved in motor functions and song learning (59). It receives topographically organized projections from NCL (50,60) and projects to the optic tectum (61). Increased activation of AId consequent to training on visual discrimination may involve a goal-directed visuomotor pathway beginning in mNCL (unpublished data) which is connected to AId and the optic tectum (60,61).

### 3.2. Dopaminoceptive neurons are associated with Learning in House Crows

Dopamine plays an important role in motivation, learning, cognition, reward and pleasure, and motor learning and the firing rates of VTA-SNc increase to signal the physical salience of the reward and reward-predicting stimuli (20). As expected, dopaminergic neurons in VTA-SNc were positive for Arc in all groups of birds other than the unrewarded Baseline controls.

Furthermore, NCL contained DARPP-32-labeled dopaminoceptive neurons of which some were positive for Arc, showing that they participated in visual discrimination. Although there were no changes in their number, for the first time, we have demonstrated that the complexity of their neurites and soma size increased significantly across NC, especially in mNCL and lNCL of Trained house crows. These findings suggest that learning to associate specific types of behavior with a reward leads to an increase in the plasticity of dopaminoceptive neurons. Similar changes in the dendritic field of pyramidal neurons were observed in mPFC and OFC of rats trained on the T maze or parallel alley maze (24).

We found that both active and inactive dopaminoceptive neurons in mNCL and lNCL were more complex in Trained and No-Association birds versus Undertrained birds, suggesting that training on visual discrimination led to an increase in branching and likely, the number of synapses.

Comparisons between these sets of neurons revealed that inactive dopaminoceptive neurons were more complex than active ones in lNCL and mNCL of Trained and No-Association birds.

However, active neurons in mNCL were more complex than inactive ones in Undertrained birds (**Figure 10(a)**). These findings suggest that the initial phase of learning leads to an increase in complexity and/or synapses of DARPP-32-labeled neurons in mNCL, whereas extensively branched neurites of dopaminoceptive neurons in lNCL and mNCL are pruned to retain only task-specific connections (62) with training.

**Figure 10.**
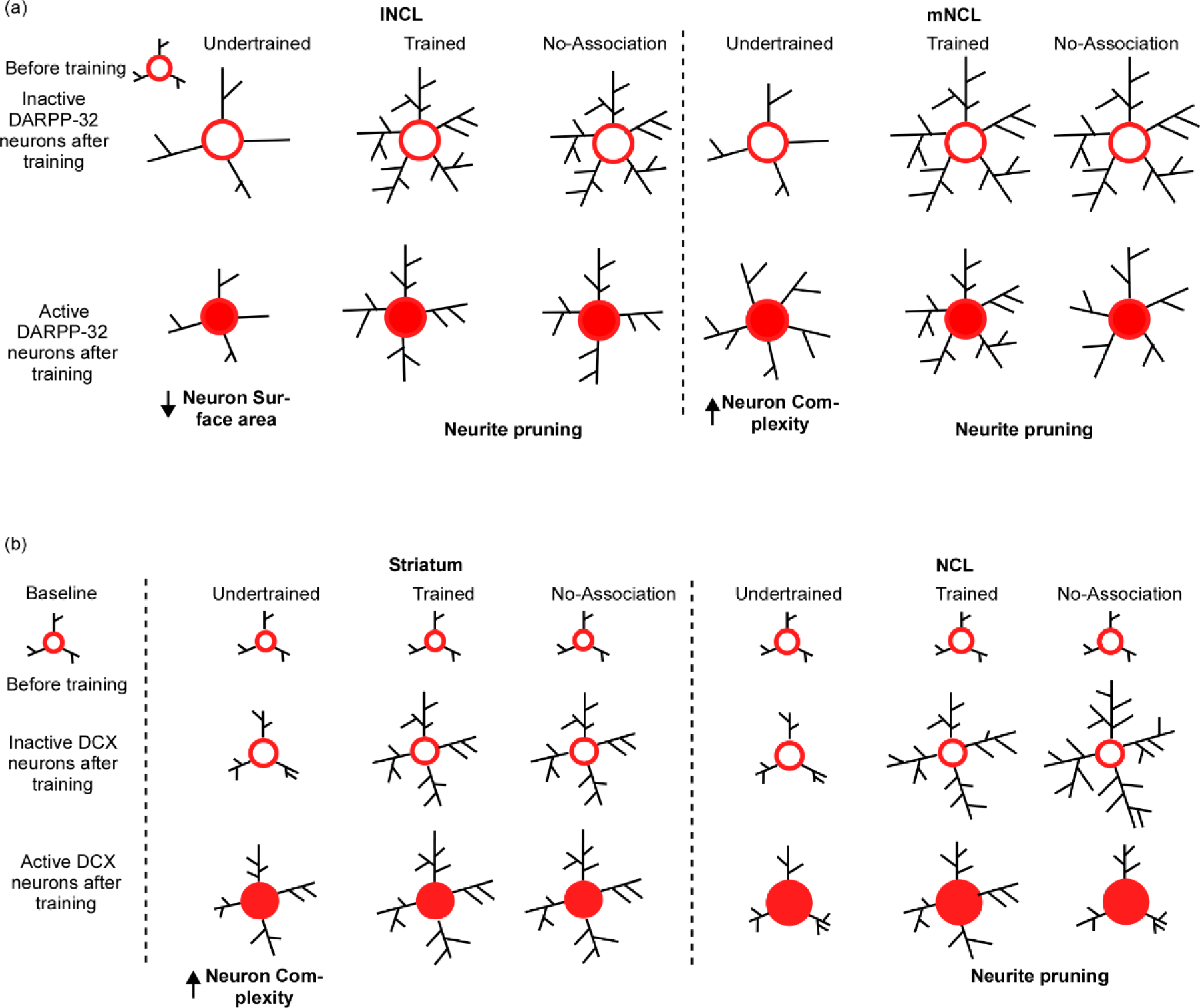
Possible mechanisms underlying changes in the structure of DARPP-32 and DCX neurons in the striatum and NCL for visual discrimination task. **(a**) We assume that dopaminoceptive (DARPP-32-positive) neurons in lNCL and mNCL regions before training (in the Undertrained, Trained, and No-Association groups) are similar to those in the Baseline group. After training, there is an increase in the neurite field in inactive dopaminoceptive neurons in the lNCL of the Undertrained group, which is less than that of active Arc- and DARPP-32-positive neurons. In contrast, active neurons demonstrate an increase in neuronal complexity in mNCL. Inactive (Arc-negative) DARPP-32 neurons in both lNCL and mNCL from Trained and No-Association groups are more complex than active (Arc-positive) dopaminoceptive neurons in these regions. Since in each case, active neurons have fewer neurites, it is possible that they are pruned as a result of learning the visual discrimination task. In the striatum (MSt), (**b**) the structure of immature DCX-positive neurons in all experimental groups before training on the visual discrimination task is likely similar to that in the Baseline group. With training (in the Trained and No-Association groups), there is an increase in the complexity and size of somata of DCX-positive neurons which are not part of active neural circuits. Whereas immature neurons which appear to be incorporated into active neural circuitry underlying the task in the Undertrained group become more complex, there was no difference in their structure in the Trained and No-Association groups. Similar changes were observed in DCX-labeled neurons in NCL. However, unlike MSt neurons, there is a decrease in complexity of these neurons suggestive of neurite pruning in Trained and No-Association birds.

Furthermore, despite the fact that neurons double-labeled for DCX and Arc (a part of the active circuit) underwent changes in their structure in Undertrained birds, these changes were not significant and were not as pronounced as those in MSt.

### 3.3. Learning-induced Adult Neurogenesis and Structural Changes in Adult-born Neurons in the House Crow brain

Learning may lead to neurogenesis in house crows (29) and other species (63–65) even in adulthood. For example, the transient depletion of immature neurons in the mouse hippocampus causes deficits in learning the active place avoidance task (66) and they are important for reconsolidation of task-induced memories (67). Amongst avian species, both fusiform and multipolar DCX-labeled neurons increase in number in the nidopallium of adult house crows (29) and pigeons (40), which was correlated with stress and homing behavior, respectively. In our study, the increase in immature DCX-labeled neurons in the striatum and NC of Trained versus Undertrained and Baseline categories of house crows may be necessary or permissive for learning and decision-making. Furthermore, the significant increase in spherical DCX-labeled neurons in different compartments of the avian basal ganglia (Area X, MSt and LSt) in Trained birds versus other groups may contribute to habitual learning and habit formation (68), learning extinction (69,70), reinforcement learning (52) and decision-making (35) besides vocal learning (46), as seen in the mammalian dorsal striatum. Interestingly, striatal lesions and exposure to an enriched environment have been linked to an increase in migrating DCX-positive neurons in the dorsal striatum of rodents as well (65).

Besides an increase in adult neurogenesis, DCX-labeled multipolar neurons were significantly more complex in different divisions of NC, Area X and MSt of Trained and No-Association birds versus those of other groups. In NCL, inactive DCX-positive neurons were more complex and appeared to be pruned when they became parts of active circuits. Whereas active DCX-labeled neurons in MSt demonstrated fewer changes in complexity across different experimental groups, they were more complex compared to inactive DCX-labeled neurons in MSt of Undertrained birds (**Figure 10 (b)**). In rodents, the dorsomedial part of the striatum is involved in attentive decision-making, whereas its dorsolateral component underlies automatizing responses during the latter part of the learning process (71–73) and neurons in these regions undergo changes in dendritic complexity with learning (62). Changes in the No-Association group were similar to those in the Trained group probably due to the trial-and-error learning method employed by the crows. Overall, these results suggest that inactive neurons become more complex in MSt in Undertrained crows during the initial part of learning the visual discrimination task.

### 3.4. Why are morphometric changes in DARPP-32- or DCX-positive neurons in No-Association crows similar to those in Trained birds?

An interesting conundrum is provided by No-Association birds. Despite being exposed only to two trials (similar to Undertrained birds) of the visual discrimination task, structural changes undergone by DARPP-32- or DCX-labeled neurons in the No-Association group are comparable to those observed in Trained birds. It is possible that employing two extreme strategies for obtaining the food reward, that is, goal-oriented behavior in case of Trained birds and trial-and-error learning in No-Association birds results in similar changes in the complexity of neurons involved in the visual discrimination task. Alternatively, the strengthening of functional connections coding for the correct strategy to obtain rewards may vary across different experimental groups. For example, neural circuits in Undertrained birds, which are at an initial phase of learning may not have been strengthened leading to low and variable neural activity and fewer changes in the complexity of mature and immature neurons in the striatum and NCL. Following extensive training on the task, it is likely that neural circuits responsible for obtaining the food reward may have been strengthened, leading to higher levels of neural activity and more elaborate neurites in neurons within NCL. Finally, in the No-Association group, since rewards are associated randomly with the shapes, the cognitive load may be the highest. Hence, high levels of neural activity as well as morphometric changes in DCX-labeled neurons of No-Association birds are similar to those in Trained birds.

### 3.5. Are changes in activity and structural differences in dopaminoceptive and DCX-labeled neurons due to visual discrimination or motor behavior?

We have shown that levels of neural activity in GP and LSt (important for motor activity) were similar across Trained, Undertrained and No-Association birds. Furthermore, neural activity in these regions was higher in the three experimental groups compared to that in Baseline birds, which spent little time in exploring the behavioral apparatus. These results suggest that neural activity in these regions of the avian basal ganglia was probably due to the motor sequence involved in retrieving the reward rather than learning the task strategy itself. Since levels of neural activity in Area X and MSt were higher in Trained and No-Association birds compared to Baseline and Undertrained birds, our results suggest that the medial striatum appears to be involved in learning the appropriate strategy to perform the task.

A recent study on carrion crows has demonstrated that neurons in NCL also code for motor planning and execution during goal-oriented behavior (74). In our study, we did not find any difference in the levels of activation in NCL between birds of the Trained and No-Association groups despite the fact that they likely employ different strategies for learning. That is, Trained birds appear to employ a goal-oriented strategy to obtain the reward. In contrast, No-Association birds appear to adopt a trial-and-error learning strategy, since they cannot predict the presence of the reward by observing the shape of the blocks provided. Despite having to follow an ‘explore all’ strategy, the patterns of neural activity in NCL of these birds was similar to that in the Trained group. Our results also demonstrate that there were significant changes in the complexity of dopaminoceptive and immature neurons in MSt and NCL of Trained and No-Association birds compared to that in Baseline and Undertrained birds. Taken together with the data on neural activity, these structural changes appear to be the result of learning the visual discrimination task rather than motor activity.

### 3.6. Conclusions

Our results suggest that diverse brain areas are involved in learning visual discrimination in corvids. Furthermore, increased adult neurogenesis and structural changes in dopaminoceptive and immature neurons may be correlated with learning in the house crow brain.

## 4. Materials and Methods

A total of 18 adult house crows (n = 12 males and 6 females) were used for the shape discrimination experiments. All experimental birds were wild-caught with the permission of the Chief Wild Life Warden, Haryana, and housed in aviaries at the Animal Facility, National Brain Research Centre, Manesar, and experimental protocols were approved by the Institutional Animal Ethics Committee, NBRC. All experimental procedures used for these studies were carried out according to guidelines laid down by the Committee for the Control and Supervision of Experiments on Animals (CCSEA), India, compliant with international standards on animal welfare. At the end of the experiments, tissue from the liver of the house crows was genotyped to determine their sex, using a previously described protocol (75).

### 4.1. Behavioral Setup

All house crows were weighed prior to starting behavioral training. Experimental crows were housed in an outdoor aviary with natural day and light conditions . Two days prior to training on the visual discrimination experiments, they were transferred to a cage (dimensions: 30′′ × 21′′ × 34′′) in a separate room also maintained in natural light and dark conditions for habituation. On the second day of this period, they were food-deprived from 4 pm in the evening until the beginning of the pretraining period which started at 9 am the next day. Birds were given ad libitum access to water but the amount of food given post-behavioral training depended on the appetite of each bird, since some of the birds did not participate in the training paradigm if they were fully satiated the previous evening. For the food reward, we used pieces of dried shrimp, chicken sausage, or vitamin-fortified white bread depending on the crows’ preference. A total of two blocks of the shape discrimination task comprising 12 trials each were conducted every day with a gap of 4-5hrs, 5 days a week. Over the weekend, experimental birds were transferred back to their aviaries where they were in visual and auditory contact with other crows and provided eggs, bread, dog food (Pedigree), and water ad libitum. There was no reduction in the body weight (∼250gm) of any of the house crows at the end of training on the behavioral paradigm.

#### 4.1.1. Pre-training

All experimental crows were initially trained to retrieve food rewards placed on a platform in the cage to habituate them to the apparatus used for the behavioral experiments. After birds learned to retrieve food from the platform, food rewards were partially hidden by two three-dimensional plastic blocks with the same surface area (a triangle and a circle). With further training, birds learned to retrieve food from under the blocks, after which they were trained for the visual discrimination experiment.

#### 4.1.2. Training

During the training trials, the experimenter placed the two blocks on the platform in the cage, closed the cage door, and hid behind a screen outside the visual range of the crows, which marked the beginning of a trial [**Supplementary Figure** 1(b)]. The positions of the shapes were randomized in the cage to prevent birds from associating the reward with spatial locations. A trial was considered complete when the crow hopped down from its perch, knocked over a shape to retrieve the food reward, and returned to the perch or at the end of two minutes, whichever was sooner. If crows did not attempt to retrieve food rewards in three consecutive trials, the block of experiments was stopped and if crows started attacking the camera, that trial was stopped.

The four groups used for our experiments are as follows:

1. Trained group (n = 4): The food reward was placed only under the triangular block. Birds were trained to retrieve food until their success rate on the task reached 80% or above in a block. After two consecutive blocks during which crows’ performance remained at 80% or above, they were considered fully trained.
2. Undertrained group (n = 4): The food reward was placed only under the triangular block. For this group, the experiment ended when they achieved a success rate of ∼ 40 to 60%.
3. No-Association group (n = 6). The food reward was randomly presented under either shape and the experiment ended after the birds in this group received two blocks of training. In each session, each shape was rewarded 50% of the times. Birds from Groups 1-3 were kept in the dark for one hour to minimize neural activity resulting from other visual stimuli at the end of a block of trials.
4. Baseline group (n = 4): Crows were placed in the cage for behavioral assessment and exposed to the shapes placed on the platform for 6hrs on 2 consecutive days without food deprivation. On the third day, they were exposed to the behavioral setup for 30min after which they were placed in the dark prior to ending the experiment. In this group, birds were exposed to the apparatus for a long period in order to saturate them with the visual stimuli elicited by the behavioral setup. As a result, we did not expect to observe neural activity in their brains due to the novelty of the visual stimuli.

At the end of the last block of trials, birds were kept in the dark for 90min and anesthetized with an overdose of ketamine (30 mg/kg) and xylazine (2mg/kg). This was followed by intracardial perfusion with 0.01M phosphate-buffered saline (PBS), followed by 4% paraformaldehyde (PFA). Brains were removed and post-fixed with 4% PFA (for one week at 4°C), after which they were cryopreserved in 30% sucrose and cryosectioned serially at 50µm in the coronal plane. These sections were later used for immunohistochemistry.

### 4.2. Behavioral analysis of motor activity

We calculated the time taken to make a decision during each trial for the Trained, Undertrained and No-Association groups. Each trial was initiated by the experimenter moving away from crows’ line of sight. This was because crows did not generally dismount from the perch if they could still see the experimenter. Trials were considered to end when crows knocked over a shape to retrieve the reward. The time interval from the beginning of the trial until knocking over the shape was quantified, as it represents the duration for motor activity relevant to learning.

### 4.3. Western blotting

Immunoblotting was performed to check the validity of the DCX antibody (sc-271390, Anti-Doublecortin Antibody (E-6); RRID: AB_10610966, Santa Cruz Biotechnology) using samples of tissue from the anterior striatum of a house crow (n = 1, female). The tissue was homogenized in SDS lysis buffer and sonicated with 25 pulses of 1s each (thrice with an interval of 5s), followed by centrifugation at 12000rpm at 4° C. The supernatant was collected and protein was estimated using the bicinchoninic acid protein estimation method (BCA, B9643, Sigma-Aldrich). We separated a protein sample (80µg) on an SDS-PAGE gel (11% Acrylamide-Bisacrylamide gel concentration). The resolved proteins were transferred to a nitrocellulose membrane which was blocked with 5% BSA (Bovine serum albumin) for 2hrs. This was followed by incubation in the primary antibody solution (1:1000, anti-DCX) for 13-16 hrs at 4° C. The primary antibody was rinsed using six washes of TBST (Tris borate saline with 0.1% Tween 20, 10min each) followed by incubation in a secondary antibody solution (1:3000, peroxidase labeled anti-mouse, PI-2000, Vector laboratories). This was followed by rinsing in TBST, after which the blot was developed with the ECL chemiluminescent reagent (WBKLS0500, Immobilon Western Chemiluminescent HRP Substrate, MERCK, USA). We obtained an intense band at ∼40kD and a very faint one at ∼32kD (**Supplementary Figure** 1(c)) when a western blot was performed on lysate from a house crow brain using an anti-DCX antibody (sc-271390, Santa Cruz Biotechnology). Bands were obtained at similar molecular weights in western blots performed on mouse brain tissue provided by the manufacturer.

### 4.4. Immunohistochemistry

#### 4.4.1. Arc, DCX, TH and DARPP-32

Coronal serial sections (50µm thick) from the right hemisphere were labeled using immunohistochemistry for Arc (Cat# ab85656, Abcam, RRID: AB_1924788), Tyrosine hydroxylase (Cat# MAB318, Merck-Millipore, RRID: AB_2201528) or DARPP 32 (Cat# ab- 40801, Abcam; RRID: AB_731843). Sections from the left hemisphere were used for immunohistochemistry against DCX. The antibodies used to detect Arc, TH, and DARPP-32 have been previously used in songbirds (36,76). Sections were incubated in an antigen unmasking solution (H-3300, Vector laboratories) for 30min at 80° C in a water bath, followed by rinsing with 0.01M PBS (only for DCX). After rinsing with 0.01 M PBS, sections were incubated in 1-3% H_2_O_2_ in 0.3% Triton-X 100 for 30 minutes to quench endogenous peroxidase activity. They were incubated in a blocking solution [5% normal goat serum, NGS, for Arc; 5% NGS and 2% Bovine serum albumin, BSA, for DARPP-32 and 5% Normal horse serum, NHS, and 2% BSA for TH and DCX; S-1000 (NGS) and S-2000 (NHS), Vector Laboratories, Burlingame, CA] for 2 hrs. This was followed by incubation in a solution containing the primary antibody [Arc; 1:1000; (incubation for 38-40hrs at 4°C); DCX; 1:500 (incubation for 16-20 hrs at 4°C); TH; 1:200 (incubation for 38-40hrs at 4°C) or DARPP-32; 1:500 (incubation for 16-20 hrs at 4°C); made in blocking buffer containing 0.3% triton-X PBS]. This was followed by rinsing the sections in PBS and incubating them in a secondary antibody solution (biotinylated anti-rabbit for Arc and DARPP-32 and biotinylated anti-mouse for DCX and TH; 1:200; BA-1000, BA-2000 Vector Laboratories) for 2hrs. After rinsing in PBS, sections were incubated in a solution containing avidin-biotin complex (ABC reagent; PK-6100, Vectastain Elite ABC HRP kit, Vector Laboratories; 1:50) for 2hrs. Sections were again rinsed in PBS and then developed in a solution containing the chromogen [Nova Red peroxidase (HRP) substrate kit (SK-4800, Vector Laboratories)] according to the manufacturer’s instructions. Finally, sections were rinsed with Milli Q and mounted on gelatin-coated slides, after which they were air-dried overnight and cover-slipped with DPX. Representative images for demonstrating staining patterns were adjusted only for brightness and contrast.

### 4.5. Double immunofluorescence

#### 4.5.1. Arc and DARPP-32

In order to study whether DARPP-32-positive neurons were activated as a result of the visual discrimination task, sections from the crow brain from different groups were double-labeled for DARPP-32 and Arc. Coronal serial sections at the level of the caudal nidopallium were used for these experiments. A different antibody was used for Arc (Cat# NBP1-56929, Novus Biologicals, RRID: AB_11010941) rather than the one used for single label (see above) due to technical problems associated with immunofluorescence. Before testing for double-immunofluorescence, we tested this antibody singly and found no difference in the expression patterns of the two Arc antibodies. For double immunofluorescence, the basic steps of quenching and permeabilization were performed as described above. Sections were then blocked with 5% NGS for 2hrs, followed by incubation in the primary antibody solution (1:500, anti-DARPP-32 made in rabbit 5% NGS) for 38-40hrs at 4°C. Sections were rinsed and incubated in the secondary antibody solution (1:200, anti-rabbit; Alexa-594, A11012, ThermoFisher Scientific, USA) for 4- 5hrs at room temperature. Later, they were washed with PBS and blocked with 5% NGS and 2% BSA for 2hrs, followed by incubation in the anti-Arc primary antibody (1:200, anti-Arc antibody made in rabbit in blocking buffer with 0.3% triton-X) for 38-40hrs at 4°C. Sections were washed with PBS and incubated in the secondary antibody solution (1:200, Alexa-488 anti-rabbit; A11008, ThermoFisher Scientific, USA) for 4-5hrs followed by three washes in PBS. Sections were then transferred onto gelatin-coated slides and cover-slipped with antifade DAPI mounting media (H-1800, Vector laboratories).

#### 4.5.2. Arc and DCX

To study whether DCX-positive neurons were activated following the visual discrimination task, serial coronal sections of the house crow brain at the level of MSt, Area X, and caudal nidopallium were double-labeled with Arc and DCX. Sections were incubated in the antigen unmasking solution for 30 minutes at 80° C in a water bath followed by rinsing with PBS. The steps for quenching and permeabilization were performed next, as described above. Sections were then blocked with 5% NGS for 2hrs, followed by incubation in the primary antibody cocktail (1:200, anti-DCX made in mouse and 1:200, anti-Arc) for 38-40hrs at 4°C. Sections were rinsed and incubated in the secondary antibody cocktail (1:200, anti-mouse; Alexa-594, Cat. # A11005 and 1:200, Alexa-488 anti-rabbit, Cat. # A11008; ThermoFisher Scientific, USA) for 4- 5hrs at room temperature. After this step, they were washed with PBS, transferred onto gelatin-coated slides, and mounted with antifade DAPI mounting medium (H-1800, Vector laboratories).

### 4.6. Quantitative Analysis of Tyrosine hydroxylase positive profiles

Sections at the level of NCL were stained with TH and demarcated into various subdivisions according to an earlier report from our lab (36). In each subdivision, the area of TH-positive profiles was determined using ImageJ (version: 1.52). The contours of different subdivisions in NC (dNC, iNC, mNCL, lNCL, and vNC) determined using TH staining patterns were used to determine the boundaries of NC subdivisions in Arc and DARPP-32-stained sections. Each image was converted to 8 bits and thresholded based on staining in lNCL, followed by quantifying the TH-positive profiles in different subdivisions of NC. We normalized the % area fraction of the five NC subdivisions with a positive control, that is, a band of TH-positive profiles located between LAD (Lamina arcopallialis dorsalis) and AId.

### 4.7. Neuron counts

A contour was drawn around the area of interest in serial sections using the Stereoinvestigator software (Microbrightfield, Williston, VT) linked to an Olympus microscope (BX-51). The optical fractionator method (77) was used to count Arc-, DCX-, and DARPP-32-positive cells. Lightly and intensely stained Arc-positive neurons were counted separately. The Arc-positive neurons were counted at 100X using a sampling grid of size 70µm × 70µm with an area sampling fraction (asf) of 5 or 10 based on the size of the area and a dissector height of 9µm with 2µm guard zones. The DCX and DARPP-32-positive neurons were counted at 40X using a grid size of 175µm × 175µm with asf of 50, 30, or 20 and dissector height of 10µm with 2µm guard zones.

Additionally, we counted the three different neuronal populations stained for DCX, namely, multipolar, fusiform, and spherical. For all fractionator counts, every 6th section was sampled. An estimated count using mean section thickness for Arc, DCX, and DARPP-32 was used to compare the neuronal population across various training groups. To accommodate size differences across brain regions between different birds, we normalized the estimated counts by the number of bins from each area, which is an indicator of the size of the area.

### 4.8. Fluorescence imaging and neuron counting

Fluorescence imaging for sections stained for Arc and DARPP-32 were performed using an Apotome microscope (Carl Zeiss1, AxioImager.z1). We imaged z-stacks (magnification: 40X for counting and 63X for imaging) at an interval of 1.5µm from different subdivisions of NC from all crows used in our experiments on visual discrimination. Representative images for demonstrating single and double label were adjusted only for brightness and contrast. Manual neuronal counts for Arc and DARPP-32 double-positive neurons were performed from these images using ImageJ software.

### 4.9. Morphometric Analysis of DARPP-32 and DCX multipolar neurons

We traced 15 DARPP or DCX positive neurons from each subdivision of the caudal nidopallium from each bird using the Neurolucida software (version:11, MBF Bioscience, USA) linked to a microscope (Olympus BX51) at a magnification of 100X. After a careful visual inspection, only those neurons which possessed intact processes were selected for tracing. Since we could not discriminate between the axons and dendrites of neurons based on staining for DARPP-32 or DCX, we have considered all processes as neurites for our study. Traces were exported to the Neurolucida Explorer software and subjected to three morphometric analyses, including 1) Neuron summary analysis for overall changes in the neurite branching and length, 2) Convex hull analysis to quantify changes in the neurite field, and 3) Sholl analysis for quantifying neuronal complexity, wherein the traced neuronal soma was placed at the center of a set of concentric circles at a fixed distance (10µm). The number of intersections that the neurite tree made with these circles was counted and for each Sholl radius, the total neurite length was analyzed.

Statistical analysis was performed on changes observed between the radii from 20–70µm. However, data from changes within the 10µm Sholl radius were not analyzed since we observed an increase in the size of somata of DARRP-32 and DCX positive neurons in Trained, Undertrained, and No-Association birds compared to Baseline controls and the inclusion of these data might introduce a false estimation of the analyzed parameters. The initial Sholl radius was reduced to 8µm for DCX neurons from the striatum due to the smaller size of their somata.

Similarly, five Arc and DARPP-32/DCX double-positive neurons each from MSt, mNCL and lNCL regions were traced using the Neurolucida software (version 2020.1.3; MBF Bioscience, USA) linked to a fluorescence microscope (Olympus BX53). The three-dimensional tracings of these neurons were analyzed in the same manner as described above for individually labeled neurons using the Neurolucida Explorer software.

### 4.10. Statistical Analysis

The R (version 4.1.1) software was used to perform statistical tests and visualize the data. In order to check the interaction effects between area and experimental conditions for neuronal counts and the morphometry data, we performed a two-way ANOVA. For non-normal data, an aligned rank transform was performed, followed by two-way ANOVA (78). For group comparisons, if data was found to be normally distributed (using the Shapiro-Wilk test) and possess equal variance (using the Levene test), a one-way ANOVA was performed. If the data did not pass these tests, a Kruskal-Wallis test was performed for judging statistical significance. In both cases, Tukey’s (multcomp library; adjusted P values are generated by the single-step method) or Dunn’s post-hoc test (FSA library using the Holm method for P value adjustment) was performed for multiple comparisons. For comparisons involving only two groups in some cases, a Welch’s two-tailed t-test or Wilcoxon rank sum test was performed.

## Data Availability Statement

All relevant data are within the manuscript and its Supporting information files.

## Acknowledgments

We would like to express our gratitude to Krishan Sharma (NBRC, Manesar) for his help with the experiments.

## Author contribution

P.P. performed the behavioral experiments, immunohistochemical staining, microscopy, neuronal counting and tracing, morphometric analyses, statistical analysis, manuscript writing and helped in study design. M.R. performed the neuronal tracing and analysis of brightfield DCX neurons in caudal nidopallium. S.I. designed and conceptualized the study and was involved in writing and reviewing the manuscript.

