## Supplementary Figure 1 for "Changes in the Dopaminergic circuitry and Adult Neurogenesis linked to Reinforcement Learning in Corvids"

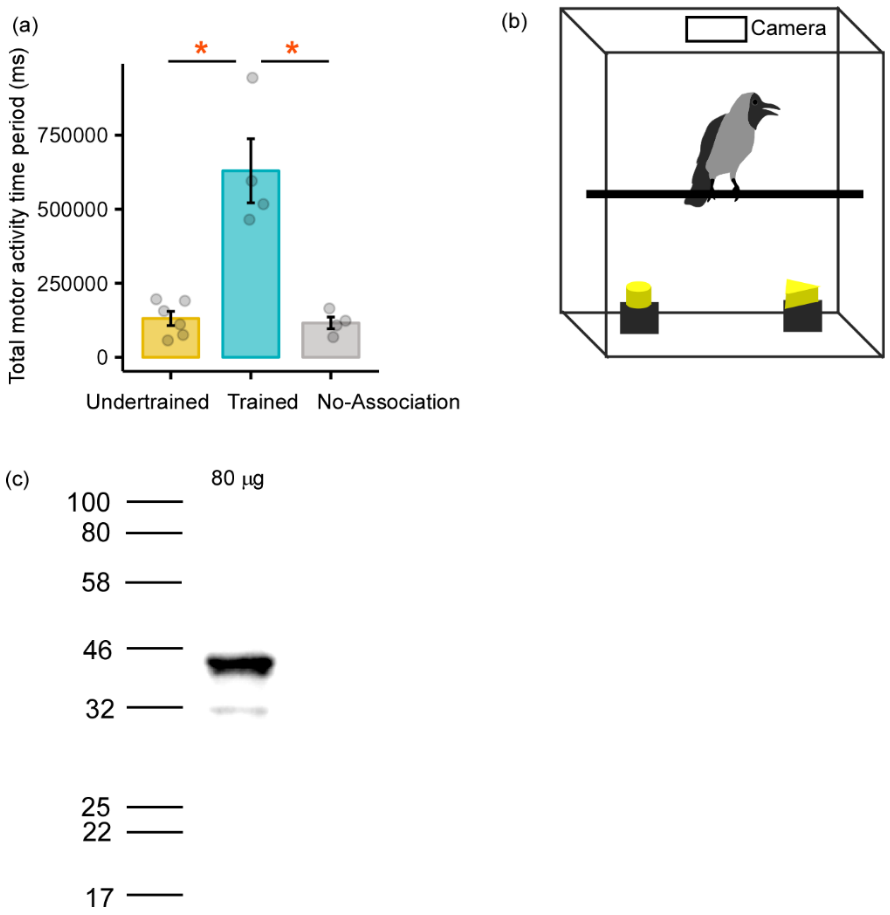


**Supplementary Figure 1**. A schematic of the behavioral setup and the validation of DCX antibody.

(**a**) There were significant differences in the total period of activity related to learning the task in Trained birds versus those in the Undertrained and No-Association groups. *, $P < 0.05$. (**b**) A schematic of the behavioral setup, demonstrating the position of the camera, perch, and the two shapes positioned inside the cage. (**c**) An intense band was obtained at ~40kD by performing a western blot on house crow brain tissue, demonstrating the specificity of the DCX antibody (sc-271390, Anti-Doublecortin Antibody (E-6); RRID: AB_10610966, Santa Cruz Biotechnology).
