## Supplementary Figure 2 for "Changes in the Dopaminergic circuitry and Adult Neurogenesis linked to Reinforcement Learning in Corvids"

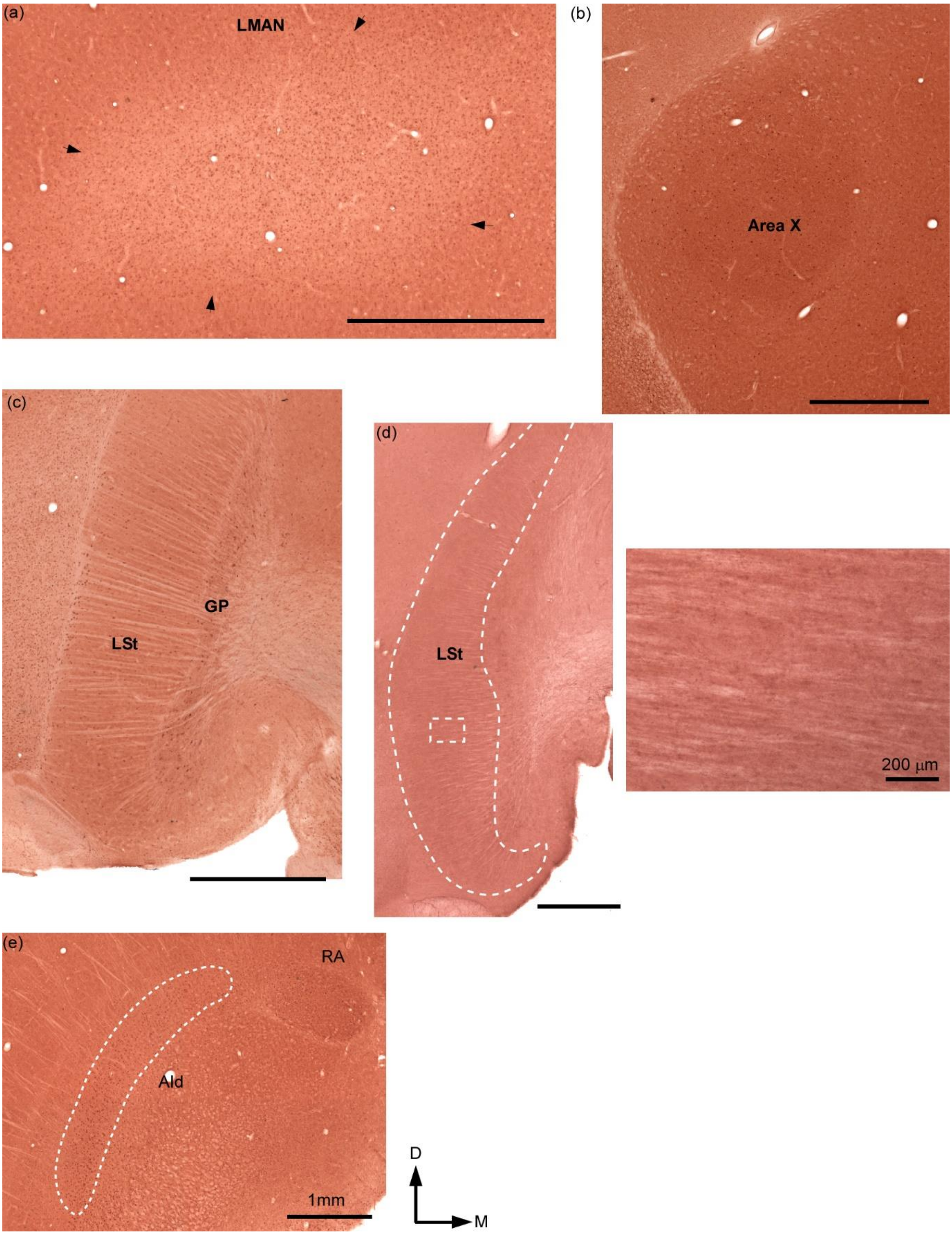


**Supplementary Figure 2**. Arc staining patterns in different brain regions.

(**a**) The boundaries of LMAN (*arrows*) can be clearly discerned based on Arc expression since it is heavily myelinated and contains Arc-positive neurons. (**b**) Area X can be clearly delineated based on Arc expression and appears slightly darker than the surrounding striatum. (**c**) Arc-positive neurons are observed in striatal areas GP and LSt. (**d**) A negative control at this level (performed by omitting the primary antibody) from the Trained group and a high-power image on the right [of the rectangular outline in (**d**)] demonstrates the lack of label in LSt. Scale bar for *inset*, 200 µm. (**e**) In the arcopallium, the song control nucleus RA and the adjacent dorsal intermediate arcopallium (AId) can be clearly demarcated based on patterns of Arc staining across different groups of house crows. Scale bar, 1mm. D, Dorsal; M, Medial.
