## Supplementary Figure 3 for "Changes in the Dopaminergic circuitry and Adult Neurogenesis linked to Reinforcement Learning in Corvids"

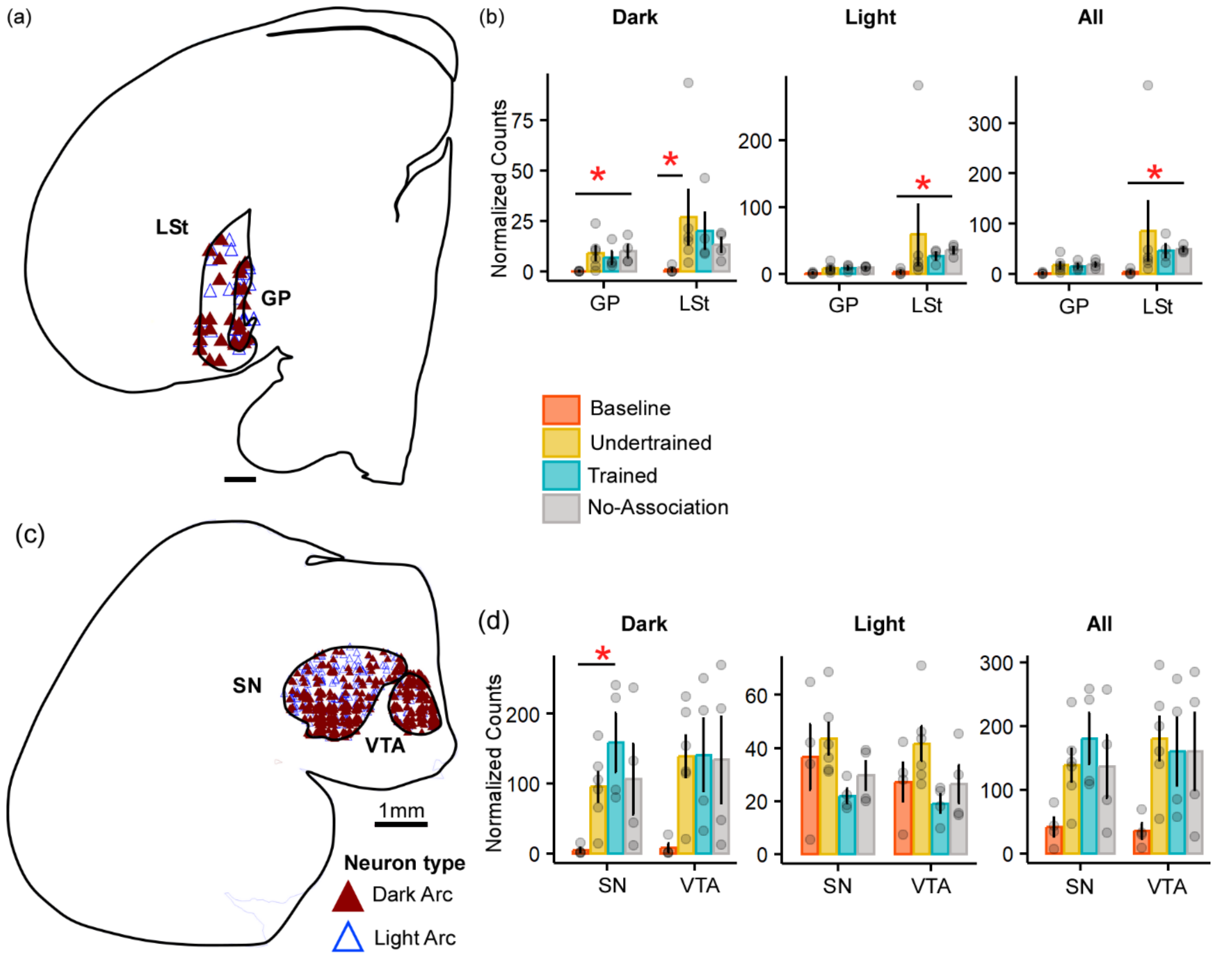


**Supplementary Figure 3.** Expression of Arc in GP, LSt, SN, and VTA.

(**a**) A schematic showing neural activity in GP and LSt in a house crow from the Trained group. (**b**) There were no significant differences in the number of Arc-positive (mean ± SEM) neurons (Dark, light, and total Arc population) in GP and LSt across the three training groups (Undertrained, Trained, and No-Association). However, the lowest number of Arc-labeled neurons was present in the Baseline group. (**c**) A schematic showing Arc expression in SN and VTA in the house crow midbrain from the Undertrained group. (**d**) The number of Arc-positive neurons (mean ± SEM; Dark, light, and total Arc population) in SN and VTA were similar in Undertrained, Trained, and No-Association groups following the visual discrimination task, suggesting that these groups were equally motivated to obtain the food reward. There were significantly fewer Arc-positive neurons in the Baseline group since they were not trained to associate shapes used for visual discrimination with a reward. Scale bar, 1mm. *, $P < 0.05$.
