## Supplementary Figure 4 for "Changes in the Dopaminergic circuitry and Adult Neurogenesis linked to Reinforcement Learning in Corvids"

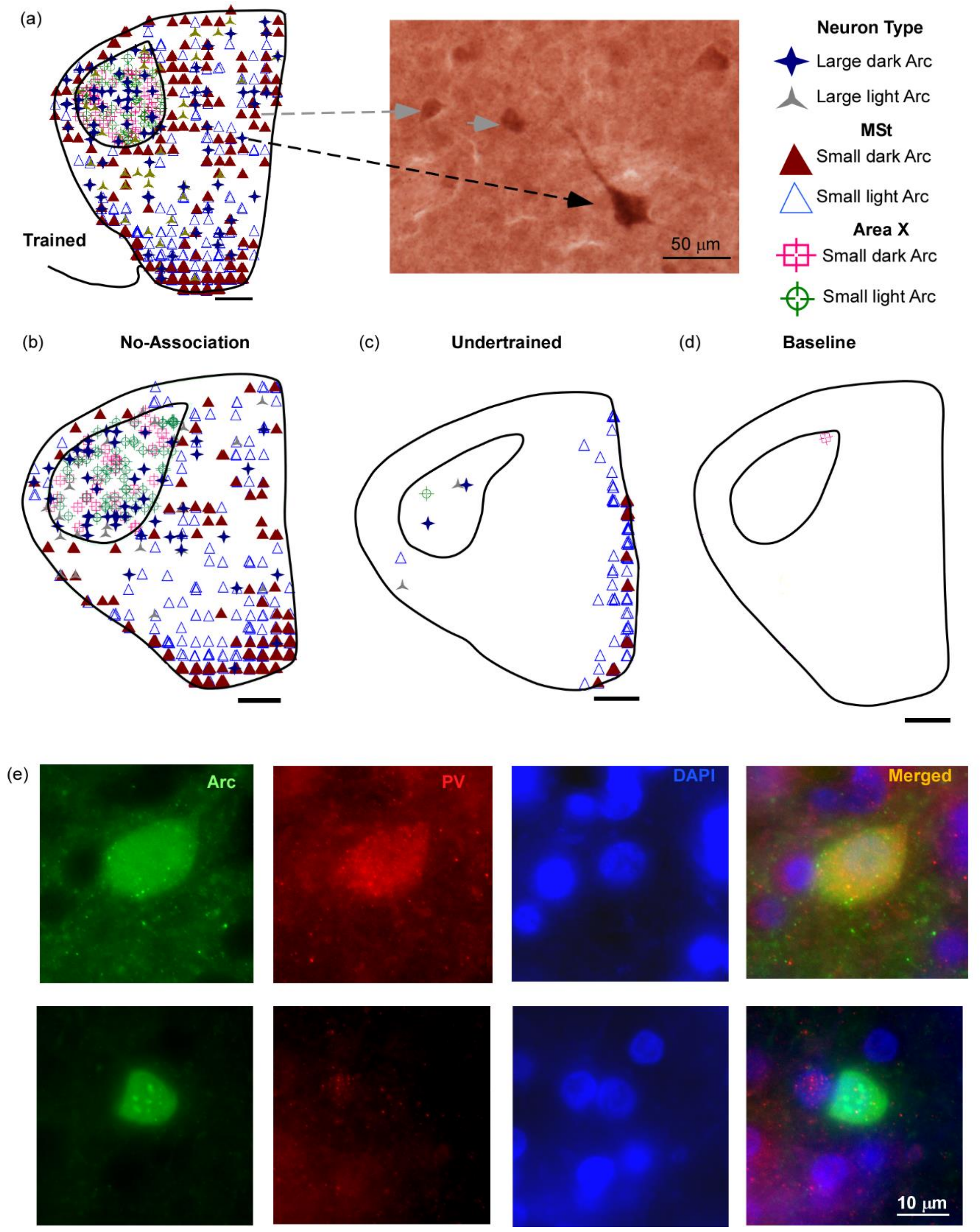


**Supplementary Figure 4.** Arc expression in the anterior striatum.

(**a**) A schematic of Area X and MSt showing Arc expression in a Trained crow and a high magnification image in the right panel showing large (*black arrow*) and small (*gray arrows*) Arc-positive neurons. A comparison of Arc expression in MSt and Area X in the (**b**) No-Association, (**c**) Undertrained, and (**d**) Baseline groups are shown in schematics. (**e**) Both large and small neurons were labeled for Arc. Double-labeling for parvalbumin (PV, first column) and Arc (second column, shown here for a house crow from the No-Association group) was performed to determine the identity of Arc-labeled neurons. The upper row demonstrates a large PV-labeled neuron which is also positive for Arc, suggesting that it may be a striatal interneuron or a pallidal neuron (Reiner, Laverghetta, et al., 2004). The lower row demonstrates an Arc-positive neuron not labeled with PV, which is typical of non-GABAergic interneurons. Scale bar, 1mm, and 10μm.
