## Supplementary Figure 5 for "Changes in the Dopaminergic circuitry and Adult Neurogenesis linked to Reinforcement Learning in Corvids"

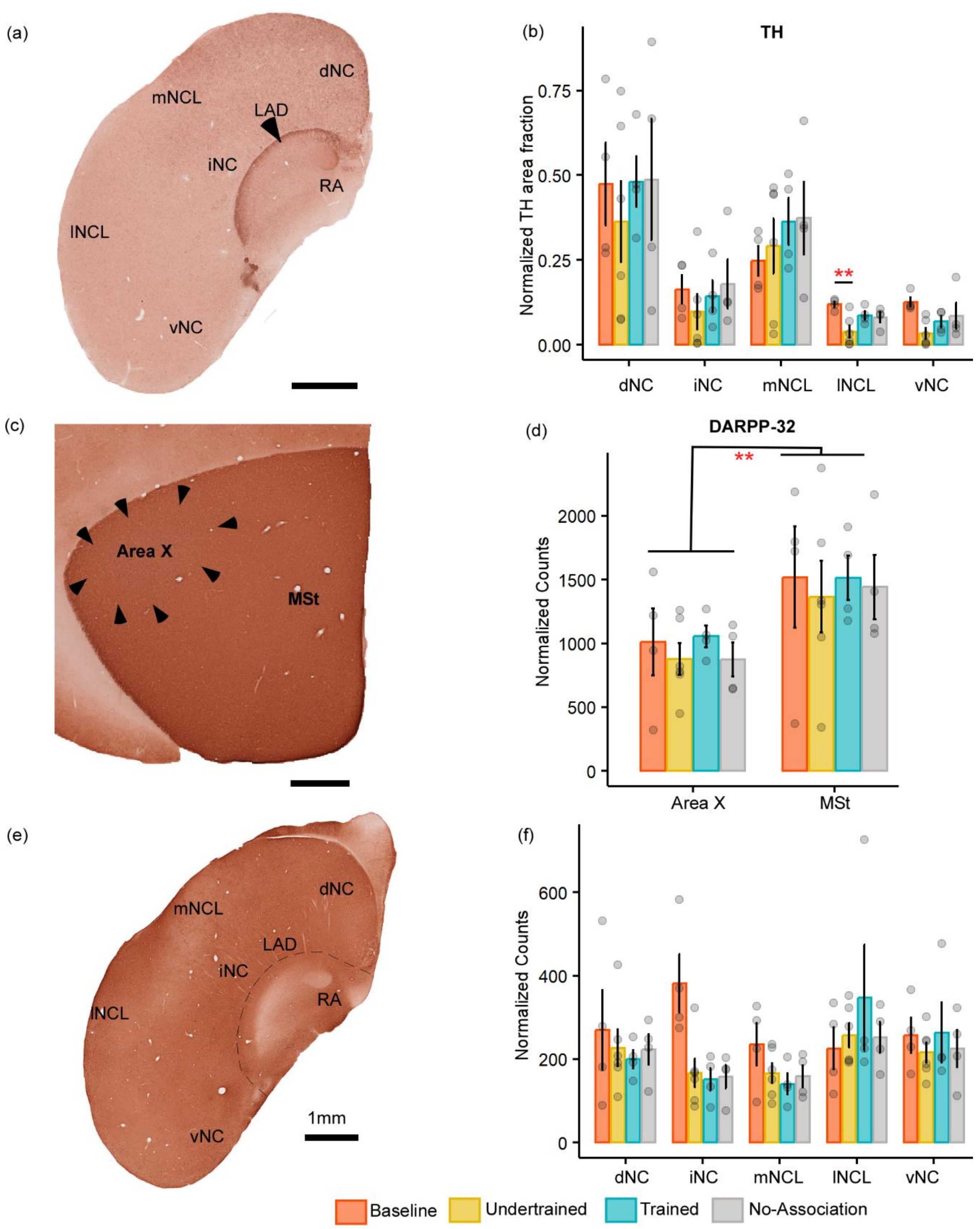


**Supplementary Figure 5.** Catecholaminergic circuitry in different groups of house crows.

(**a**) The locations of different subdivisions of the caudal nidopallium in house crows demonstrated in a coronal section from a Trained bird stained for TH (Sen, Parishar et al., 2019). (**b**) Quantification of the area covered by TH profiles revealed that there were no significant differences in staining for TH in dNC, iNC, mNCL, and vNC across different groups of house crow. In lNCL, TH expression was higher in Baseline controls compared to that in Undertrained birds. (**c**) A DARPP-32-stained coronal section demonstrates that Area X can be discerned from the surrounding MSt, since staining intensity is slightly lower and it has fewer DARPP-positive cells. (**d**) There were no significant differences in overall counts (mean ± SEM) of DARPP-positive neurons in Area X or MSt in any of the experimental groups. (**e**) A DARPP-32-stained section at the level of the caudal nidopallium demonstrating that RA and AId were characteristically devoid of label (cf. Sen, Parishar et al., 2019). (**f**) There were no significant differences in the number of DARPP-32 neurons (mean ± SEM) in any of the subdivisions of NC in the four experimental groups of house crows. Scale bar, 1mm. **, $P < 0.01$.
