## Supplementary Figure 6 for "Changes in the Dopaminergic circuitry and Adult Neurogenesis linked to Reinforcement Learning in Corvids"

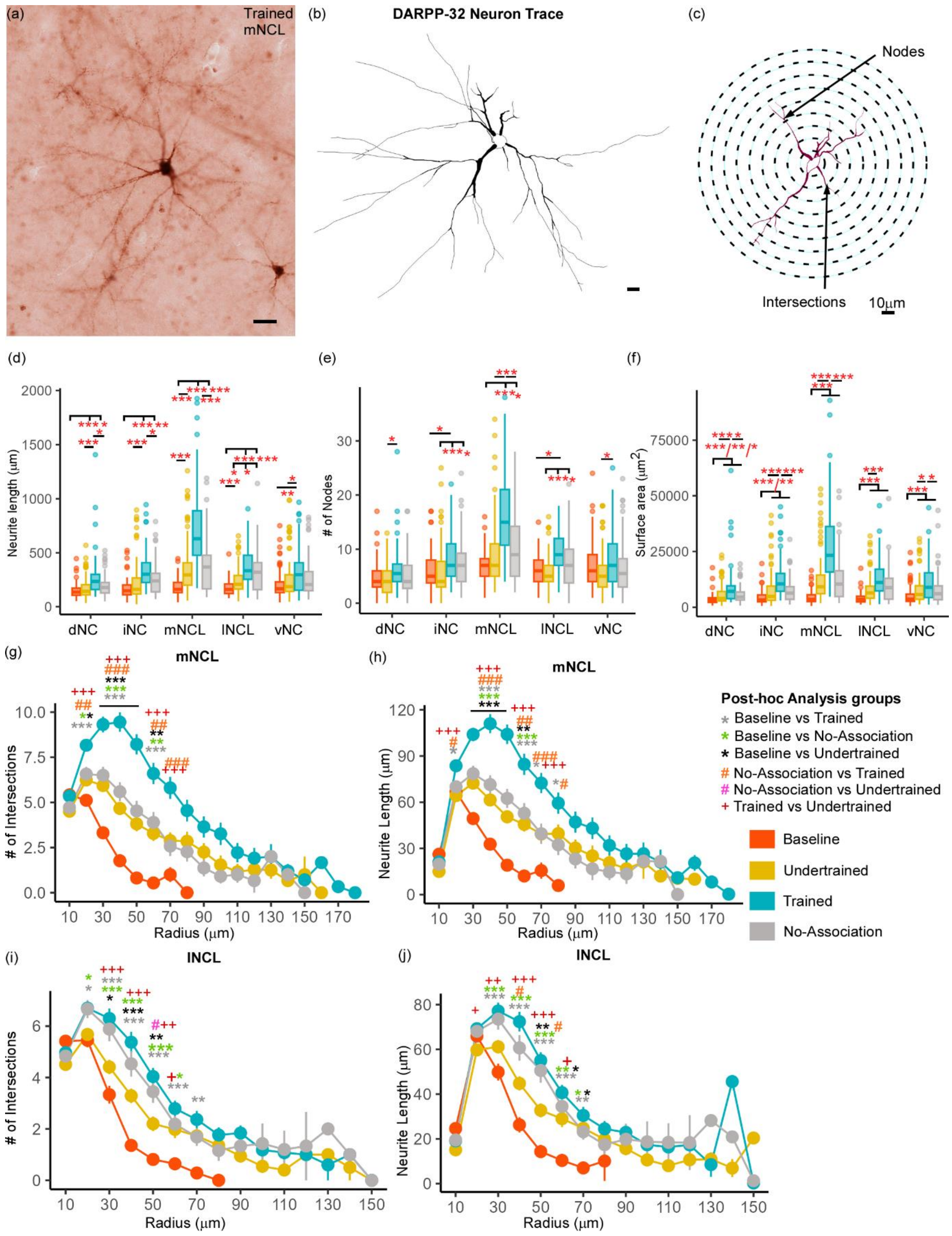


**Supplementary Figure 6.** A comparison of DARPP-32 neurons reconstructed in three dimensions.

(**a**) An example of a DARPP-32-stained neuron from the mNCL region of the Trained group and its (**b**) three-dimensional reconstruction. (**c**) The contour of a DARPP-32 neuron superimposed on concentric Sholl shells. Arrowheads indicate the nodes and point of intersections with the Sholl radii. Scale bar, 10μm. We observed a statistically significant increase in the (**d**) neurite length, and (**e**) number of nodes, which indicate an increase in neurite branching in Trained house crows compared to those in Undertrained, No-Association and Baseline birds in all five subdivisions of NC. (**f**) An increase in the neurite field (measured by 3D surface area) was seen in the Trained group compared to the Undertrained, No-Association and Baseline groups in different divisions of NC. A Sholl analysis demonstrated an increase in the number of intersections and neurite length (mean ± SEM) between the Sholl radii 20-70 μm in the Trained group compared to that in Baseline, Undertrained and No-Association groups in (**g** and **h**) mNCL, (**i** and **j**) lNCL. */#/+ P<0.05; **/##/++ P<0.01; ***/###/+++ P<0.001.
