## Supplementary Figure 7 for "Changes in the Dopaminergic circuitry and Adult Neurogenesis linked to Reinforcement Learning in Corvids"

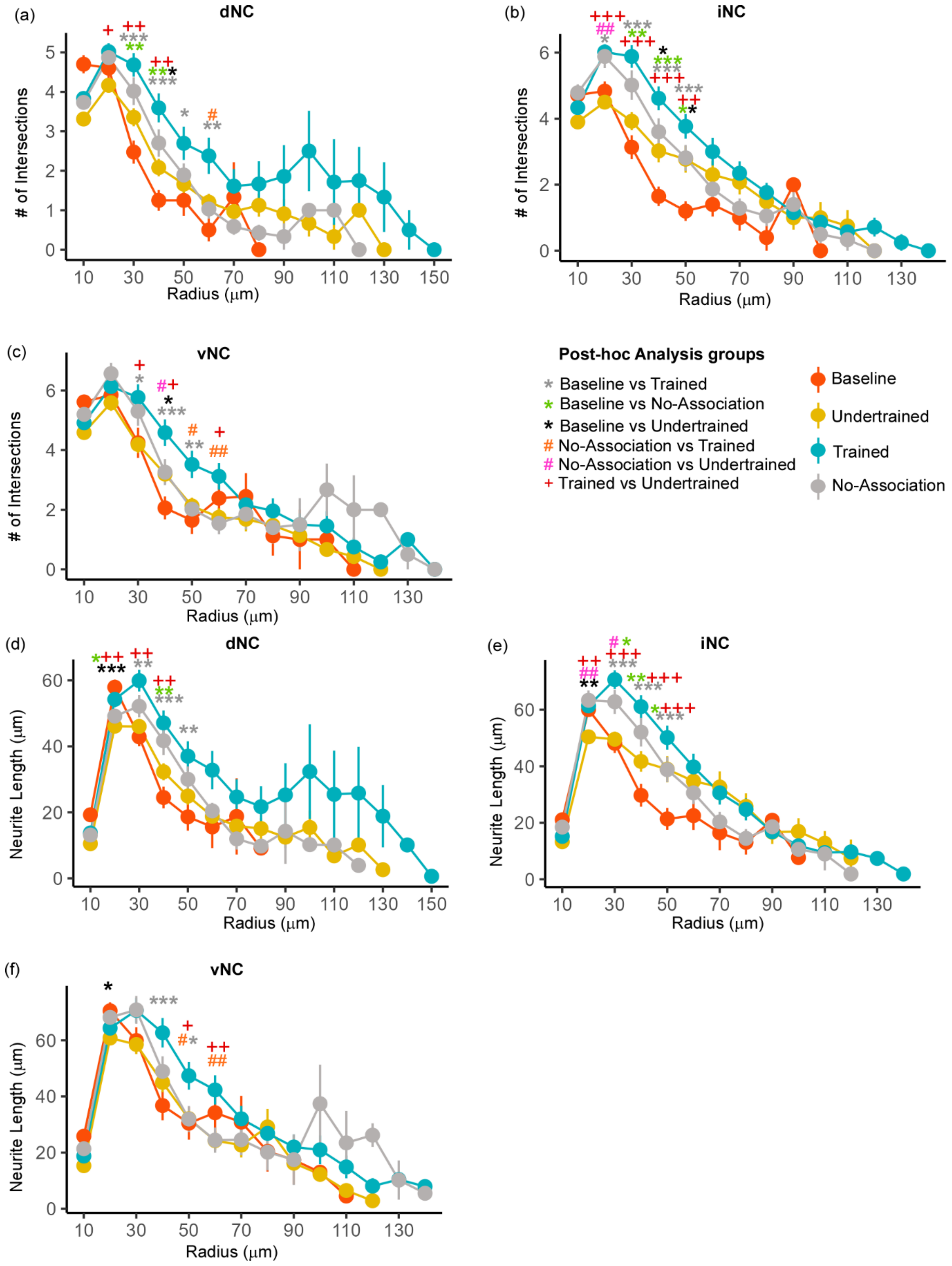


**Supplementary Figure 7.** Sholl analysis of neurite branching and neurite length in DARPP-32 neurons of NC.

An increase in the number of intersections (mean ± SEM) was observed between the Sholl radii 20-70 µm in the Trained group in (**a**) dNC, (**b**) iNC, and (**c**) vNC. In the three subdivisions of NC, the highest number of intersections made by DARPP-32-labeled neurites with Sholl radii were in Trained birds versus that in Undertrained, No-Association and Baseline house crows. At a few points, the number of intersections was significantly higher in the No-Association and Undertrained groups versus that in Baseline controls.  Similar changes were observed for neurite lengths in (**d**) dNC, (**e**) iNC, and (**f**) vNC. */#/+ $P<0.05$; **/##/++ $P<0.01$; ***/###/+++ $P<0.001$.
