## Supplementary Figure 8 for "Changes in the Dopaminergic circuitry and Adult Neurogenesis linked to Reinforcement Learning in Corvids"

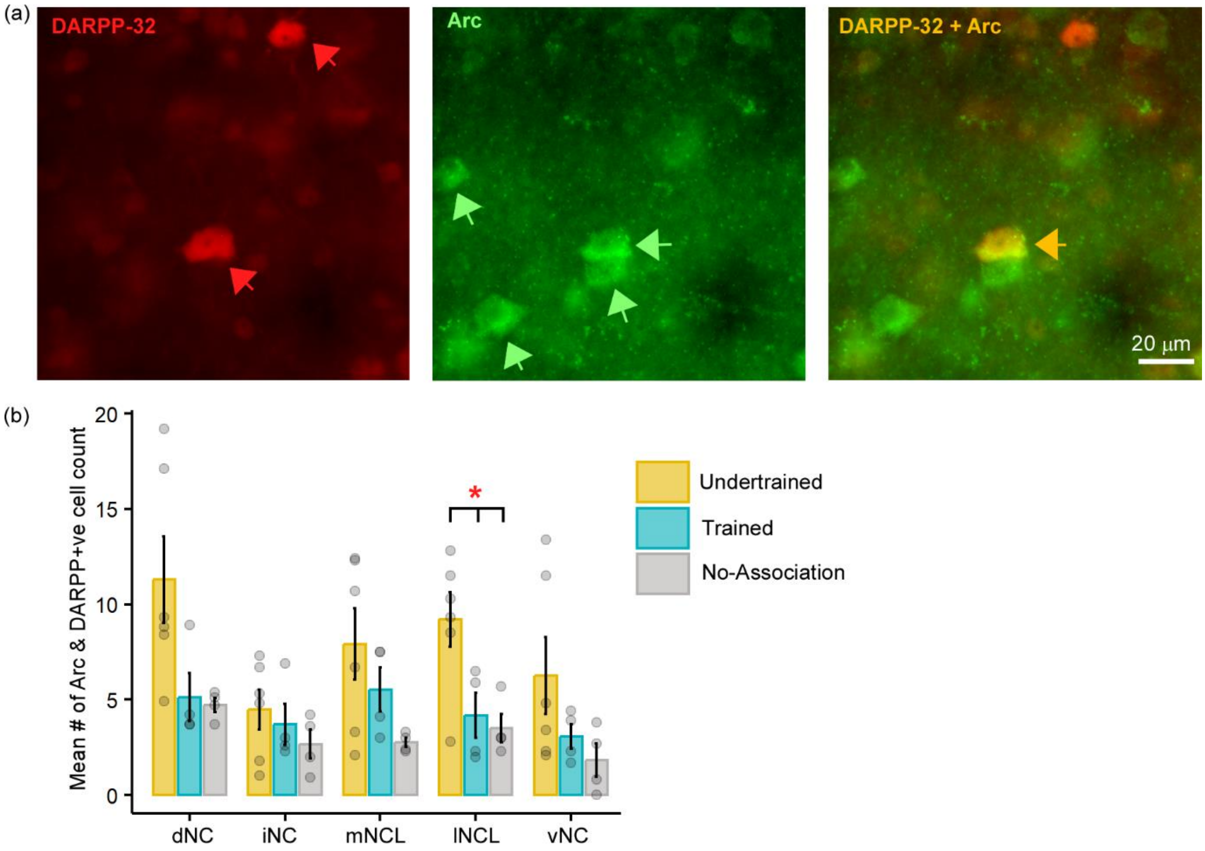


**Supplementary Figure 8.** Double label for Arc and DARPP-32 in NC.

(**a**) Representative images of Arc and DARPP-32 double-label from the lNCL region of an Undertrained bird demonstrating neurons positive for DARPP-32 (*red arrows*), Arc (*green arrows*), and for both (*orange arrows*). (**b**) Counts of neurons double-labeled for Arc and DARPP-32 (mean ± SEM) in NC demonstrated a significantly high number of double-labeled neurons in lNCL of Undertrained birds versus those in the Trained and No-Association groups. Scale bar, 20µm. *, $P < 0.05$.
