## Supplementary Figure 9 for "Changes in the Dopaminergic circuitry and Adult Neurogenesis linked to Reinforcement Learning in Corvids"

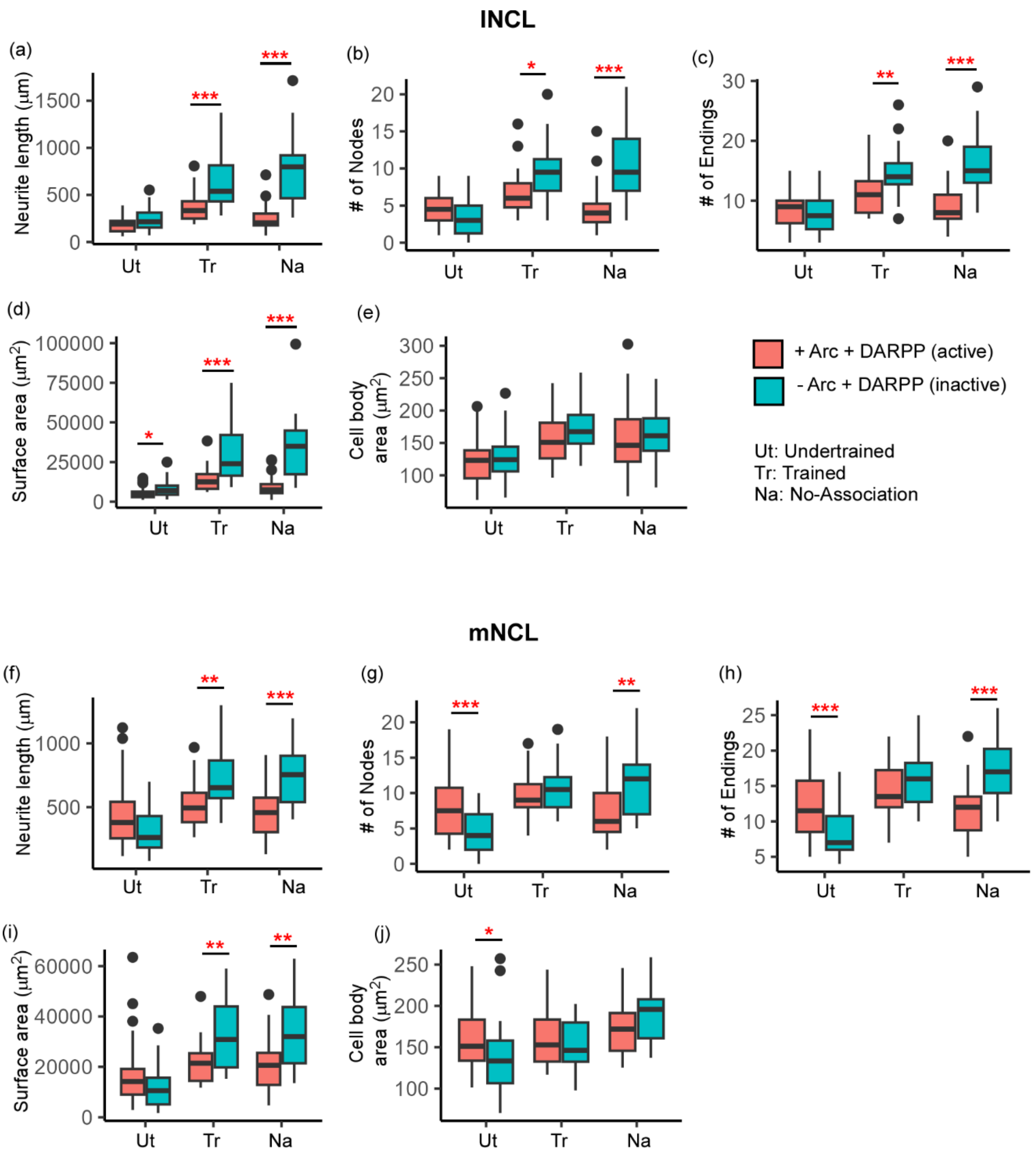


**Supplementary Figure 9.** A comparison of active and inactive DARPP-32-positive neurons in NCL.

In lNCL, the (**a**) neurite length, the number of (**b**) nodes and (**c**) endings were significantly greater in inactive DARPP-32-positive neurons (blue boxplots) compared to active neurons (orange boxplots) of Trained and No-Association birds. (**d**) In all three groups, the area of the neurite field was also higher in inactive versus active neurons. However, (**e**) the area of somata was similar across both sets of neurons in all groups. (**f**) The neurite length of the inactive neurons was significantly greater than that of active neurons in mNCL of the Trained and No-Association groups. However, active neurons in the Undertrained group had significantly more (**g**) nodes and (**h**) endings. In the No-Association group, inactive mNCL neurons were significantly more branched versus active ones. (**i**) The area of neurite fields was significantly greater in inactive mNCL neurons in Trained and No-Association birds versus that in active neurons, which was similar to that in lNCL. (**j**) In mNCL, active neurons of the Undertrained group had larger somata compared to inactive ones. * $P<0.05$; ** $P<0.01$; *** $P<0.001$.
