## Supplementary Figure 10 for "Changes in the Dopaminergic circuitry and Adult Neurogenesis linked to Reinforcement Learning in Corvids"

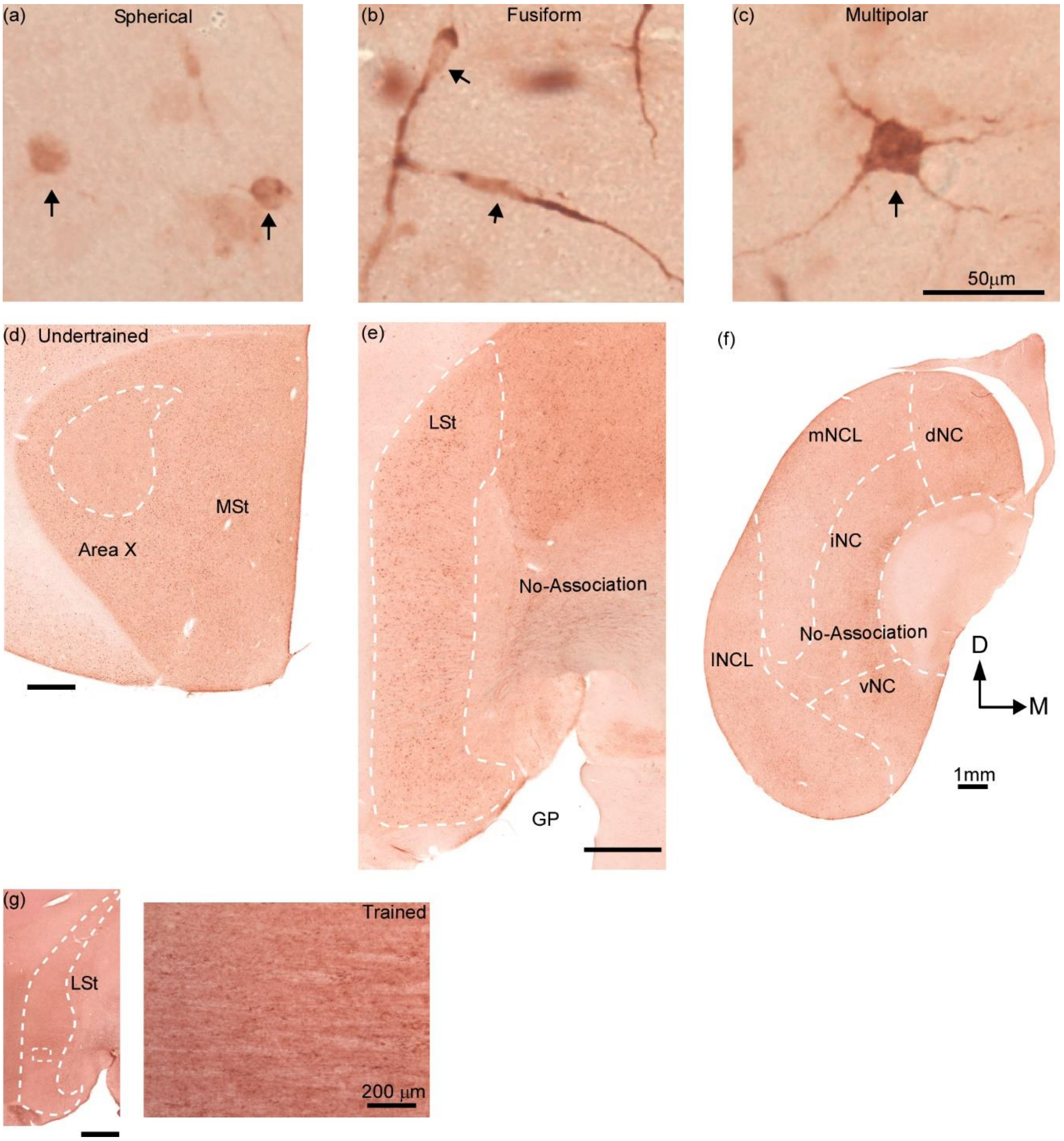


**Supplementary Figure 10.** Doublecortin (DCX) expression in different regions of the house crow brain.

High magnification images of DCX neurons demonstrating differences in their morphology including (**a**) spherical, (**b**) fusiform and (**c**) multipolar neurons (*arrows*). Scale bar, 50 µm. (**d**) In the coronal plane, Area X can be delineated from the surrounding MSt due to lower levels of staining for DCX. (**e**) The boundaries of LSt were clearly demarcated due to the presence of a larger number of DCX-positive neurons in this region compared to that in the adjoining GP. (**f**) Staining for DCX demonstrated that there was a higher density of immature neurons in NC compared to that in the arcopallium. Scale bar, 1mm. (**g**) An image of a negative control was acquired by staining a section at the level of LSt from a Trained bird by following all steps for immunohistochemistry but omitting incubation in the primary antibody solution against DCX. The *inset* on the right demonstrates the lack of staining within LSt at high power. Scale bars, 1mm; 200µm.
