## Supplementary Figure 11 for "Changes in the Dopaminergic circuitry and Adult Neurogenesis linked to Reinforcement Learning in Corvids"

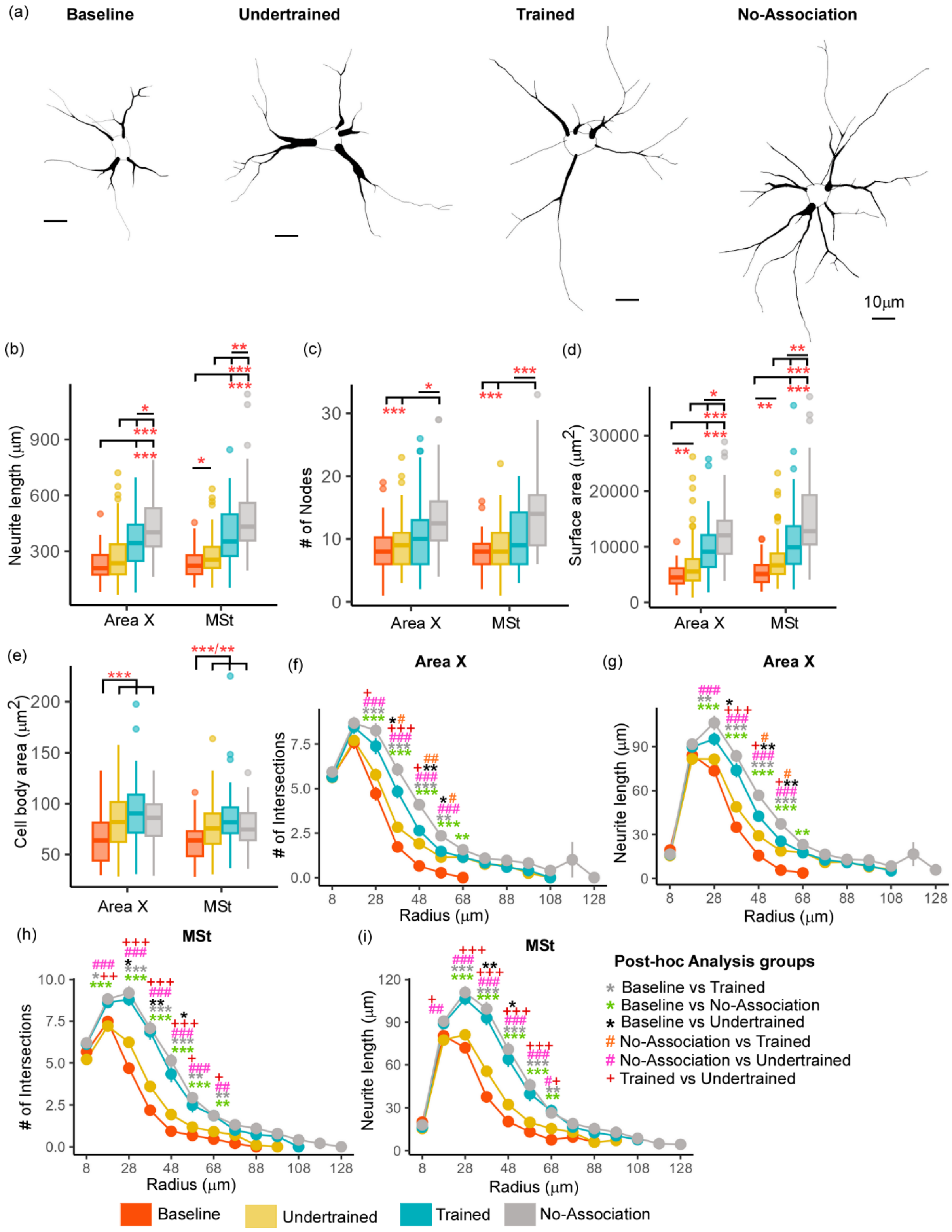


**Supplementary Figure 11.** A comparison of structural changes in DCX-positive neurons in Area X and MSt.

(**a**) Reconstructions of the three-dimensional structure of DCX-positive neurons demonstrating that they were the most complex in terms of the number and branching of dendrites in the No-Association group. Scale bar, 10μm. Comparisons of (**b**) neurite length, (**c**) number of nodes, (**d**) neurite field area measured by performing a convex hull analysis, and (**e**) area of somata were analyzed to study changes in the structure of DCX-positive neurons induced by learning and decision-making. The neurite length of these neurons was significantly greater in Trained and No-Association birds compared to that in Baseline and Undertrained birds. Neurite length of DCX-labeled neurons was the greatest in the No-Association group, followed by that in the Trained group. Similar results were observed for the area of the neurite field. For the number of nodes, significant differences were only observed when comparisons were made between the No-Association versus Trained, Baseline, and Undertrained groups. Somata of DCX-positive neurons in the Trained, Undertrained, and No-Association groups were larger in area compared to those in the Baseline group. The (**f**) number of intersections and (**g**) neurite length of DCX-positive neurons was compared across different behavioral groups in Area X. We found that neurite branching of DCX-positive neurons in No-Association and Trained birds was significantly greater compared to that in Undertrained and Baseline control groups. The largest changes were observed in these parameters in the No-Association group followed by those in the Trained and Undertrained groups. (**h**) The number of intersections and (**i**) neurite length in DCX-positive neurons traced from MSt were significantly higher in the No-Association and Trained groups versus other groups. However, there were no differences in these parameters when the Trained and No-Association groups were compared. */+/#, $P<0.05$; **/++/##, $P<0.01$; ***/+++/###, $P<0.001$.
