## Supplementary Figure 12 for "Changes in the Dopaminergic circuitry and Adult Neurogenesis linked to Reinforcement Learning in Corvids"

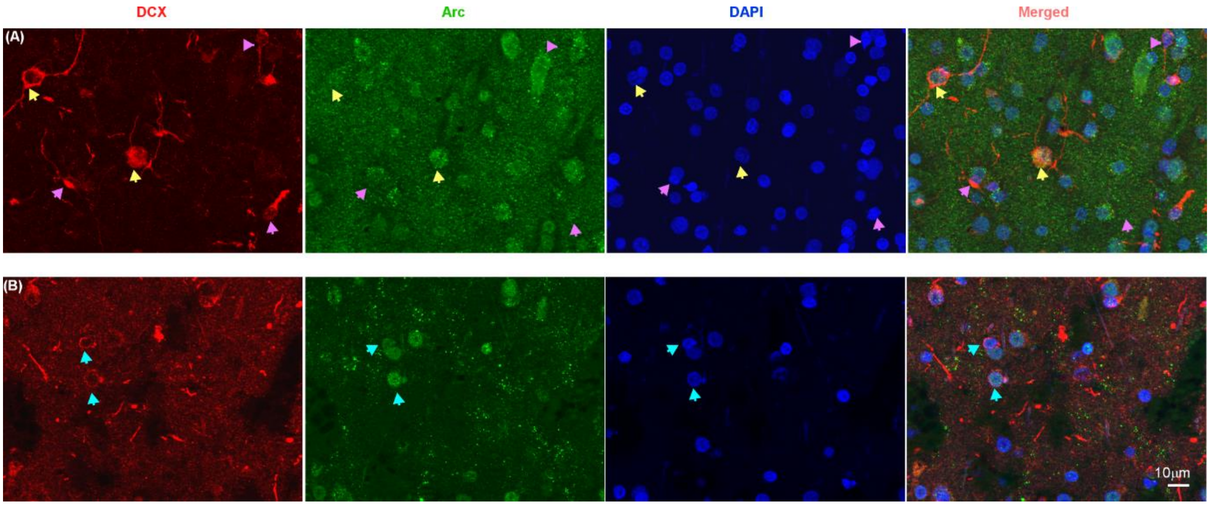


**Supplementary Figure 12.** Double-labeled Arc and DCX neurons in MSt.

Selected images are shown from the same z-stack but at different planes from MSt in an Undertrained bird. (**a**) In the upper panel, multipolar and fusiform neurons are labelled with DCX (magenta), Arc (green), the nuclear marker DAPI (blue) and the merged image (last column). Neurons labeled with DCX are indicated by *yellow arrows*, whereas Arc-positive neurons are indicated by *pink arrows*. The *yellow arrow* in the center of the image demonstrates a DCX- and Arc- co-labeled multipolar neuron, whereas in the image on the top left, a *yellow arrow* indicates an inactive (Arc-negative) DCX-positive multipolar neuron. The merged image clearly demonstrates that fusiform cells (*pink arrows*) are devoid of the Arc label. (**b**) In the bottom panel, *blue arrows* in images from left to right showing staining for DCX (magenta), Arc (green), and DAPI (blue), and in the merged image (last column) indicate the presence of active spherical neurons which are devoid of processes in MSt. Scale bar: 10 µm.
