## Supplementary Figure 13 for "Changes in the Dopaminergic circuitry and Adult Neurogenesis linked to Reinforcement Learning in Corvids"

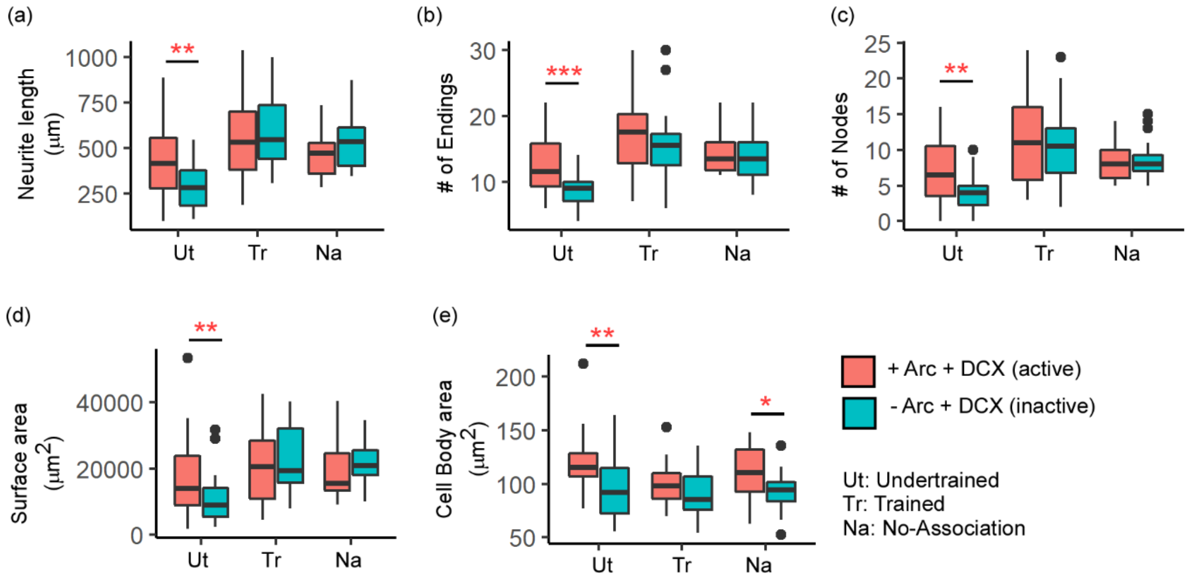


**Supplementary Figure 13.** A comparison of the complexity of active and inactive DCX neurons in MSt.

The (**a**) length of neurites was significantly greater in active DCX-positive neurons versus inactive ones in the Undertrained group in MSt. In the same group, the (**b**) number of endings and (**c**) number of nodes was also greater in active DCX-labeled neurons compared to those which were inactive. (**d**) Active DCX-positive neurons in Undertrained birds additionally demonstrated an expansion in the area of neurites versus inactive Arc-negative DCX-positive neurons. (**e**) The area of somata of active DCX-labeled neurons in the Undertrained and No-Association birds, compared to this measure for inactive neurons. *, P < 0.05; **, P <0.01; ***, P < 0.001.
