## Supplementary Figure 14 for "Changes in the Dopaminergic circuitry and Adult Neurogenesis linked to Reinforcement Learning in Corvids"

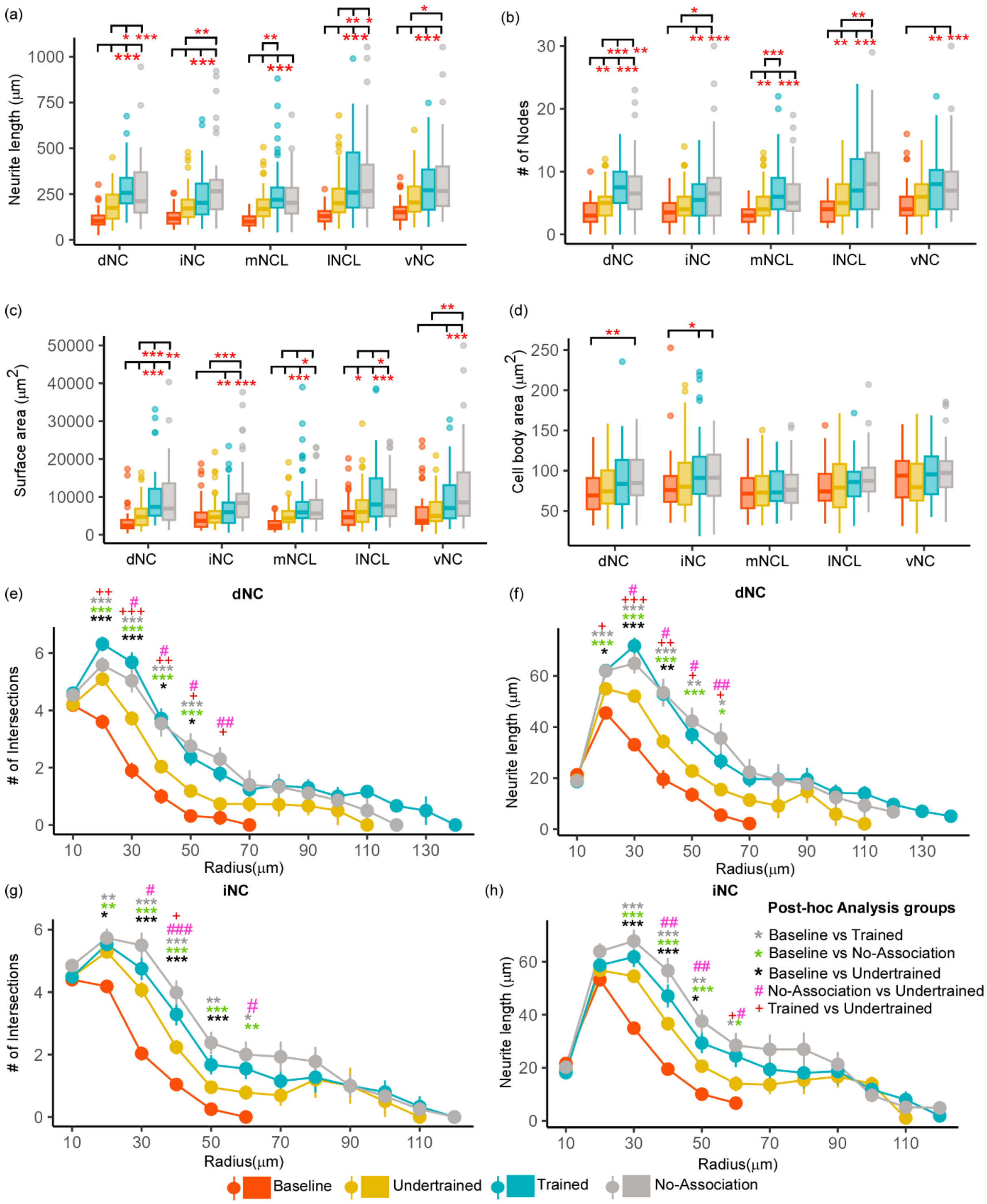


**Supplementary Figure 14.** Structural changes in multipolar DCX-positive neurons in NC.

The (**a**) neurite length, (**b**) number of nodes and (**c**) neurite field area was significantly greater in Undertrained, Trained and No-Association groups compared to that in Baseline controls in all subdivisions of NC. Furthermore, these parameters were significantly greater in NC of Trained and No-Association birds versus that in Undertrained birds. (**d**) The area of somata was significantly greater in dNC and iNC of No-Association birds versus that in Baseline controls. A Sholl analysis demonstrated that the number of intersections and neurite length were significantly greater in Trained and No-Association birds versus that in Baseline and Undertrained crows in (**e** and **f**) dNC and (**g** and **h**) iNC. */+/#, P<0.05; **/++/##, P<0.01; ***/+++/###, P<0.001.
