## Supplementary Figure 15 for "Changes in the Dopaminergic circuitry and Adult Neurogenesis linked to Reinforcement Learning in Corvids"

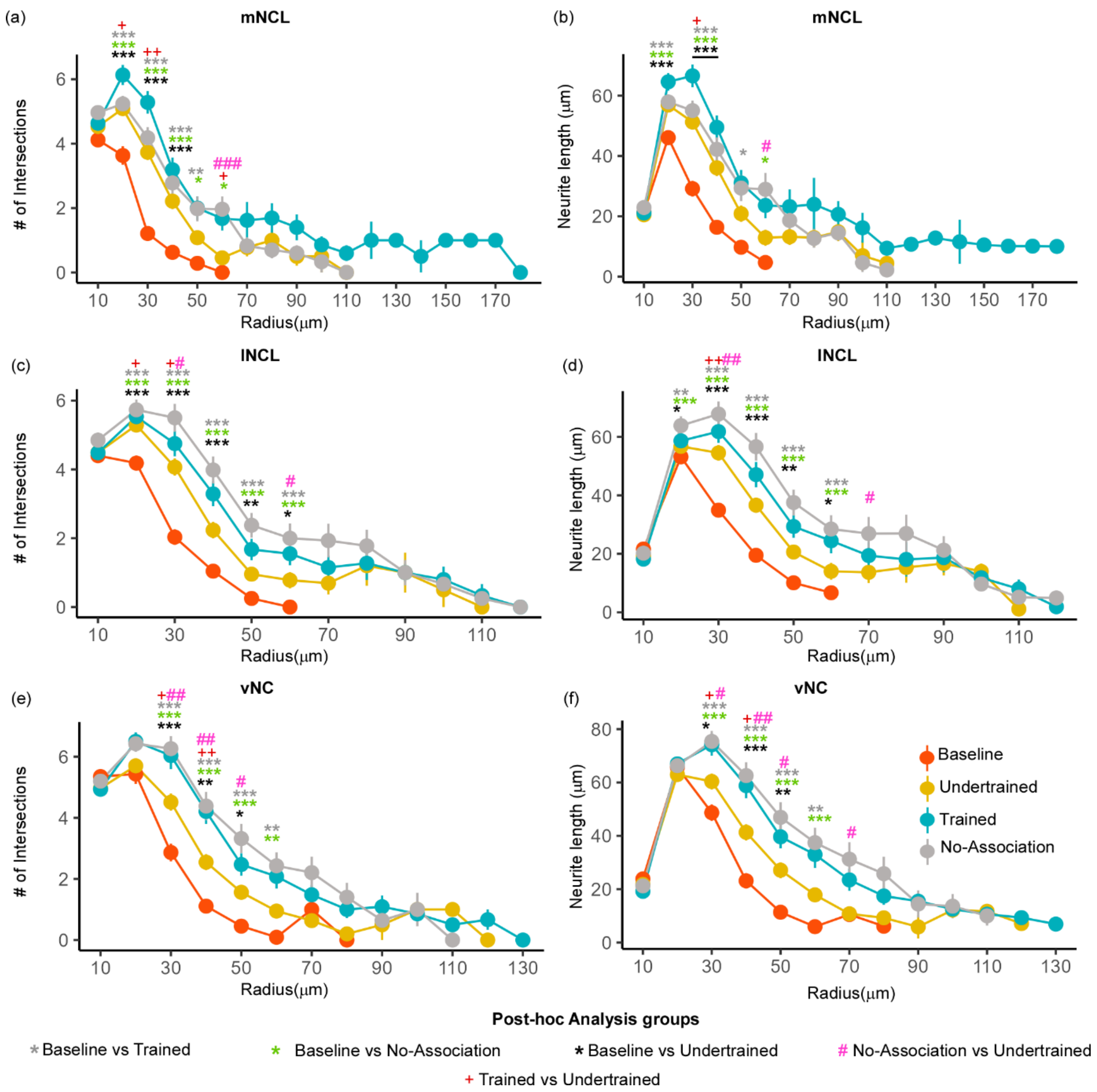


**Supplementary Figure 15.** A Sholl analysis of DCX-positive multipolar neurons in different subdivisions of NC.

The number of intersections and neurite length were significantly greater in DCX-labeled neurons in the Trained and No-Association groups versus Baseline and Undertrained groups in (**a** and **b**) mNCL, (**c** and **d**) lNCL, and (**e** and **f**) vNC. */+/#, P<0.05; **/++/##, P<0.01; ***/+++/###, P<0.001.
