## Supplementary Figure 16 for "Changes in the Dopaminergic circuitry and Adult Neurogenesis linked to Reinforcement Learning in Corvids"

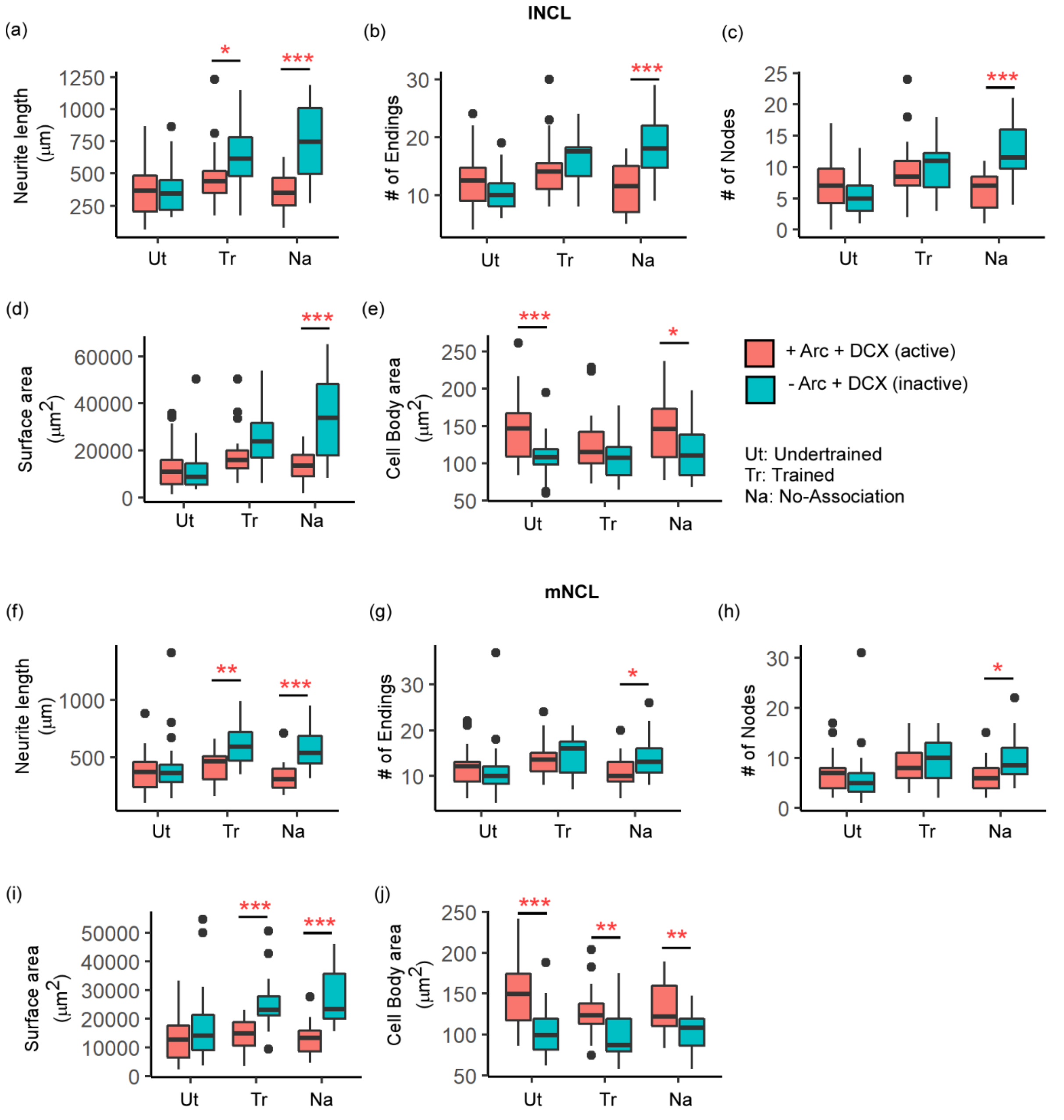


**Supplementary Figure 16.** Comparison of active and inactive DCX-positive neurons in mNCL and lNCL.

The (**a**) length of neurites of inactive DCX-positive neurons was significantly greater in the Trained and No-Association groups compared to active DCX-positive neurons in lNCL. The (**b**) number of endings, (**c**) number of nodes and (**d**) the area of the neurite field was significantly greater only in inactive DCX-positive neurons of the No-Association group. The (**e**) area of the somata was significantly greater in the active versus inactive population of DCX-labeled neurons in the Undertrained and No-Association groups. Similar changes in (**f**) neurite length, (**g**) number of endings and (**h**) number of nodes were observed in the reconstructed active and inactive DCX neurons in mNCL. (**i**) The area of the neurite field was greater in inactive DCX-labeled neurons in mNCL of the Trained and No-Association groups compared to active DCX-positive neurons. (**j**) In all three experimental groups, the size of the somata was greater in active DCX-labeled neurons compared to inactive neurons. *, $P < 0.05$; **, $P < 0.01$; ***, $P < 0.001$.
