## Supplementary material for "Changes in the Dopaminergic circuitry and Adult Neurogenesis linked to Reinforcement Learning in Corvids": Table S1

**Table S1**. Statistical details of Arc expression.

| Area | Neuronal type | Test | P value | F/χ2 value | Degrees of Freedom (df) | Post-hoc P value (Tukey's /Dunn's) |
|---|---|---|---|---|---|---|
| dNC | Dark | Kruskal-Wallis test | 0.03205 | 8.8017 | 3 | Bl vs Tr: 0.0216 |
| | Light | Kruskal-Wallis test | 0.02141 | 9.6883 | 3 | Bl vs Tr: 0.0194 |
| | All | Kruskal-Wallis test | 0.01973 | 9.8674 | 3 | Bl vs Tr: 0.0156 |
| iNC | Dark | Kruskal-Wallis test | 0.03273 | 8.7548 | 3 | Bl vs Tr: 0.0223 |
| | Light | Kruskal-Wallis test | 0.01468 | 10.512 | 3 | Bl vs Na: 0.0296 |
| | | | | | | Bl vs Tr: 0.0303 |
| | All | Kruskal-Wallis test | 0.01468 | 10.512 | 3 | Bl vs Na: 0.0296 |
| | | | | | | Bl vs Tr: 0.0303 |
| mNCL | Dark | Kruskal-Wallis test | 0.01973 | 9.8674 | 3 | Bl vs Tr: 0.0156 |
| | Light | Kruskal-Wallis test | 0.01962 | 9.8793 | 3 | Bl vs Na: 0.0371 |
| | | | | | | Bl vs Tr: 0.0364 |
| | All | Kruskal-Wallis test | 0.0222 | 9.6077 | 3 | Bl vs Na: 0.0452 |
| | | | | | | Bl vs Tr: 0.0364 |
| lNCL | Dark | Kruskal-Wallis test | 0.0203 | 9.8047 | 3 | Bl vs Tr: 0.0124 |
| | Light | Kruskal-Wallis test | 0.0193 | 9.9152 | 3 | Bl vs Na: 0.0296 |
| | | | | | | Bl vs Tr: 0.0452 |
| | All | Kruskal-Wallis test | 0.0222 | 9.6077 | 3 | Bl vs Na: 0.0452 |
| | | | | | | Bl vs Tr: 0.0364 |
| vNC | Light | Kruskal-Wallis test | 0.0201 | 9.8183 | 3 | Bl vs Na: 0.0210 |
| | All | Kruskal-Wallis test | 0.0306 | 8.9039 | 3 | Bl vs Na: 0.0323 |
| AId | Dark | Kruskal-Wallis test | 0.0074 | 11.987 | 3 | Bl vs Tr: 0.0046 |
| | Light | ANOVA | 0.001 | 9.573 | 3,14 | Bl vs Na: 0.0069 |
| | | | | | | Bl vs Tr: 0.0022 |
| | | | | | | Ut vs Tr: 0.0179 |
| | All | ANOVA | 0.0018 | 8.501 | 3,14 | Bl vs Tr: 0.0014 |
| | | | | | | Ut vs Tr: 0.0130 |
| RA | Dark | Kruskal-Wallis test | 0.0064 | 12.282 | 3 | Bl vs Tr: 0.0032 |
| | Light | Kruskal-Wallis test | 0.0138 | 10.64 | 3 | Bl vs Tr: 0.0136 |
| | All | Kruskal-Wallis test | 0.0118 | 10.972 | 3 | Bl vs Tr: 0.0108 |
| GP | Dark | Kruskal-Wallis test | 0.02195 | 9.6333 | 3 | Bl vs Na: 0.0259 |
| LSt | Dark | Kruskal-Wallis test | 0.02872 | 9.0432 | 3 | Bl vs Ut: 0.0296 |
| | Light | Kruskal-Wallis test | 0.01489 | 10.482 | 3 | Bl vs Na: 0.0110 |
| | All | Kruskal-Wallis test | 0.02392 | 9.4454 | 3 | Bl vs Na: 0.0323 |
| SN | Dark | Kruskal-Wallis test | 0.03965 | 8.3304 | 3 | Bl vs Tr: 0.0397 |
| LMAN | Dark | Kruskal-Wallis test | 0.01167 | 11.01 | 3 | Bl vs Na: 0.0054 |
| | Light | ANOVA | 0.0472 | 3.415 | 3, 14 | |
| | Total | Kruskal-Wallis test | 0.03438 | 8.6462 | 3 | Bl vs Na: 0.0214 |
| Area X | Large-D | Kruskal-Wallis test | 0.00693 | 12.136 | 3 | Bl vs Na: 0.0303 |
| | | | | | | Bl vs Tr: 0.0124 |
| | Dark | Kruskal-Wallis test | 0.007814 | 11.878 | 3 | Bl vs Tr: 0.0269 |
| | Large-L | Kruskal-Wallis test | 0.01654 | 10.253 | 3 | Bl vs Na: 0.0452 |
| | | | | | | Bl vs Tr: 0.0240 |
| | Light | Kruskal-Wallis test | 0.005007 | 12.835 | 3 | Bl vs Na: 0.0317 |
| | | | | | | Bl vs Tr: 0.0249 |
| | All | Kruskal-Wallis test | 0.008757 | 11.632 | 3 | Bl vs Na: 0.0247 |
| | | | | | | Bl vs Tr: 0.0194 |
| MSt | Large -D | Kruskal-Wallis test | 0.01331 | 10.725 | 3 | Bl vs Tr: 0.0412 |
| | Dark | Kruskal-Wallis test | 0.01444 | 10.253 | 3 | Bl vs Na: 0.0240 |
| | | | | | | Bl vs Tr: 0.0371 |
| | Large-L | Kruskal-Wallis test | 0.007448 | 11.981 | 3 | Bl vs Tr: 0.0062 |
| | Light | Kruskal-Wallis test | 0.01751 | 10.128 | 3 | Bl vs Na: 0.0203 |
| | All | Kruskal-Wallis test | 0.01702 | 10.19 | 3 | Bl vs Na: 0.0310 |
