## Supplementary material for "Changes in the Dopaminergic circuitry and Adult Neurogenesis linked to Reinforcement Learning in Corvids": Table S2

**Table S2.** Statistical details of DARPP-32 neuron summary analysis.

| Area | Neuron Morphometry parameters | Kruskal-Wallis P value | χ2 value | df | Post hoc P value (Dunn's) |
|---|---|---|---|---|---|
| dNC | Neurite length | 2.73E-07 | 33.339 | 3 | Bl vs Na: 2.355e-02<br>Bl vs Tr: 4.9580e-07<br>Na vs Tr: 2.7498e-02<br>Tr vs Ut: 2.0628e-05 |
| | Endings | 0.0001449 | 20.331 | 3 | Tr vs Ut: 5.4015e-05 |
| | Node | 3.55E-03 | 13.572 | 3 | Tr vs Ut: 0.0014 |
| | Cell body area | 0.0001712 | 19.982 | 3 | Bl vs Na: 0.0013<br>Bl vs Tr: 0.0242<br>Bl vs Ut: 0.0001 |
| | Neurite field area | 6.49E-08 | 36.295 | 3 | Bl vs Na: 2.5422e-03<br>Bl vs Tr: 1.6625e-08<br>Bl vs Ut: 1.6684e-02<br>Na vs Tr: 2.2899e-02<br>Tr vs Ut: 9.2214e-04 |
| iNC | Neurite length | 2.35E-09 | 43.097 | 3 | Bl vs Na: 4.8596e-03<br>Bl vs Tr: 5.7186e-09<br>Na vs Tr: 1.1865e-02<br>Tr vs Ut: 1.4886e-06 |
| | Endings | 0.0006685 | 17.117 | 3 | Tr vs Ut: 0.00038 |
| | Nodes | 0.0001785 | 19.895 | 3 | Bl vs Tr: 0.0260<br>Na vs Ut: 0.0405<br>Tr vs Ut: 0.0001 |
| | Cell body area | 1.48E-06 | 29.855 | 3 | Bl vs Na: 6.906e-03<br>Bl vs Tr: 8.2909e-04<br>Bl vs Ut: 4.6052e-07 |
| | Neurite field area | 1.97E-12 | 57.537 | 3 | Bl vs Na: 7.4356e-04<br>Bl vs Tr: 5.0211e-13<br>Bl vs Ut: 4.6483e-03<br>Na vs Tr: 5.8146e-04<br>Tr vs Ut: 1.4372e-06 |
| mNCL | Neurite length | <2.2e-16 | 101.58 | 3 | Bl vs Na: 2.9169e-06<br>Bl vs Tr: 5.6585e-23<br>Bl vs Ut: 2.4537e-06<br>Na vs Tr: 1.0405e-06<br>Tr vs Ut: 3.7504e-09 |
| | Endings | 6.98E-13 | 59.65 | 3 | Bl vs Na: 3.7367e-02<br>Bl vs Tr: 1.2931e-12<br>Na vs Tr: 5.1978e-06<br>Tr vs Ut: 1.0520e-08 |
| | Nodes | 9.40E-13 | 59.045 | 3 | Bl vs Na: 1.6916e-02<br>Bl vs Tr: 2.8474e-12<br>Na vs Tr: 3.2131e-05 |

| | | | | | |
|---|---|---|---|---|---|
| | | | | | Tr vs Ut: 4.3118e-09 |
| | Cell body area | 1.12E-11 | 54.004 | 3 | Bl vs Na: 4.2205e-04<br>Bl vs Tr: 1.0789e-03<br>Bl vs Ut: 1.3300e-12<br>Na vs Ut: 4.9192e-03<br>Tr vs Ut: 2.4721e-03 |
| | Neurite field area | <2.2e-16 | 115.18 | 3 | Bl vs Na: 1.9548e-07<br>Bl vs Tr: 4.3815e-26<br>Bl vs Ut: 9.8288e-09<br>Na vs Tr: 2.0009e-07<br>Tr vs Ut: 3.4799e-08 |
| lNCL | Neurite length | 5.80E-11 | 50.652 | 3 | Bl vs Na: 2.5589e-05<br>Bl vs Tr: 5.1715e-11<br>Bl vs Ut: 8.4143e-03<br>Na vs Tr: 4.6662e-02<br>Na vs Ut: 4.4775e-02<br>Tr vs Ut: 2.8331e-05 |
| | Endings | 3.11E-07 | 33.071 | 3 | Bl vs Tr: 4.8049e-05<br>Na vs Tr: 2.73350e-02<br>Na vs Ut: 4.2754e-02<br>Tr vs Ut: 3.7009e-07 |
| | Nodes | 6.00E-08 | 36.457 | 3 | Bl vs Tr: 1.129886e-04<br>Na vs Tr: 1.9903e-02<br>Na vs Ut: 1.6804e-02<br>Tr vs Ut: 3.0047e-08 |
| | Cell body area | 6.24E-05 | 22.094 | 3 | Bl vs Na: 2.7019e-02<br>Bl vs Tr: 1.3456e-02<br>Bl vs Ut: 1.8031e-05 |
| | Neurite field area | 1.11E-12 | 58.715 | 3 | Bl vs Na: 6.8181e-07<br>Bl vs Tr: 5.7865e-13<br>Bl vs Ut: 3.9893e-05<br>Tr vs Ut: 5.5516e-04 |
| vNC | Neurite length | 0.005778 | 12.528 | 3 | Bl vs Tr: 0.0055<br>Tr vs Ut: 0.0356 |
| | Endings | 0.0233 | 9.5029 | 3 | Tr vs Ut: 0.0240 |
| | Node | 0.01441 | 10.552 | 3 | Tr vs Ut: 0.0150 |
| | Cell body area | 1.48E-06 | 29.855 | 3 | Bl vs Na: 4.1177e-02<br>Bl vs Tr: 1.5321e-02<br>Bl vs Ut: 5.4914e-07<br>Na vs Ut: 3.2848e-02 |
| | Neurite field area | 3.09E-05 | 23.556 | | Bl vs Na: 4.9593e-02<br>Bl vs Tr: 7.4806e-06<br>Bl vs Ut: 2.2047e-02<br>Na vs Tr: 3.7804e-02<br>Tr vs Ut: 4.1275e-02 |
