## Supplementary material for "Changes in the Dopaminergic circuitry and Adult Neurogenesis linked to Reinforcement Learning in Corvids": Table S3

**Table S3.** Statistical details of Sholl analysis of DARPP-32 positive neurons.

| Area | Radius (µm) | Intersections (KW P-Value) | χ2 value | df | Intersections (Dunn's value) | Neurite length (KW P-Value) | χ2 value | df | Neurite length (Dunn's P value) |
|---|---|---|---|---|---|---|---|---|---|
| dNC | 20 | 0.0269 | 9.18 | 3 | Tr vs Ut: 0.0322 | 0.00038 | 18.26 | 3 | Bl vs Na: 0.0287<br>Bl vs Ut: 0.0004<br>Tr vs Ut: 0.0357 |
| | 30 | 1.84E-06 | 29.40 | 3 | Bl vs Na: 3.0374e-03<br>Bl vs Tr: 9.4381e-07<br>Tr vs Ut: 2.7109e-03 | 0.00165 | 15.20 | 3 | Bl vs Tr: 0.0032<br>Tr vs Ut: 0.0063 |
| | 40 | 4.75E-07 | 32.20 | 3 | Bl vs Na: 2.2596e-03<br>Bl vs Tr: 2.1773e-07<br>Bl vs Ut: 3.0638e-02<br>Tr vs Ut: 1.1945e-03 | 3.38E-05 | 23.37 | 3 | Bl vs Na: 7.0483e-03<br>Bl vs Tr: 3.2721e-05<br>Tr vs Ut: 5.7953e-03 |
| | 50 | 0.02413 | 9.43 | 3 | Bl vs Tr: 0.0236 | 0.01042 | 11.26 | 3 | Bl vs Tr: 0.0092 |
| | 60 | 0.002 | 14.76 | 3 | Bl vs Tr: 0.0056<br>Na vs Tr: 0.0337 | 0.02791 | 9.11 | 3 | - |
| iNC | 20 | 2.58E-05 | 23.93 | 3 | Bl vs Tr: 1.1176e-02<br>Na vs Ut: 8.6244e-03<br>Tr vs Ut: 3.0370e-05 | 0.00026 | 19.04 | 3 | Bl vs Ut: 0.0043<br>Na vs Ut: 0.0070<br>Tr vs Ut: 0.0017 |
| | 30 | 2.34E-07 | 33.66 | 3 | Bl vs Na: 1.7962e-03<br>Bl vs Tr: 3.1773e-07<br>Tr vs Ut: 2.1951e-04 | 1.46E-06 | 29.89 | 3 | Bl vs Na: 2.9255e-02<br>Bl vs Tr: 4.5964e-05<br>Na vs Ut: 2.6668e-02<br>Tr vs Ut: 1.2260e-05 |
| | 40 | 1.93E-08 | 38.78 | 3 | Bl vs Na: 5.0004e-04<br>Bl vs Tr: 1.1133e-08<br>Bl vs Ut: 2.2282e-02<br>Tr vs Ut: 2.8742e-04 | 2.25E-07 | 33.74 | 3 | Bl vs Na: 1.4663e-03<br>Bl vs Tr: 2.0916e-07<br>Tr vs Ut: 4.0450e-04 |
| | 50 | 1.78E-05 | 24.70 | 3 | Bl vs Na: 1.1158e-02<br>Bl vs Tr: 8.8748e-06<br>Bl vs Ut: 4.9909e-02<br>Tr vs Ut: 7.8737e-03 | 6.16E-05 | 22.12 | 3 | Bl vs Na: 2.5730e-02<br>Bl vs Tr: 2.9233e-05<br>Tr vs Ut: 1.4129e-02 |
| vNC | 20 | | | | - | 0.03128 | 8.86 | 3 | Bl vs Ut: 0.0249 |
| | 30 | 0.0108 | 11.17 | 3 | Tr vs Ut: 0.0455 | 0.03872 | 8.38 | 3 | - |
| | 40 | 2.96E-05 | 23.65 | 3 | Bl vs Tr: 7.7687e-06<br>Bl vs Ut: 1.4937e-02<br>Na vs Tr: 3.9034e-02<br>Tr vs Ut: 3.9634e-02 | 0.00051 | 17.69 | 3 | Bl vs Tr: 0.0001 |
| | 50 | 0.001999 | 14.80 | 3 | Bl vs Tr: 0.0022<br>Na vs Tr: 0.0297 | 0.007796 | 11.88 | 3 | Bl vs Tr: 0.0280<br>Na vs Tr: 0.0239<br>Tr vs Ut: 0.0403 |
| | 60 | 0.004441 | 13.09 | 3 | Na vs Tr: 0.0077<br>Tr vs Ut: 0.0148 | 0.002419 | 14.39 | 3 | Na vs Tr: 0.0059<br>Tr vs Ut: 0.0052 |
| mNCL | 20 | 2.81E-08 | 38.01 | 3 | Bl vs Na: 4.1674e-02<br>Bl vs Tr: 7.8858e-09<br>Bl vs Ut: 3.0420e-02<br>Na vs Tr: 1.2458e-03<br>Tr vs Ut: 1.2365e-04 | 0.000534 | 17.59 | 3 | Bl vs Tr: 0.0151<br>Na vs Tr: 0.0154<br>Tr vs Ut: 0.0002 |
| | 30 | < 2.2e-16 | 79.83 | 3 | Bl vs Na: 2.6309e-06<br>Bl vs Tr: 3.8931e-18<br>Bl vs Ut: 2.1389e-05<br>Na vs Tr: 1.8467e-04<br>Tr vs Ut: 7.9668e-07 | 4.03E-14 | 65.45 | 3 | Bl vs Na: 1.0602e-04<br>Bl vs Tr: 4.9156e-15<br>Bl vs Ut: 3.0858e-04<br>Na vs Tr: 2.3652e-04<br>Tr vs Ut: 3.9870e-06 |
| | 40 | < 2.2e-16 | 96.31 | 3 | Bl vs Na: 2.2412e-07<br>Bl vs Tr: 1.1804e-21<br>Bl vs Ut: 2.7063e-06<br>Na vs Tr: 1.8470e-05<br>Tr vs Ut: 1.4012e-08 | < 2.2e-16 | 82.25 | 3 | Bl vs Na: 4.2816e-06<br>Bl vs Tr: 1.5906e-18<br>Bl vs Ut: 4.3885e-05<br>Na vs Tr: 4.8581e-05<br>Tr vs Ut: 6.6858e-08 |
| | 50 | < 2.2e-16 | 80.26 | 3 | Bl vs Na: 5.5715e-06<br>Bl vs Tr: 2.4181e-17<br>Bl vs Ut: 2.4742e-05<br>Na vs Tr: 2.8534e-05<br>Tr vs Ut: 1.6873e-08 | < 2.2e-16 | 78.56 | 3 | Bl vs Na: 6.0695e-06<br>Bl vs Tr: 1.3745e-16<br>Bl vs Ut: 1.3537e-04<br>Na vs Tr: 7.5297e-05<br>Tr vs Ut: 4.4473e-09 |
| | 60 | 5.20E-09 | 41.47 | 3 | Bl vs Na: 1.6285e-03<br>Bl vs Tr: 6.4245e-08<br>Bl vs Ut: 3.1659e-03<br>Na vs Tr: 2.9580e-03<br>Tr vs Ut: 2.7759e-05 | 4.44E-09 | 41.79 | 3 | Bl vs Na: 8.1200e-04<br>Bl vs Tr: 3.1963e-08<br>Bl vs Ut: 2.4197e-03<br>Na vs Tr: 2.8485e-03<br>Tr vs Ut: 4.9739e-05 |
| | 70 | 9.16E-06 | 26.08 | 3 | Na vs Tr: 0.0001<br>Tr vs Ut: 0.0001 | 6.84E-06 | 26.69 | 3 | Bl vs Tr: 3.2846e-02<br>Na vs Tr: 5.1357e-04 |

| | | | | | | | | | |
|---|---|---|---|---|---|---|---|---|---|
| | | | | | | | | | Tr vs Ut: 5.3509e-05 |
| lNCL | 20 | 0.002 | 14.79 | 3 | Bl vs Na: 0.0462<br>Bl vs Tr: 0.0149<br>Tr vs Ut: 0.0194 | 0.01482 | 10.49 | 3 | Tr vs Ut: 0.01731569 |
| | 30 | 6.74E-08 | 36.22 | 3 | Bl vs Na: 8.4806e-05<br>Bl vs Tr: 2.0187e-07<br>Bl vs Ut: 3.7966e-02<br>Tr vs Ut: 8.5413e-04 | 2.47E-06 | 28.79 | 3 | Bl vs Na: 4.3451e-04<br>Bl vs Tr: 7.5684e-06<br>Tr vs Ut: 4.1220e-03 |
| | 40 | 5.88E-12 | 55.32 | 3 | Bl vs Na: 6.2025e-07<br>Bl vs Tr: 3.5984e-12<br>Bl vs Ut: 8.3372e-05<br>Tr vs Ut: 6.2643e-04 | 1.09E-11 | 54.06 | 3 | Bl vs Na: 4.1766e-06<br>Bl vs Tr: 7.9563e-12<br>Bl vs Ut: 1.5311e-03<br>Na vs Tr: 4.1158e-02<br>Tr vs Ut: 4.7434e-05 |
| | 50 | 7.96E-10 | 45.31 | 3 | Bl vs Na: 8.2402e-06<br>Bl vs Tr: 1.9344e-09<br>Bl vs Ut: 6.2895e-03<br>Tr vs Ut: 9.3860e-05 | 5.62E-11 | 50.72 | 3 | Bl vs Na: 8.5389e-07<br>Bl vs Tr: 1.3496e-10<br>Bl vs Ut: 1.4748e-03<br>Na vs Ut: 2.9203e-02<br>Tr vs Ut: 8.5504e-05 |
| | 60 | 0.0009078 | 16.47 | 3 | Bl vs Na: 0.0279<br>Bl vs Tr: 0.0007<br>Tr vs Ut: 0.0473 | 2.85E-05 | 23.73 | 3 | Bl vs Na: 2.0767e-03<br>Bl vs Tr: 1.8713e-05<br>Bl vs Ut: 1.7150e-02<br>Tr vs Ut: 1.4338e-02 |
| | 70 | 0.0073 | 12.01 | 3 | Bl vs Tr: 0.0047 | 0.007757 | 11.89 | 3 | Bl vs Na: 0.0340<br>Bl vs Tr: 0.0037<br>Bl vs Ut: 0.0291 |

KW – Kruskal-Wallis test
