## Supplementary material for "Changes in the Dopaminergic circuitry and Adult Neurogenesis linked to Reinforcement Learning in Corvids": Table S4

**Table S4.** Statistical details of structural analysis of Arc and DARPP-32 (active) colabeled neurons.

| Area | Neuron Morphometry parameters | Kruskal-Wallis P value | χ2 value | df | Post hoc P value (Dunn's test) |
|---|---|---|---|---|---|
| mNCL | Nodes | 2.52E-02 | 7.36 | 2 | Na vs Tr: 0.0271 |
| lNCL | Neurite length | 6.52E-05 | 19.275 | 2 | Na vs Tr: 2.4718e-02<br>Tr vs Ut: 3.4199e-05 |
| | Nodes | 1.15E-02 | 8.9323 | 2 | Na vs Tr: 0.0231<br>Tr vs Ut: 0.0191 |
| | Cell body area | 2.49E-02 | 7.3861 | 2 | Na vs Ut: 0.0464 |
| | Neurite field area | 5.88E-06 | 24.089 | 2 | Na vs Tr: 3.2e-02<br>Tr vs Ut: 3.3784e-06<br>Na vs Ut: 2.3523e-02 |
