## Supplementary material for "Changes in the Dopaminergic circuitry and Adult Neurogenesis linked to Reinforcement Learning in Corvids": Table S5

**Table S5.** Statistical details of Sholl analysis of Arc and DARPP-32 (active) colabeled neurons.

| Area | Radius (μm) | Intersections (P values) | χ2 value | df | Intersections (Dunn's P value) | Neurite length (P values) | χ2 value | df | Neurite length (Dunn's/Tukey's test P value) |
|---|---|---|---|---|---|---|---|---|---|
| lNCL | 30 | 0.0003 | 16.12 | 2 | Tr vs Ut: 0.0001 | 0.0091 | 5.05 | 2, 67 | Tr vs Ut: 0.0066 |
| | 40 | 0.001 | 13.68 | 2 | Tr vs Ut: 0.0008 | 0.000711 | 14.50 | 2 | Tr vs Ut: 0.0004 |
| | 50 | 0.0001 | 17.31 | 2 | Tr vs Ut: 9.8013e-05 | 0.0002046 | 16.99 | 2 | Tr vs Ut: 0.0001 |
| | 60 | 0.0186 | 7.96 | 2 | Na vs Tr: 0.0466<br>Tr vs Ut: 0.0383 | 0.003978 | 11.05 | 2 | Na vs Tr: 0.0220<br>Tr vs Ut: 0.0074 |
