## Supplementary material for "Changes in the Dopaminergic circuitry and Adult Neurogenesis linked to Reinforcement Learning in Corvids": Table S6

**Table S6.** Statistical details of structural analysis of inactive DARPP-32 neurons in NCL.

| Area | Neuron Morphometry parameters | Kruskal-Wallis/ ANOVA P value | χ2 / F value | df | Post hoc P value (Dunn’s) |
|---|---|---|---|---|---|
| mNCL | Neurite length | 6.14E-09 | 37.816 | 2 | Na vs Ut: 5.126717e-07<br>Tr vs Ut: 8.460955e-07 |
| | Endings | 2.68E-11 | 35.78 | 2, 67 | Na vs Ut: <1e-05<br>Tr vs Ut: <1e-05 |
| | Nodes | 2.74E-08 | 34.827 | 2 | Na vs Ut: 9.844824e-07<br>Tr vs Ut: 1.894894e-06 |
| | Cell body area | 4.83E-05 | 11.57 | 2 | Na vs Ut: <1e-04<br>Tr vs Na: 0.0075 |
| | Neurite field area | 1.67E-08 | 35.812 | 2 | Na vs Ut: 9.844824e-07<br>Tr vs Ut: 1.894894e-06 |
| lNCL | Neurite length | 1.85E-09 | 40.213 | 2 | Na vs Ut: 4.548690e-08<br>Tr vs Ut: 2.066172e-06 |
| | Endings | 2.42E-08 | 35.075 | 2 | Na vs Ut: 4.414069e-07<br>Tr vs Ut: 8.300441e-06 |
| | Nodes | 1.17E-08 | 36.526 | 2 | Na vs Ut: 2.144890e-07<br>Tr vs Ut: 9.387586e-07 |
| | Cell body area | 6.69E-04 | 54.004 | 2, 67 | Na vs Ut: 0.00924<br>Tr vs Ut: 0.00148 |
| | Neurite field area | 3.53E-09 | 38.922 | 2 | Na vs Ut: 2.144890e-07<br>Tr vs Ut: 9.387586e-07 |
