## Supplementary material for "Changes in the Dopaminergic circuitry and Adult Neurogenesis linked to Reinforcement Learning in Corvids": Table S7

**Table S7.** Statistical details of Sholl analysis of inactive DARPP-32 neurons in NCL.

| Area | Radius (µm) | Intersections ( P-Value) | χ2 / F value | df | Intersections (Dunn's / Tukey's P value) | Neurite length ( P-Value) | χ2 / F value | df | Neurite length (Dunn's / Tukey's P value) |
|---|---|---|---|---|---|---|---|---|---|
| mNCL | 20 | 3.25E-07 | 18.83 | 2, 67 | Na vs Ut: <1e-05<br>Tr vs Ut: 4.94e-05 | 6.91E-06 | 14.26 | 2, 67 | Na vs Ut: <1e-04<br>Tr vs Ut: 0.00281 |
| | 30 | 7.47E-09 | 25.06 | 2, 67 | Na vs Ut: <1e-06<br>Tr vs Ut: <1e-06 | 3.17E-09 | 26.58 | 2, 67 | Na vs Ut: <1e-05<br>Tr vs Ut: <1e-05 |
| | 40 | 3.22E-08 | 22.76 | 2, 65 | Na vs Ut: <1e-04<br>Tr vs Ut: <1e-04 | 2.73E-08 | 23.04 | 2, 65 | Na vs Ut: <1e-05<br>Tr vs Ut: <1e-05 |
| | 50 | 2.06E-08 | 23.87 | 2, 62 | Na vs Ut: <1e-05<br>Tr vs Ut: <1e-05 | 3.75E-08 | 22.81 | 2, 62 | Na vs Ut: <1e-04<br>Tr vs Ut: <1e-04 |
| | 60 | 0.000144 | 10.37 | 2, 57 | Na vs Ut: 0.00147<br>Tr vs Ut: 0.00033 | 9.90E-06 | 14.2 | 2, 57 | Na vs Ut: 0.00016<br>Tr vs Ut: <1e-04 |
| | 70 | 0.0374 | 6.5721 | 2 | - | 0.0301 | 3.756 | 2, 51 | Na vs Ut: 0.0429 |
| lNCL | 20 | 0.000152 | 10.06 | 2, 67 | Na vs Ut: 0.00032<br>Tr vs Ut: 0.00439 | 0.0169 | 4.339 | 2, 67 | Na vs Ut: 0.0147 |
| | 30 | 2.07E-11 | 36.32 | 2, 67 | Na vs Ut: <1e-05<br>Tr vs Ut: <1e-05 | 2.67E-07 | 30.27 | 2 | Na vs Ut: 2.9256e-06<br>Tr vs Ut: 4.1204e-05 |
| | 40 | 5.67E-09 | 37.975 | 2 | Na vs Ut: 3.143e-07<br>Tr vs Ut: 1.0734e-06 | 1.37E-11 | 37.41 | 2, 66 | Na vs Ut: <1e-05<br>Tr vs Ut: <1e-05 |
| | 50 | 2.44E-07 | 30.453 | 2 | Na vs Ut: 2.534947e-06<br>Tr vs Ut: 1.666073e-05 | 1.31E-07 | 31.69 | 2 | Na vs Ut: 1.8189e-06<br>Tr vs Ut: 8.8622e-06 |
| | 60 | 6.19E-05 | 19.379 | 2 | Na vs Ut: 0.0001<br>Tr vs Ut: 0.0009 | 3.93E-05 | 20.289 | 2 | Na vs Ut: 7.9315e-05<br>Tr vs Ut: 5.5381e-04 |
| | 70 | 0.01246 | 8.7698 | 2 | Na vs Ut: 0.0099 | 0.0137 | 8.5801 | 2 | Na vs Ut: 0.0102 |
