## Supplementary material for "Changes in the Dopaminergic circuitry and Adult Neurogenesis linked to Reinforcement Learning in Corvids": Table S8

**Table S8.** Statistical information of comparisons between active and inactive DARPP-32 neurons structural features in NCL.

| Area | Experimental group | Parameter | Test | n | df | t/W | P |
|---|---|---|---|---|---|---|---|
| lNCL | Undertrained | Neurite field area | Wilcoxon rank sum test | 30, 30 | | 302 | 0.02847 |
| | Trained | Neurite length | Wilcoxon rank sum test | 20, 20 | | 73 | 0.00037 |
| | Trained | Nodes | Wilcoxon rank sum test | 20, 20 | | 113 | 0.01864 |
| | Trained | Endings | Wilcoxon rank sum test | 20, 20 | | 103 | 0.0088 |
| | Trained | Neurite field area | Wilcoxon rank sum test | 20, 20 | | 79 | 0.00075 |
| | No-Association | Neurite length | Wilcoxon rank sum test | 20, 20 | | 33 | 7.57E-07 |
| | No-Association | Nodes | Wilcoxon rank sum test | 20, 20 | | 60.5 | 0.00016 |
| | No-Association | Endings | Wilcoxon rank sum test | 20, 20 | | 52 | 6.35E-05 |
| | No-Association | Neurite field area | Wilcoxon rank sum test | 20, 20 | | 33 | 7.57E-07 |
| mNCL | Undertrained | Nodes | Wilcoxon rank sum test | 30, 30 | | 681.5 | 0.0006 |
| | | Endings | Wilcoxon rank sum test | 30, 30 | | 679.5 | 0.00068 |
| | | Cell body area | Welch's t test | 30, 30 | 57.007 | 2.2074 | 0.03133 |
| | Trained | Neurite length | Wilcoxon rank sum test | 20, 20 | | 95 | 0.00389 |
| | Trained | Neurite field area | Wilcoxon rank sum test | 20, 20 | | 104 | 0.00871 |
| | No-Association | Neurite length | Welch's t test | 20, 20 | 36.602 | -4.5857 | 5.13E-05 |
| | No-Association | Nodes | Welch's t test | 20, 20 | 37.714 | -3.1189 | 0.00347 |
| | No-Association | Endings | Welch's t test | 20, 20 | 37.832 | -4.0088 | 2.77E-04 |
| | No-Association | Neurite field area | Welch's t test | 20, 20 | 36.844 | -3.445 | 1.44E-03 |
