## Supplementary material for "Changes in the Dopaminergic circuitry and Adult Neurogenesis linked to Reinforcement Learning in Corvids": Table S9

**Table S9.** Statistical details of DCX positive neuron numbers across various regions.

| Area | Neuron Morphometry parameters | ANOVA P value | χ2 value | df | Post hoc P value (Tukey's) |
|---|---|---|---|---|---|
| Area X | Spherical | 0.00272 | 7.751 | 3, 14 | Tr vs Bl: 0.00164 |
| | | | | | Tr vs Ut: 0.02079 |
| MSt | Spherical | 0.00147 | 8.942 | 3, 14 | Tr vs Bl: 0.00127 |
| | | | | | Tr vs Ut: 0.00611 |
| dNC | Spherical | 0.0196 | 4.577 | 3, 14 | Tr vs Bl: 0.0194 |
| iNC | Spherical | 0.0161 | 4.857 | 3, 14 | Tr vs Bl: 0.00951 |
| mNCL | Spherical | 0.0267 | 4.153 | 3, 14 | Tr vs Bl: 0.0248 |
| lNCL | Spherical | 0.0387 | 3.667 | 3, 14 | Tr vs Bl: 0.0304 |
| vNC | Spherical | 0.0175 | 4.738 | 3, 14 | Tr vs Bl: 0.0104 |
