## Supplementary material for "Changes in the Dopaminergic circuitry and Adult Neurogenesis linked to Reinforcement Learning in Corvids": Table S10

**Table S10.** Statistical details of structural analysis of DCX neurons in Area X and MSt.

| Area | Neuron Morphometry parameters | Kruskal P value | χ2 value | df | Post hoc P value (Dunn's) |
|---|---|---|---|---|---|
| Area X | Neurite length | 1.68E-15 | 71.886 | 3 | Bl vs Na: 2.7614e-13<br>Bl vs Tr: 1.0340e-06<br>Na vs Tr: 3.3577e-02<br>Na vs Ut: 1.0028e-09<br>Tr vs Ut: 5.4905e-04 |
| | Endings | 3.19E-07 | 33.022 | 3 | Bl vs Na: 7.0659e-07<br>Bl vs Tr: 2.9107e-02<br>Na vs Tr: 2.6922e-02<br>Na vs Ut: 1.9751e-05 |
| | Node | 1.83E-06 | 29.421 | 3 | Bl vs Na: 8.2048e-06<br>Na vs Tr: 1.9709e-02<br>Na vs Ut: 1.5103e-05 |
| | Cell body area | 7.19E-07 | 31.346 | 3 | Bl vs Na: 1.1221e-04<br>Bl vs Tr: 1.6871e-06<br>Bl vs Ut: 7.0123e-05 |
| | Convex hull area | <2.2e-16 | 96.007 | 3 | Bl vs Na: 8.9660e-18<br>Bl vs Tr: 3.0441e-10<br>Bl vs Ut: 9.7218e-03<br>Na vs Tr: 2.2490e-02<br>Na vs Ut: 4.7805e-11<br>Tr vs Ut: 4.8279e-05 |
| MSt | Neurite length | <2.2e-16 | 97.362 | 3 | Bl vs Na: 2.5606e-18<br>Bl vs Tr: 7.7442e-09<br>Bl vs Ut: 1.2189e-02<br>Na vs Tr: 6.8511e-03<br>Na vs Ut: 1.7258e-12<br>Tr vs Ut: 1.4138e-04 |
| | Endings | 3.16E-11 | 51.892 | 3 | Bl vs Na: 6.4528e-10<br>Bl vs Tr: 1.4710e-02<br>Na vs Tr: 1.0794e-03<br>Na vs Ut: 4.2590e-09 |
| | Node | 3.73E-10 | 46.856 | 3 | Bl vs Na: 2.2765e-09<br>Na vs Tr: 9.5374e-05<br>Na vs Ut: 4.9110e-08 |
| | Cell body area | 5.21E-06 | 27.252 | 3 | Bl vs Na: 1.4980e-03<br>Bl vs Tr: 3.4184e-06<br>Bl vs U: 7.3460e-04 |
| | Convex hull area | <2.2e-16 | 104.73 | 3 | Bl vs Na: 2.3367e-20<br>Bl vs Tr: 3.0679e-10<br>Bl vs Ut: 2.7254e-03<br>Na vs Tr: 3.4063e-03<br>Na vs Ut: 4.8908e-12<br>Tr vs Ut: 2.5958e-04 |
