## Supplementary material for "Changes in the Dopaminergic circuitry and Adult Neurogenesis linked to Reinforcement Learning in Corvids": Table S11

**Table S11**. Statistical details of Sholl analysis of inactive DCX neurons in Area X and MSt.

| **Area** | **Radius (μm)** | **Intersections (P value)** | **χ2** | **df** | **Intersections (Dunn's P value)** | **Neurite length (P value)** | **χ2** | **df** | **Neurite length (Dunn's P value)** |
|---|---|---|---|---|---|---|---|---|---|
| Area X | 28 | 1.05E-08 | 40.03 | 3 | Bl vs Na: 8.8285e-08<br>Bl vs Tr: 1.2090e-04<br>Na vs Ut: 4.5461e-05<br>Tr vs Ut: 1.5231e-02 | 1.62E-06 | 29.67 | 3 | Bl vs Na: 4.7901e-06<br>Bl vs Tr: 1.7566e-03<br>Na vs Ut: 6.1731e-04 |
| | 38 | <2.2e-16 | 86.07 | 3 | Bl vs Na: 9.3555e-16<br>Bl vs Tr: 2.0132e-09<br>Bl vs Ut: 1.8099e-02<br>Na vs Tr: 3.9360e-02<br>Na vs Ut: 5.0444e-10<br>Tr vs Ut: 7.3697e-05 | 2.32E-15 | 71.24 | 3 | Bl vs Na: 7.5432e-13<br>Bl vs Tr: 2.5125e-08<br>Bl vs Ut: 3.4016e-02<br>Na vs Ut: 3.2989e-08<br>Tr vs Ut: 1.6367e-04 |
| | 48 | 5.27E-14 | 64.90 | 3 | Bl vs Na: 6.4355e-14<br>Bl vs Tr: 2.5824e-06<br>Bl vs Ut: 5.7402e-03<br>Na vs Tr: 3.3033e-03<br>Na vs Ut: 7.0574e-07<br>Tr vs Ut: 3.4252e-02 | 2.25E-14 | 66.63 | 3 | BL vs Na: 1.0655e-13<br>Bl vs Tr: 4.1978e-07<br>Bl vs Ut: 6.4435e-03<br>Na vs Tr: 1.1140e-02<br>Na vs Ut: 3.2540e-07<br>Tr vs Ut: 1.3019e-02 |
| | 58 | 1.54E-07 | 34.52 | 3 | Bl vs Na: 3.2219e-07<br>Bl vs Tr: 2.1425e-03<br>Bl vs Ut: 3.7791e-02<br>Na vs Tr: 4.3333e-02<br>Na vs Ut: 2.7473e-04 | 7.13E-11 | 50.23 | 3 | Bl vs Na: 1.7291e-10<br>Bl vs Tr: 9.2329e-06<br>Bl vs Ut: 4.0353e-03<br>Na vs Tr: 4.4294e-02<br>Na vs Ut: 2.8619e-05<br>Tr vs Ut: 3.8143e-02 |
| | 68 | 0.01377 | 10.65 | 3 | Bl vs Na: 0.00898 | 0.005703 | 12.56 | 3 | Bl vs Na: 0.00515 |
| MSt | 18 | 2.52E-05 | 23.98 | 3 | Bl vs Na: 0.0070<br>Bl vs Tr: 0.0278<br>Na vs Ut: 0.0002<br>Tr vs Ut: 0.0028 | 0.000686 | 17.06 | 3 | Na vs Ut: 0.0025<br>Tr vs Ut: 0.0140 |
| | 28 | <2.2e-16 | 76.56 | 3 | Bl vs Na: 6.7050e-13<br>Bl vs Tr: 2.4917e-10<br>Bl vs Ut: 1.2603e-02<br>Na vs Ut: 2.6161e-07<br>Tr vs Ut: 1.2414e-05 | 2.24E-12 | 57.28 | 3 | Bl vs Na: 2.4572e-09<br>Bl vs Tr: 4.9921e-07<br>Na vs Ut: 7.6749e-07<br>Tr vs Ut: 6.2201e-05 |
| | 38 | <2.2e-16 | 97.32 | 3 | Bl vs Na: 5.1717e-16<br>Bl vs Tr: 5.7017e-13<br>Bl vs Ut: 5.0555e-03<br>Na vs Ut: 3.7436e-09<br>Tr vs Ut: 3.9490e-07 | < 2.2e-16 | 103.53 | 3 | Bl vs Na: 2.2520e-17<br>Bl vs Tr: 2.9542e-13<br>Bl vs Ut: 3.7772e-03<br>Na vs Ut: 5.0266e-10<br>Tr vs Ut: 3.6621e-07 |
| | 48 | < 2.2e-16 | 83.46 | 3 | Bl vs Na: 3.9975e-14<br>Bl vs Tr: 4.8811e-09<br>Bl vs Ut: 4.1834e-02<br>Na vs Ut: 2.5992e-10<br>Tr vs Ut: 1.1173e-05 | < 2.2e-16 | 85.99 | 3 | Bl vs Na: 4.7081e-14<br>Bl vs Tr: 5.4184e-10<br>Bl vs Ut: 3.7562e-02<br>Na vs Ut: 4.1003e-10<br>Tr vs Ut: 1.9306e-06 |

| | | | | | | | | |
|---|---|---|---|---|---|---|---|---|
| 58 | 6.96E-08 | 36.15 | 3 | Bl vs Na: 5.9417e-06<br>Bl vs Tr: 4.7789e-03<br>Na vs Ut: 5.0157e-06<br>Tr vs Ut: 1.9323e-02 | 1.29E-10 | 49.03 | 3 | Bl vs Na: 2.3068e-07<br>Bl vs Tr: 1.3014e-04<br>Na vs Ut: 1.5210e-07<br>Tr vs Ut: 4.1277e-04 |
| 68 | 0.00021 | 19.50 | 3 | Bl vs Na: 0.0070<br>Bl vs Tr: 0.0079<br>Na vs Ut: 0.0081<br>Tr vs Ut: 0.0130 | 2.67E-05 | 23.86 | 3 | Bl vs Na: 0.0004<br>Bl vs Tr: 0.0004<br>Na vs Ut: 0.0131<br>Tr vs Ut: 0.0148 |
