## Supplementary material for "Changes in the Dopaminergic circuitry and Adult Neurogenesis linked to Reinforcement Learning in Corvids": Table S12

**Table S12.** Statistical details of structural analysis of active and inactive DCX neurons in MSt.

| Cell type | Area | Neuron Morphometry parameters | Kruskal-Wallis P value | χ2 value | df | Post hoc P value (Dunn's) |
|---|---|---|---|---|---|---|
| Active | MSt | Cell body area | 0.0123 | 8.7885 | 2 | Tr vs Ut: 0.0092 |
| Inactive | MSt | Neurite length | 2.93E-08 | 34.691 | 2 | Na vs Ut: 2.954254e-06<br>Tr vs Ut: 1.429155e-06 |
| | | Endings | 2.67E-08 | 34.88 | 2 | Na vs Ut: 7.2774e-06<br>Tr vs Ut: 5.6143e-07 |
| | | Nodes | 2.35E-08 | 35.13 | 2 | Na vs Ut: 9.8818e-06<br>Tr vs Ut: 3.6874e-07 |
| | | Neurite field area | 3.84E-07 | 29.545 | 2 | Na vs Ut: 1.1762e-05<br>Tr vs Ut: 1.5009e-05 |
