## Supplementary material for "Changes in the Dopaminergic circuitry and Adult Neurogenesis linked to Reinforcement Learning in Corvids": Table S13

**Table S13.** Statistical details of Sholl analysis of active and inactive DCX neurons in MSt.

| Cell type | Area | Radius (µm) | df | Intersections (Kruskal-Wallis / ANOVA P-Value) | Intersections (Dunn's / Tukey's P value) | Neurite length (Kruskal-Wallis / ANOVA P-Value) | χ2/F value | df | Neurite length (Dunn's / Tukey's P value) |
|---|---|---|---|---|---|---|---|---|---|
| Active | MSt | 18 | | | - | 0.0156 | 4.43 | 2, 67 | Tr vs Ut: 0.0121 |
| Inactive | MSt | 18 | 2 | 1.25E-04 | Na vs Ut: 0.0006<br>Tr vs Ut: 0.0014 | 4.21E-04 | 15.54 | 2 | Na vs Ut: 0.0015<br>Tr vs Ut: 0.0038 |
| | | 28 | 2, 67 | 5.31E-08 | Na vs Ut: <1e-04<br>Tr vs Ut: <1e-04 | 4.43E-07 | 18.34 | 2, 67 | Na vs Ut: <1e-04<br>Tr vs Ut: <1e-04 |
| | | 38 | 2, 66 | 3.01E-07 | Na vs Ut: <1e-04<br>Tr vs Ut: <1e-04 | 2.31E-07 | 19.43 | 2, 66 | Na vs Ut: <1e-05<br>Tr vs Ut: <1e-05 |
| | | 48 | 2 | 1.11E-06 | Na vs Ut: 0.00002<br>Tr vs Ut: 0.00002 | 2.34E-07 | 19.49 | 2, 65 | Na vs Ut: <1e-05<br>Tr vs Ut: <1e-05 |
| | | 58 | 2 | 3.59E-05 | Na vs Ut: 1.3675e-03<br>Tr vs Ut: 7.7969e-05 | 1.08E-05 | 13.88 | 2, 61 | Na vs Ut: <1e-04<br>Tr vs Ut: <1e-04 |
| | | 68 | 2 | 0.01022 | Na vs Ut: 0.0233<br>Tr vs Ut: 0.0176 | 0.00433 | 10.88 | 2 | Na vs Ut: 0.0264<br>Tr vs Ut: 0.0047 |
