## Supplementary material for "Changes in the Dopaminergic circuitry and Adult Neurogenesis linked to Reinforcement Learning in Corvids": Table S14

**Table S14.** Statistical information of comparisons between active and inactive DCX neurons structural features in MSt.

| Area | Experimental group | Parameter | Test | n | df | t/W | P |
|---|---|---|---|---|---|---|---|
| MSt | Undertrained | Neurite length | Wilcoxon rank sum test | 30, 30 | | 270 | 0.0073 |
| | Undertrained | Endings | Wilcoxon rank sum test | 30, 30 | | 208.5 | 0.0003 |
| | Undertrained | Nodes | Wilcoxon rank sum test | 30, 30 | | 253.5 | 0.0035 |
| | Undertrained | Neurite field area | Wilcoxon rank sum test | 30, 30 | | 270 | 0.0073 |
| | Undertrained | Soma Area | Welch's t-test | 30, 30 | 56.485 | -3.3243 | 0.0015 |
| | No-Association | Soma Area | Welch's t-test | 20, 20 | 35.069 | -2.4415 | 0.01981 |
