## Supplementary material for "Changes in the Dopaminergic circuitry and Adult Neurogenesis linked to Reinforcement Learning in Corvids": Table S15

**Table S15**. Statistical details of structural analysis of DCX neurons in NC subdivisions.

| Area | Neuron Morphometry parameters | Kruskal P value | χ2 value | df | Post hoc P value (Dunn's) |
|---|---|---|---|---|---|
| dNC | Neurite length | < 2.2e-16 | 87.582 | 3 | Bl vs Na: 2.818817e-12<br>Bl vs Tr: 1.343757e-17<br>Bl vs Ut: 3.639237e-07<br>Na vs Ut: 2.135021e-02<br>Tr vs Ut: 6.848278e-05 |
| | Nodes | 1.54E-11 | 53.358 | 3 | Bl vs Na: 3.565885e-08<br>Bl vs Tr: 8.021323e-10<br>Bl vs Ut: 4.405485e-03<br>Na vs Ut: 3.127850e-03<br>Tr vs Ut: 2.820970e-04 |
| | Neurite field area | 4.46E-15 | 69.912 | 3 | Bl vs Na: 2.485072e-10<br>Bl vs Tr: 1.792783e-13<br>Bl vs Ut: 1.236318e-04<br>Na vs Ut: 3.890183e-03<br>Tr vs Ut: 9.620369e-05 |
| | Soma Area | 0.006497 | 12.275 | 3 | Bl vs Na: 0.00513 |
| iNC | Neurite length | 2.48E-13 | 61.752 | 3 | Bl vs Na: 5.156065e-13<br>Bl vs Tr: 2.415872e-08<br>Bl vs Ut: 1.381076e-05<br>Na vs Ut: 1.241345e-03 |
| | Nodes | 0.0003052 | 18.769 | 3 | Bl vs Na: 0.00073<br>Bl vs Tr: 0.00599<br>Na vs Ut: 0.03407 |
| | Neurite field area | 4.01E-07 | 32.551 | 3 | Bl vs Na: 2.736157e-07<br>Bl vs Tr: 7.344751e-03<br>Na vs Ut: 2.343574e-04 |
| | Soma Area | 0.008818 | 11.617 | 3 | Bl vs Na: 0.03804<br>Bl vs Tr: 0.01658 |
| mNCL | Neurite length | < 2.2e-16 | 85.203 | | Bl vs Na: 1.694703e-12<br>Bl vs Tr: 9.235544e-17<br>Bl vs Ut: 1.454247e-09<br>Tr vs Ut: 6.447023e-03 |
| | Nodes | 3.69E-11 | 51.573 | | Bl vs Na: 3.886423e-06<br>Bl vs Tr: 3.087404e-11<br>Bl vs Ut: 1.126038e-03<br>Tr vs Ut: 2.497804e-04 |
| | Neurite field area | 2.12E-14 | 66.75 | | Bl vs Na: 3.198573e-11<br>Bl vs Tr: 2.285996e-12<br>Bl vs Ut: 5.475690e-07<br>Na vs Ut: 4.802876e-02<br>Tr vs Ut: 2.169973e-02 |
| lNCL | Neurite length | < 2.2e-16 | 79.901 | | Bl vs Na: 5.657415e-14<br>Bl vs Tr: 5.144917e-14 |

| | | | | |
|---|---|---|---|---|
| | | | | Bl vs Ut: 5.819495e-08 |
| | | | | Na vs Ut: 1.443792e-02 |
| | | | | Tr vs Ut: 9.996455e-03 |
| | Nodes | 1.28E-08 | 39.632 | Bl vs Na: 5.092833e-08 |
| | | | | Bl vs Tr: 3.234468e-06 |
| | | | | Bl vs Ut: 3.641940e-03 |
| | | | | Na vs Ut: 8.334678e-03 |
| | Neurite field area | 6.48E-07 | 31.561 | Bl vs Na: 2.509658e-05 |
| | | | | Bl vs Tr: 7.933987e-06 |
| | | | | Bl vs Ut: 3.975881e-02 |
| | | | | Na vs Ut: 2.268959e-02 |
| | | | | Tr vs Ut: 1.193686e-02 |
| vNC | Neurite length | 1.65E-10 | 48.523 | Bl vs Na: 1.883061e-09 |
| | | | | Bl vs Tr: 5.385661e-08 |
| | | | | Bl vs Ut: 3.106734e-04 |
| | | | | Na vs Ut: 2.204467e-02 |
| | Nodes | 0.0001229 | 20.676 | Bl vs Na: 0.00047 |
| | | | | Bl vs Tr: 0.00122 |
| | Neurite field area | 2.09E-07 | 33.894 | Bl vs Na: 4.981936e-07 |
| | | | | Bl vs Tr: 1.541250e-04 |
| | | | | Na vs Ut: 1.986350e-03 |
