## Supplementary material for "Changes in the Dopaminergic circuitry and Adult Neurogenesis linked to Reinforcement Learning in Corvids": Table S16

**Table S16**. Statistical details of Sholl analysis of DCX neurons in NC subdivisions.

| Area | Radius (μm) | Intersections (P value) | χ2 value | df | Intersections (Dunn's P value) | Neurite length (P value) | χ2 value | df | Neurite length (Dunn's P value) |
|---|---|---|---|---|---|---|---|---|---|
| dNC | 20 | 3.75E-10 | 46.84 | 3 | Bl vs Na: 4.585896e-06<br>Bl vs Tr: 2.896733e-10<br>Bl vs Ut: 3.080685e-04<br>Tr vs Ut: 3.458343e-03 | 1.06E-06 | 30.54 | 3 | Bl vs Na: 9.566860e-06<br>Bl vs Tr: 1.423749e-05<br>Bl vs Ut: 1.595202e-02<br>Tr vs Ut: 4.899398e-02 |
| | 30 | 4.65E-14 | 65.15 | 3 | Bl vs Na: 3.982865e-09<br>Bl vs Tr: 2.205398e-13<br>Bl vs Ut: 5.332721e-05<br>Na vs Ut: 3.440019e-02<br>Tr vs Ut: 2.231618e-04 | 9.38E-14 | 63.73 | 3 | Bl vs Na: 9.913265e-09<br>Bl vs Tr: 3.282826e-13<br>Bl vs Ut: 8.500791e-05<br>Na vs Ut: 4.012314e-02<br>Tr vs Ut: 1.827075e-04 |
| | 40 | 7.78E-08 | 35.92 | 3 | Bl vs Na: 3.883177e-05<br>Bl vs Tr: 2.675517e-07<br>Bl vs Ut: 2.591807e-02<br>Na vs Ut: 2.887066e-02<br>Tr vs Ut: 1.100249e-03 | 1.01E-08 | 40.10 | 3 | Bl vs Na: 1.524015e-06<br>Bl vs Tr: 6.732655e-08<br>Bl vs Ut: 5.165195e-03<br>Na vs Ut: 1.569242e-02<br>Tr vs Ut: 3.238231e-03 |
| | 50 | 1.16E-05 | 25.59 | 3 | Bl vs Na: 0.0001885800<br>Bl vs Tr: 0.0001131071<br>Bl vs Ut: 0.0486381887<br>Na vs Ut: 0.0209814017<br>Tr vs Ut: 0.0146355544 | 7.47E-05 | 21.72 | 3 | Bl vs Na: 0.0008980322<br>Bl vs Tr: 0.0011681818<br>Na vs Ut: 0.0169900890<br>Tr vs Ut: 0.0203214057 |
| | 60 | 4.79E-04 | 17.82 | 3 | Na vs Ut: 0.001744675<br>Tr vs Ut: 0.023201548 | 0.0004694 | 17.86 | 3 | Bl vs Na: 0.018244873<br>Bl vs Tr: 0.028744224<br>Na vs Ut: 0.006209491<br>Tr vs Ut: 0.027372011 |
| iNC | 20 | 4.50E-04 | 17.95 | 3 | Bl vs Na: 0.001055695<br>Bl vs Tr: 0.001887158<br>Bl vs Ut: 0.013421908 | | | | |
| | 30 | 1.82E-11 | 53.02 | 3 | Bl vs Na: 5.958740e-11<br>Bl vs Tr: 1.072118e-07<br>Bl vs Ut: 3.614554e-06<br>Na vs Ut: 3.271787e-02 | 3.00E-09 | 42.59 | 3 | Bl vs Na: 1.042134e-08<br>Bl vs Tr: 1.115683e-06<br>Bl vs Ut: 6.845753e-05 |
| | 40 | 5.58E-09 | 41.33 | 3 | Bl vs Na: 1.330541e-08<br>Bl vs Tr: 9.561257e-06<br>Bl vs Ut: 9.333206e-03<br>Na vs Ut: 7.456280e-04<br>Tr vs Ut: 4.030937e-02 | 1.95E-09 | 43.47 | 3 | Bl vs Na: 1.400143e-09<br>Bl vs Tr: 3.578969e-06<br>Bl vs Ut: 3.338069e-04<br>Na vs Ut: 5.724331e-03 |
| | 50 | 4.59E-07 | 32.27 | 3 | Bl vs Na: 9.550034e-07<br>Bl vs Tr: 1.718148e-03<br>Na vs Ut: 2.760243e-04 | 4.96E-06 | 27.35 | 3 | Bl vs Na: 3.649621e-06<br>Bl vs Tr: 2.351212e-03<br>Bl vs Ut: 4.116088e-02<br>Na vs Ut: 3.561309e-03 |
| | 60 | 1.16E-03 | 15.95 | 3 | Bl vs Na: 0.00769535<br>Bl vs Tr: 0.01636620<br>Na vs Ut: 0.02812947 | 0.001621 | 15.24 | 3 | Bl vs Na: 0.02387079<br>Bl vs Tr: 0.02537852<br>Na vs Ut: 0.02678765<br>Tr vs Ut: 0.04169965 |

| | | | | | | | | | |
|---|---|---|---|---|---|---|---|---|---|
| mNCL | 20 | 1.73E-08 | 39.00 | 3 | Bl vs Na: 1.983993e-04 | 2.33E-06 | 28.92 | 3 | Bl vs Na: 7.371346e-04 |
| | | | | | Bl vs Tr: 5.732879e-09 | | | | Bl vs Tr: 1.599728e-06 |
| | | | | | Bl vs Ut: 2.564593e-04 | | | | Bl vs Ut: 6.170838e-04 |
| | | | | | Tr vs Ut: 2.059328e-02 | | | | |
| | 30 | < 2.2e-16 | 79.42 | 3 | Bl vs Na: 5.736234e-10 | 2.77E-13 | 61.53 | 3 | Bl vs Na: 1.139491e-07 |
| | | | | | Bl vs Tr: 8.720984e-17 | | | | Bl vs Tr: 2.523560e-13 |
| | | | | | Bl vs Ut: 1.738298e-09 | | | | Bl vs Ut: 2.775307e-07 |
| | | | | | Tr vs Ut: 5.274188e-03 | | | | Tr vs Ut: 1.096734e-02 |
| | 40 | 1.03E-06 | 30.61 | 3 | Bl vs Na: 3.497607e-05 | 3.33E-09 | 42.38 | 3 | Bl vs Na: 2.830228e-06 |
| | | | | | Bl vs Tr: 5.245500e-07 | | | | Bl vs Tr: 1.042309e-09 |
| | | | | | Bl vs Ut: 2.332137e-04 | | | | Bl vs Ut: 3.374997e-05 |
| | | | | | | | | | Tr vs Ut: 1.469346e-02 |
| | 50 | 0.003764 | 13.45 | 3 | Bl vs Na: 0.019700334 | 0.007858 | 11.87 | 3 | Bl vs Tr: 0.01385610 |
| | | | | | Bl vs Tr: 0.009277887 | | | | |
| | 60 | 0.000101 | 21.07 | 3 | Bl vs Na: 0.025756400 | 0.002494 | 14.33 | 3 | Bl vs Na: 0.04418072 |
| | | | | | Na vs Ut: 0.000475648 | | | | Na vs Ut: 0.01182431 |
| | | | | | Tr vs Ut: 0.012925888 | | | | |
| lNCL | 20 | 8.74E-10 | 45.12 | 3 | Bl vs Na: 8.172467e-08 | 0.000165 | 20.05 | 3 | Bl vs Na: 0.0001484976 |
| | | | | | Bl vs Tr: 1.531016e-08 | | | | Bl vs Tr: 0.0033991177 |
| | | | | | Bl vs Ut: 2.170761e-04 | | | | Bl vs Ut: 0.0145200127 |
| | | | | | Tr vs Ut: 3.828554e-02 | | | | |
| | 30 | 1.63E-12 | 57.92 | 3 | Bl vs Na: 6.292596e-11 | 6.82E-14 | 64.38 | 3 | Bl vs Na: 1.654101e-11 |
| | | | | | Bl vs Tr: 9.879320e-10 | | | | Bl vs Tr: 2.989246e-11 |
| | | | | | Bl vs Ut: 2.183683e-05 | | | | Bl vs Ut: 1.742387e-05 |
| | | | | | Na vs Ut: 1.052563e-02 | | | | Na vs Ut: 6.190890e-03 |
| | | | | | Tr vs Ut: 2.949063e-02 | | | | Tr vs Ut: 6.144994e-03 |
| | 40 | 1.57E-10 | 48.63 | 3 | Bl vs Na: 2.018935e-09 | 1.63E-12 | 57.93 | 3 | Bl vs Na: 5.194983e-11 |
| | | | | | Bl vs Tr: 1.365312e-08 | | | | Bl vs Tr: 3.559509e-10 |
| | | | | | Bl vs Ut: 1.581720e-05 | | | | Bl vs Ut: 7.348038e-07 |
| | 50 | 8.11E-06 | 26.34 | 3 | Bl vs Na: 1.037431e-05 | 7.48E-06 | 26.50 | 3 | Bl vs Na: 1.293657e-05 |
| | | | | | Bl vs Tr: 7.554106e-05 | | | | Bl vs Tr: 2.622548e-05 |
| | | | | | Bl vs Ut: 7.881417e-03 | | | | Bl vs Ut: 2.904527e-03 |
| | 60 | 2.51E-05 | 23.99 | 3 | Bl vs Na: 5.555223e-05 | 0.000165 | 20.05 | 3 | Bl vs Na: 0.0001119339 |
| | | | | | Bl vs Tr: 3.376646e-04 | | | | Bl vs Tr: 0.0006402055 |
| | | | | | Bl vs Ut: 2.254855e-02 | | | | Bl vs Ut: 0.0104985715 |
| | | | | | Na vs Ut: 2.462277e-02 | | | | |
| | 70 | | | | | 0.04398 | 6.25 | 3 | Na vs Ut: 0.04461682 |
| vNC | 20 | 0.02899 | 9.02 | 3 | | | | | |
| | 30 | 1.74E-10 | 48.41 | 3 | Bl vs Na: 2.563286e-09 | 7.02E-07 | 31.39 | 3 | Bl vs Na: 4.738316e-06 |
| | | | | | Bl vs Tr: 5.481956e-08 | | | | Bl vs Tr: 2.292456e-05 |
| | | | | | Bl vs Ut: 5.849791e-04 | | | | Bl vs Ut: 2.257601e-02 |
| | | | | | Na vs Ut: 7.121111e-03 | | | | Na vs Ut: 2.478406e-02 |
| | | | | | Tr vs Ut: 2.596918e-02 | | | | Tr vs Ut: 3.610709e-02 |
| | 40 | 6.74E-11 | 50.35 | 3 | Bl vs Na: 5.691402e-09 | 3.56E-11 | 51.65 | 3 | Bl vs Na: 1.035833e-09 |
| | | | | | Bl vs Tr: 7.380146e-09 | | | | Bl vs Tr: 8.772967e-09 |
| | | | | | Bl vs Ut: 1.734934e-03 | | | | Bl vs Ut: 4.253376e-04 |
| | | | | | Na vs Ut: 4.969408e-03 | | | | Na vs Ut: 6.351867e-03 |

| | | | | | | | | |
|---|---|---|---|---|---|---|---|---|
| | | | | Tr vs Ut: 3.984141e-03 | | | | Tr vs Ut: 1.395540e-02 |
| 50 | 4.08E-06 | 27.76 | 3 | Bl vs Na: 2.911524e-06 | 2.00E-07 | 33.98 | 3 | Bl vs Na: 3.664008e-07 |
| | | | | Bl vs Tr: 3.884803e-04 | | | | Bl vs Tr: 5.972800e-06 |
| | | | | Bl vs Ut: 1.791795e-02 | | | | Bl vs Ut: 2.984771e-03 |
| | | | | Na vs Ut: 1.829421e-02 | | | | Na vs Ut: 3.006880e-02 |
| 60 | 0.00048 | 17.82 | 3 | Bl vs Na: 0.001010108 | 0.000321 | 18.66 | 3 | Bl vs Na: 0.0004884742 |
| | | | | Bl vs Tr: 0.001831081 | | | | Bl vs Tr: 0.0015469228 |
| 70 | | | | | 0.0242 | 9.42 | 3 | Na vs Ut: 0.01942243 |
