## Supplementary material for "Changes in the Dopaminergic circuitry and Adult Neurogenesis linked to Reinforcement Learning in Corvids": Table S17

**Table S17**. Statistical details of structural analysis of inactive DCX neurons in NCL subdivisions.

| Area | Parameters | Kruskal-Wallis P value | χ2 value | df | Post hoc P value (Dunn’s) |
|---|---|---|---|---|---|
| mNCL | Neurite length | 3.27E-05 | 20.659 | 2 | Na vs Ut: 0.0006<br>Tr vs Ut: 0.0002 |
| | Endings | 0.0027 | 11.787 | 2 | Na vs Ut: 0.0241<br>Tr vs Ut: 0.0048 |
| | Nodes | 0.0007 | 14.368 | 2 | Na vs Ut: 0.0051<br>Tr vs Ut: 0.0027 |
| | Neurite field area | 0.0002 | 16.475 | 2 | Na vs Ut: 0.0019<br>Tr vs Ut: 0.0014 |
| lNCL | Neurite length | 6.06E-06 | 24.029 | 2 | Na vs Ut: 1.8124e-05<br>Tr vs Ut: 8.0732e-04 |
| | Endings | 2.21E-06 | 26.042 | 2 | Na vs Ut: 5.4507e-06<br>Tr vs Ut: 7.2868e-04 |
| | Nodes | 1.30E-06 | 27.102 | 2 | Na vs Ut: 2.9842e-06<br>Tr vs Ut: 6.6800e-04 |
| | Neurite field area | 1.54E-06 | 26.766 | 2 | Na vs Ut: 4.1588e-06<br>Tr vs Ut: 2.7428e-04 |
