## Supplementary material for "Changes in the Dopaminergic circuitry and Adult Neurogenesis linked to Reinforcement Learning in Corvids": Table S18

**Table S18**. Statistical details of Sholl analysis of inactive DCX neurons in NCL subdivisions.

| **Area** | **Radius (μm)** | **Intersections (P-Value)** | **χ2 /F value** | **df** | **Intersections (Dunn's / Tukey's P value)** | **Neurite length (P-Value)** | **χ2 /F value** | **df** | **Neurite length (Dunn's / Tukey's P value)** |
|---|---|---|---|---|---|---|---|---|---|
| lNCL | 20 | 0.00461 | 5.84 | 2, 67 | Na vs Ut: 0.0102<br>Tr vs Ut: 0.0244 | - | | | - |
| | 30 | 1.52E-08 | 23.84 | 2, 67 | Na vs Ut: <1e-04<br>Tr vs Ut: <1e-04 | 2.55E-05 | 21.15 | 2 | Na vs Ut: 5.94e-05<br>Tr vs Ut: 2.079e-03 |
| | 40 | 1.79E-08 | 23.55 | 2, 67 | Na vs Ut: <1e-05<br>Tr vs Ut: <1e-05 | 5.15E-08 | 21.78 | 2, 67 | Na vs Ut: 1.14e-06<br>Tr vs Ut: 1.71e-06 |
| | 50 | 1.92E-05 | 21.73 | 2 | Na vs Ut: 5.9789e-05<br>Tr vs Ut: 9.8375e-04 | 1.32E-05 | 22.47 | 2 | Na vs Ut: 4.43e-05<br>Tr vs Ut: 7.56e-04 |
| | 60 | 0.0004 | 15.64 | 2 | Na vs Ut: 0.0003<br>Tr vs Ut: 0.0229 | 0.000 | 14.83 | 2 | Na vs Ut: 0.0008<br>Tr vs Ut: 0.0097 |
| | 70 | - | | | - | 0.0191 | 7.92 | 2 | Na vs Ut: 0.0167 |
| mNCL | 30 | 0.001846 | 12.59 | 2 | Na vs Ut: 0.0172<br>Tr vs Ut: 0.0036 | 0.01492 | 8.41 | 2 | Tr vs Ut: 0.0123 |
| | 40 | 4.11E-05 | 20.20 | 2 | Na vs Ut: 0.0008<br>Tr vs Ut: 0.0002 | 0.00261 | 6.51 | 2, 67 | Na vs Ut: 0.0397<br>Tr vs Ut: 0. 0.0034 |
| | 50 | 4.10E-05 | 20.20 | 2 | Na vs Ut: 0.0004<br>Tr vs Ut: 0.0003 | 3.59E-05 | 20.47 | 2 | Na vs Ut: 0.0003<br>Tr vs Ut: 0.0003 |
| | 60 | 0.0001324 | 17.86 | 2 | Na vs Ut: 0.0014<br>Tr vs Ut: 0.0005 | 0.001692 | 12.76 | 2 | Na vs Ut: 0.0057<br>Tr vs Ut: 0.0064 |
| | 70 | 0.00649 | 10.08 | 2 | Na vs Ut: 0.0077<br>Tr vs Ut: 0.0416 | 0.003973 | 11.06 | 2 | Na vs Ut: 0.0090<br>Tr vs Ut: 0.0121 |
