## Supplementary material for "Changes in the Dopaminergic circuitry and Adult Neurogenesis linked to Reinforcement Learning in Corvids": Table S19

**Table S19**. Statistical details of comparisons between active and inactive DCX neurons in NCL subdivisions.

| Area | Experimental group | Parameter | Test | n | df | t/W | P |
|---|---|---|---|---|---|---|---|
| lNCL | Undertrained | Soma area | Wilcoxon rank sum test | 30, 30 | | 211 | 2.90E-04 |
| | Trained | Neurite length | Wilcoxon rank sum test | 20, 20 | | 280.5 | 0.03042 |
| | No-Association | Neurite length | Wilcoxon rank sum test | 20, 20 | | 352 | 1.14E-05 |
| | No-Association | Endings | Welch's t-test | 20, 20 | 36.558 | 4.645 | 4.30E-05 |
| | No-Association | Nodes | Welch's t-test | 20, 20 | 34.863 | 4.8907 | 2.25E-05 |
| | No-Association | Neurite field area | Wilcoxon rank sum test | 20, 20 | | 337 | 0.000104 |
| | No-Association | Soma area | Welch's t-test | 20, 20 | 37.187 | -2.1684 | 0.03659 |
| mNCL | Undertrained | Soma area | Wilcoxon rank sum test | 30,30 | | 140 | 4.74E-06 |
| | Trained | Neurite length | Welch's t-test | 20, 20 | 35.141 | 3.4994 | 0.001287 |
| | Trained | neurite field area | Wilcoxon rank sum test | 20, 20 | | 350 | 1.57E-05 |
| | Trained | Soma area | Welch's t-test | 20, 20 | 37.939 | -3.0798 | 0.003841 |
| | No-Association | Neurite length | Welch's t-test | 20, 20 | 35.454 | 5.2081 | 8.26E-06 |
| | No-Association | Endings | Wilcoxon rank sum test | 20, 20 | | 284 | 0.02325 |
| | No-Association | Nodes | Wilcoxon rank sum test | 20, 20 | | 293 | 0.01194 |
| | No-Association | Neurite field area | Wilcoxon rank sum test | 20, 20 | | 372 | 2.64E-07 |
| | No-Association | Soma area | Welch's t-test | 20, 20 | 34.91 | -2.8849 | 0.006668 |
